# Inhibitory proteins block substrate access by occupying the active site cleft of *Bacillus subtilis* intramembrane protease SpoIVFB

**DOI:** 10.1101/2021.07.09.451828

**Authors:** Sandra Olenic, Lim Heo, Michael Feig, Lee Kroos

**Affiliations:** Department of Biochemistry and Molecular Biology, Michigan State University, East Lansing, MI 48824

## Abstract

Intramembrane proteases function in numerous signaling pathways that impact health, but knowledge about regulation of intramembrane proteolysis is limited. We examined inhibition of intramembrane metalloprotease SpoIVFB by proteins BofA and SpoIVFA. We found that BofA residues in and near a predicted transmembrane segment are required for SpoIVFB inhibition, and cross-linking experiments indicated that this transmembrane segment occupies the SpoIVFB active site cleft. SpoIVFA is also required for SpoIVFB inhibition. The inhibitory proteins block access of the substrate N-terminal Proregion to the membrane-embedded SpoIVFB active site, based on additional cross-linking experiments; however, the inhibitory proteins did not prevent interaction between the substrate C-terminal region and the SpoIVFB soluble domain. A structural model was built of SpoIVFB in complex with BofA and parts of SpoIVFA and substrate, using partial homology and constraints from cross-linking and co-evolutionary analyses. The model predicts that conserved BofA residues interact to stabilize a transmembrane segment and a membrane-embedded C-terminal region. SpoIVFA is predicted to bridge the BofA C-terminal region and SpoIVFB, forming a membrane-embedded inhibition complex. Our results reveal a novel mechanism of intramembrane protease inhibition with clear implications for relief from inhibition *in vivo* and design of inhibitors as potential therapeutics.

## Introduction

Intramembrane proteases (IPs) are membrane proteins containing a membrane-embedded active site. IPs cleave membrane-associated substrates within a transmembrane segment (TMS) or near the membrane surface in a process referred to as regulated intramembrane proteolysis (RIP) (1). Released substrate fragments impact diverse signaling pathways in a wide variety of organisms (2). IPs also profoundly impact protein degradation (3). There are four known IP families: metallo IPs like SpoIVFB, aspartyl IPs like γ-secretase, serine IPs (rhomboids), and the glutamyl IP Rce1 (2, 4). Crystal structures have been solved for one or more IP in each family (4–9), revealing that TMSs arrange to form a channel that delivers water to the active site for hydrolysis of a substrate peptide bond. Structures have also been solved for rhomboid·peptide (inhibitor and substrate) complexes (10–12) and for γ-secretase·substrate complexes (13, 14), which may guide the design of IP modulators as therapeutics.

Metallo IPs activate transcription factors via RIP in all three domains of life (1, 2). In mammals, S2P is involved in regulation of cholesterol homeostasis and responses to endoplasmic reticulum stress and viral infection (15, 16). S2P homologs in plants and in the pathogenic fungus *Cryptococcus neoformans* play important roles in chloroplast development and virulence, respectively (17, 18). In bacteria, metallo IPs enhance pathogenicity, control stress responses and polar morphogenesis, produce mating signals, and clear signal peptides from the membrane (19–21). In bacilli and most clostridia that form spores, SpoIVFB cleaves inactive Pro-σ^K^ to σ^K^, which directs RNA polymerase to transcribe genes necessary for endospore formation (22–25). Endospores are dormant and are able to survive harsh environmental conditions (26), enhancing the persistence of pathogenic species (27–29). Understanding regulation of SpoIVFB could lead to new strategies to manipulate endosporulation and other processes involving IPs in bacteria and eukaryotes.

During endospore formation, *B. subtilis* forms a polar septum that divides the mother cell (MC) and forespore (FS) compartments (30) (Figure 1-figure supplement 1). The MC engulfs the FS, surrounding it with a second membrane and pinching it off within the MC. Regulation of SpoIVFB involves inhibition by BofA and SpoIVFA (31–33). The three proteins form a complex in the outer FS membrane during engulfment (34–36). BofA was implicated as the direct inhibitor of SpoIVFB (37), while SpoIVFA appeared to localize and stabilize the inhibition complex (36, 38). Signaling from the FS relieves inhibition of SpoIVFB (Figure 1- figure supplement 1). Two proteases, SpoIVB and CtpB, are exported into the space between the two membranes surrounding the FS (39, 40). SpoIVB cleaves the C-terminal end of SpoIVFA (41–44) and CtpB can cleave the C-terminal ends of both SpoIVFA and BofA (42, 43, 45). Once inhibition is removed, SpoIVFB cleaves the N-terminal 21-residue Proregion from Pro-σ^K^, releasing σ^K^ into the MC (46–48). σ^K^ directs RNA polymerase to transcribe genes whose products form the spore coat and lyse the MC, releasing a mature spore (49, 50).

**Figure 1.**
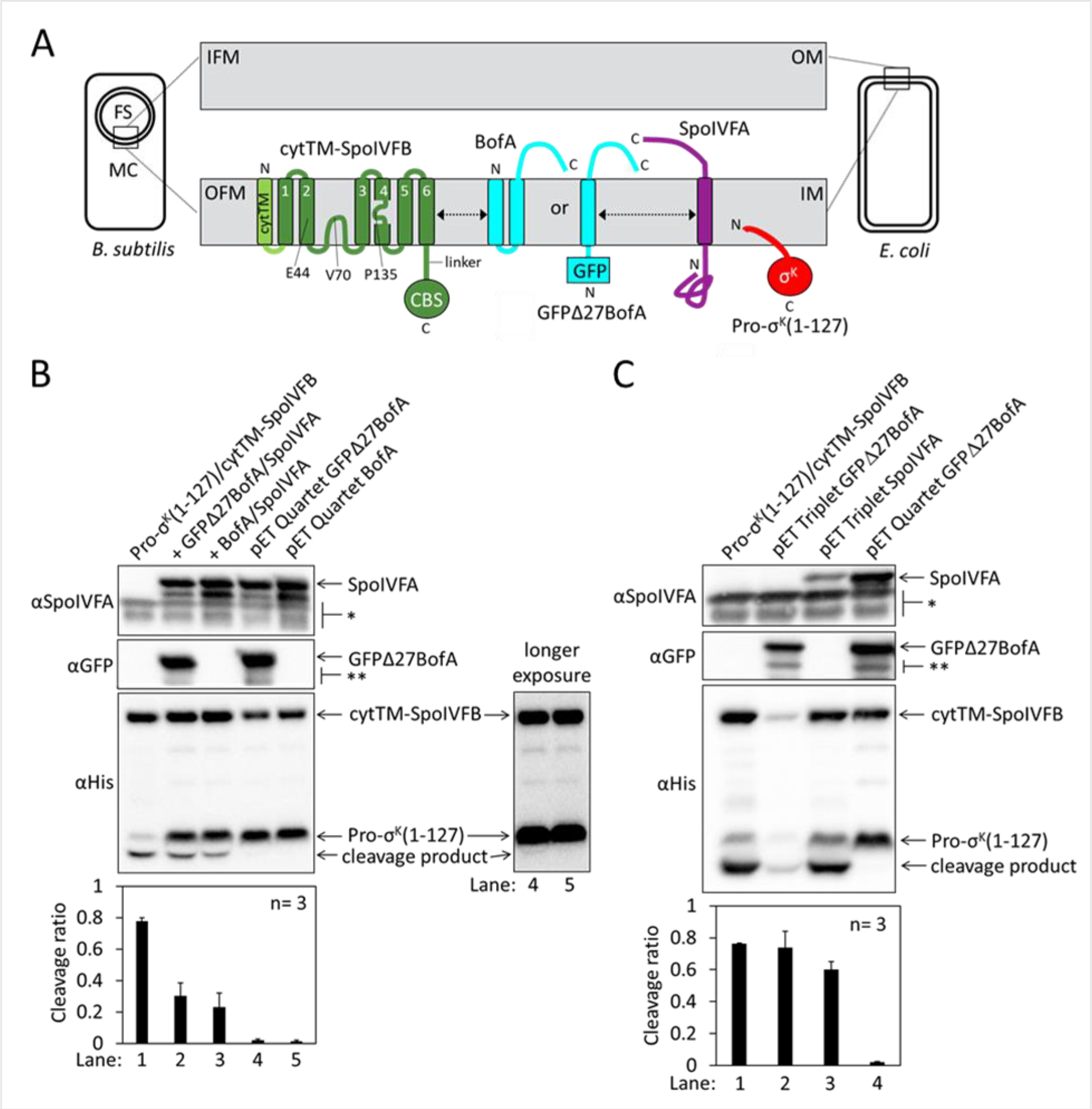
with 4 supplements Inhibition of Pro-σ^K^ cleavage. (**A**) Diagram of SpoIVFB inhibition during *B. subtilis* endosporulation and upon heterologous expression in *E. coli*. During endosporulation (*Left*), SpoIVFB and its inhibitory proteins BofA and SpoIVFA are produced in the mother cell (MC) and localize to the outer forespore (FS) membrane (OFM). Pro-σ^K^ is also produced in the MC and associates with membranes. When these proteins are synthesized in *E. coli* (*Right*), they localize to the inner membrane (IM). The expanded view of the membranes (*Center*) shows a SpoIVFB variant with an extra N-terminal transmembrane segment (cytTM), and highlights several residues (E44, V70, P135) at or near the active site in the membrane domain, which is connected to the CBS domain by an interdomain linker. When produced in *E. coli*, cytTM-SpoIVFB cleaves Pro-σ^K^(1-127), removing its N- terminal Proregion [Note: Pro-σ^K^(1-126) was renamed Pro-σ^K^(1-127) as explained in (56)]. Co- production of SpoIVFA and either full-length BofA or GFPΔ27BofA (lacking predicted TMS1) inhibits Pro-σ^K^(1-127) cleavage. The dashed double-headed arrows indicate that SpoIVFB, BofA or GFPΔ27BofA, and SpoIVFA form a complex of unknown structure. (**B**) Cleavage assays comparing inhibition by SpoIVFA and either GFPΔ27BofA or full-length BofA in *E. coli*. Pro-σ^K^(1-127) and cytTM-SpoIVFB were produced alone (lane 1, pYZ2) or in combination with GFPΔ27BofA and SpoIVFA (lane 2, pYZ46) or full-length BofA and SpoIVFA (lane 3, pSO212). Alternatively, “pET Quartet” plasmids were used to produce Pro-σ^K^(1-127), cytTM- SpoIVFB, SpoIVFA, and either GFPΔ27BofA (lane 4, pSO40) or full-length BofA (lane 5, pSO213). Samples collected after 2 h of IPTG induction were subjected to immunoblot analysis with SpoIVFA (*Top*), GFP (*Middle*), or penta-His antibodies (*Bottom*, 2 and 30 s exposures). The single star (*) indicates cross-reacting proteins below SpoIVFA and the double star (**) indicates breakdown species of GFPΔ27BofA. A breakdown species below SpoIVFA (not indicated) is observed in some experiments. The graph shows quantification of the cleavage ratio (cleavage product/[Pro-σ^K^(1-127) + cleavage product]) for three biological replicates. Error bars, 1 standard deviation. (**C**) Cleavage assays comparing inhibition by either GFPΔ27BofA or SpoIVFA. pET Triplet plasmids were used to produce Pro-σ^K^(1-127), cytTM-SpoIVFB, and either GFPΔ27BofA (lane 2, pSO64) or SpoIVFA (lane 3, pSO65). Samples were subjected to immunoblot analysis and quantification as in (**B**). Figure 1-source data 1 Immunoblot images (raw and annotated) and quantification of cleavage assays (Figure 1 B and C).

Regulation of SpoIVFB by direct interaction with the inhibitory proteins BofA and SpoIVFA differs from known regulation of other IPs. Rather, in eukaryotic cells some IP substrates initially localize to a different organelle than their cognate IP, and regulation involves membrane trafficking to deliver the substrate to the enzyme (3, 51). In bacteria as well as eukaryotes, many IP substrates require an initial extramembrane cleavage by another protease prior to the intramembrane cleavage, and the initial cleavage is typically the regulated step (19-21, 52, 53). In contrast, Pro-σ^K^ is cleaved only once, by SpoIVFB. Cleavage of SpoIVFA and BofA by SpoIVB and CtpB is regulated (Figure 1-figure supplement 1), but how this relieves inhibition of SpoIVFB has been unknown. Understanding the mechanism of inhibition could have broad implications since the catalytic core of metallo IPs includes three conserved TMSs (54) and general principles may emerge that inform potential inhibition strategies for other IP types.

Cleavage of Pro-σ^K^ by SpoIVFB can be reproduced by expressing these proteins in *Escherichia coli* (37) (Figure 1A). Cleavage could be inhibited by additionally producing GFPΔ27BofA (a functional fusion protein lacking predicted TMS1 of BofA) and SpoIVFA (37, 45). However, we found that the extent of cleavage inhibition was incomplete and variable.

Here, we describe an improved cleavage inhibition assay. Using the assay and bioinformatics, we identified three conserved residues of BofA important for inhibition of SpoIVFB in *E. coli* and in sporulating *B. subtilis.* One of the residues is near the middle of predicted TMS2 of BofA, close to the sole Cys residue of BofA, C46. We exploited C46 for disulfide cross-linking experiments that showed BofA TMS2 can occupy the SpoIVFB active site cleft. Additional experiments in *E. coli* indicate that BofA and SpoIVFA block access of the N-terminal Proregion of Pro-σ^K^ to the membrane-embedded SpoIVFB active site, yet the C-terminal soluble regions of Pro-σ^K^ and SpoIVFB can interact. These results reveal a novel mechanism of IP inhibition and were used in combination with prior cross-linking studies (55, 56), partial homology, and evolutionary co-variation of amino acid residues, to constrain a structural model of SpoIVFB in complex with BofA and parts of SpoIVFA and Pro-σ^K^. The model predicts that conserved residues in TMS2 and the C-terminal region of BofA interact, and that SpoIVFA bridges the BofA C-terminal region and SpoIVFB to form an inhibition complex. The predicted inhibition complex has implications for activation of SpoIVFB and its orthologs in other endospore formers, as well as for translational efforts to design therapeutic IP inhibitors.

## Results

### Both BofA and SpoIVFA Are Required to Inhibit SpoIVFB in *E. coli*

To study RIP of Pro- σ^K^, *E*. *coli* was engineered to synthesize variants of SpoIVFB and Pro-σ^K^, and the inhibitory proteins BofA and SpoIVFA, in various combinations. The SpoIVFB variant contains the extra TMS cytTM (57) (Figure 1A), which improves accumulation (48). The substrate variant Pro- σ^K^(1–127) lacks the C-terminal half of Pro-σ^K^, but is accurately cleaved by SpoIVFB and the cleavage product is easily separated from Pro-σ^K^(1–127) by SDS-PAGE (58). When Pro-σ^K^(1–127) and cytTM-SpoIVFB were produced from the same plasmid, about 80% of the substrate was cleaved (Figure 1B). The additional production of GFPΔ27BofA and SpoIVFA from a second plasmid reduced cleavage to about 30%. A similar result was seen when full-length BofA (for which no antibody has been described) and SpoIVFA were produced from a second plasmid.

Since the two plasmids had the same replication origin, their copy number may be unequal in some cells, despite using different selection markers. Unequal copy numbers could result in Pro-σ^K^(1–127) cleavage in cells producing too little of the inhibitory proteins.

Therefore, we designed single “pET Quartet” plasmids to synthesize all four proteins. When pET Quartets produced either GFPΔ27BofA or BofA, very little Pro-σ^K^(1–127) was cleaved (Figure 1B). A longer exposure of the immunoblot revealed a faint cleavage product with pET Quartet GFPΔ27BofA, but not with pET Quartet BofA, indicating that full-length BofA inhibits cleavage slightly better than GFPΔ27BofA.

We tested whether both inhibitory proteins are required for cleavage inhibition by engineering “pET Triplet” plasmids to synthesize Pro-σ^K^(1–127), cytTM-SpoIVFB, and either GFPΔ27BofA, full-length BofA, or SpoIVFA. With pET Triplets containing GFPΔ27BofA (Figure 1C) or BofA (Figure 1-figure supplement 2A), cleavage inhibition was lost and accumulation of cytTM-SpoIVFB, Pro-σ^K^(1–127), and cleavage product was greatly diminished, suggesting that BofA inhibits synthesis and/or enhances degradation of the other proteins when SpoIVFA is absent. With pET Triplet SpoIVFA, cleavage inhibition was also nearly lost (Figure 1C). We conclude that both inhibitory proteins are necessary to prevent Pro-σ^K^(1–127) cleavage by cytTM-SpoIVB in *E. coli*. Both inhibitory proteins are also required to prevent cleavage of full-length Pro-σ^K^ in *E. coli* (Figure 1-figure supplement 2B). The heterologous system mimics the endogenous pathway of *B. subtilis*, since both inhibitory proteins are necessary to prevent cleavage of Pro-σ^K^ by SpoIVFB during sporulation (31, 36, 59).

We also showed that the *E. coli* system mimics the endogenous pathway with respect to partial relief from inhibition by SpoIVB (40, 42, 43, 45) (Figure 1-figure supplement 3) or an F66A substitution in SpoIVFB (59) (Figure 1-figure supplement 4). We reasoned that the strong cleavage inhibition observed with pET Quartet plasmids in *E. coli* (Figure 1B) could be used to test the effects of Ala substitutions in the inhibitory proteins. Since most endospore formers encode BofA, but about half lack a recognizable gene for SpoIVFA (22, 23), we focused first on identifying residues of BofA important for inhibition of SpoIVFB.

### Three Conserved Residues of BofA Are Important for Inhibition of SpoIVFB in *E. coli*

To identify residues of BofA that may play a role in SpoIVFB inhibition, an alignment of 70 BofA orthologs was analyzed (Figure 2-figure supplement 1A). Thirteen highly conserved residues reside in predicted TMS2 and the C-terminal region. The lack of highly conserved residues in TMS1 may explain why GFPΔ27BofA (lacking predicted TMS1) is a functional inhibitor (36, 37) (Figure 1B). An alignment of just the 31 BofA orthologs from species with a recognizable gene for SpoIVFA revealed an additional four conserved residues that we deemed of interest for Ala substitutions (Figure 2-figure supplement 1B). We also substituted Ala for F85 and I87 (as well as for I86, which is conserved), since deletion of three residues from the C-terminal end of BofA caused a loss of function or stability in *B. subtilis* (33, 60).

**Figure 2.**
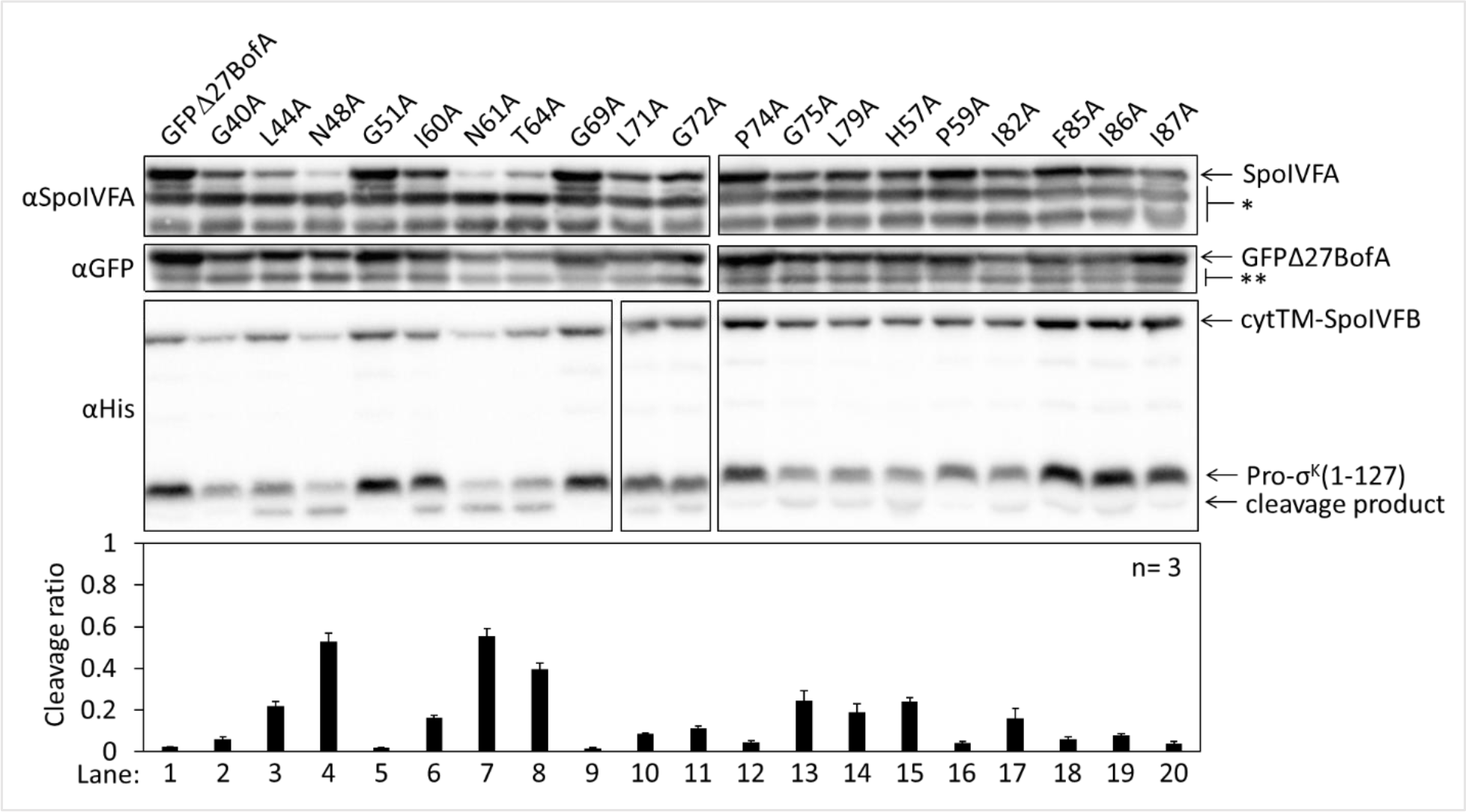
with 2 supplements Effects of alanine substitutions in GFPΔ27BofA on inhibition of Pro-σ^K^(1-127) cleavage in *E. coli*. pET Quartet plasmids were used to produce Pro-σ^K^(1-127), cytTM-SpoIVFB, SpoIVFA, and GFPΔ27BofA (lane 1, pSO40) or Ala-substituted GFPΔ27BofA (lanes 2-20, pSO44-pSO58 and pSO60-pSO63). Samples collected after 2 h of IPTG induction were subjected to immunoblot analysis with SpoIVFA, GFP, and penta-His antibodies. Single (*) and double (**) stars are explained in the Figure 1B legend, as is the graph. Figure 2-source data 1 Immunoblot images (raw and annotated) and quantification of cleavage assays.

The Ala substitutions were made in GFPΔ27BofA and co-produced with Pro-σ^K^(1–127), cytTM-SpoIVFB, and SpoIVFA from pET Quartet plasmids in *E. coli*. Three GFPΔ27BofA variants, N48A, N61A, and T64A led to 40-60% cleavage of Pro-σ^K^(1–127), indicating that these substitutions strongly impaired inhibition of cytTM-SpoIVFB (Figure 2). Less of the N61A and T64A variants was observed than most of the other GFPΔ27BofA variants, and for all three variants, less SpoIVFA and cytTM-SpoIVFB was observed, perhaps indicative of reduced complex formation leading to protein instability, as inferred from studies in *B. subtilis* (36, 61). We also examined the effects of the GFPΔ27BofA N48A, N61A, and T64A variants during *B. subtilis* sporulation (see the next section).

The other GFPΔ27BofA variants had less effect on Pro-σ^K^(1–127) cleavage, indicating less effect on cytTM-SpoIVFB inhibition in *E. coli* (Figure 2). Although the F85A, I86A, and I87A variants had little effect, both a triple-Ala variant and a variant lacking residues 85-87 strongly impaired inhibition of cytTM-SpoIVFB (Figure 2-figure supplement 2A); the latter mimicking the results in *B. subtilis* (33, 60). We also used pET Quartet plasmids in *E. coli* to show for the first time that the nine residues preceding predicted TMS2 of GFPΔ27BofA contribute to its inhibitory function, perhaps by moving the GFP tag away from the membrane (Figure 2-figure supplement 2B).

### The Conserved Residues of BofA Are Important for SpoIVFB Inhibition during *B. subtilis* Sporulation

To test the effects of the three GFPΔ27BofA variants (N48A, N61A, and T64A) during sporulation, each was produced ectopically under control of the MC-specific *bofA* promoter in a *spoIVB165 bofA::erm*double mutant. The *spoIVB165* null mutation alone would block Pro-σ^K^ cleavage, but the *bofA::erm* null mutation allows unregulated cleavage in the double mutant, providing a genetic background in which ectopic production of GFPΔ27BofA under control of a MC-specific promoter leads to inhibition of SpoIVFB and loss of Pro-σ^K^ cleavage (36, 37). As anticipated, Pro-σ^K^ was cleaved primarily between 4 and 5 h poststarvation (PS) in wild type, very little cleavage was observed in the *spoIVB165* single mutant, and cleavage occurred prematurely at 4 h in the *spoIVB165 bofA::erm* double mutant (Figure 3A). Very little SpoIVFA and SpoIVFB was detected in the double mutant, consistent with the need for BofA to stabilize these proteins (36). Also as expected, ectopic production of GFPΔ27BofA in the double mutant inhibited SpoIVFB so that little Pro-σ^K^ cleavage was observed (lanes 7 & 8). GFPΔ27BofA allowed slightly more Pro-σ^K^ cleavage at 5 h (lane 8) than BofA in the *spoIVB165* single mutant (lane 4), suggesting that full-length BofA is a slightly better inhibitor of SpoIVFB during *B. subtilis* sporulation, as observed in *E. coli* (Figure 1B, longer exposure).

**Figure 3.**
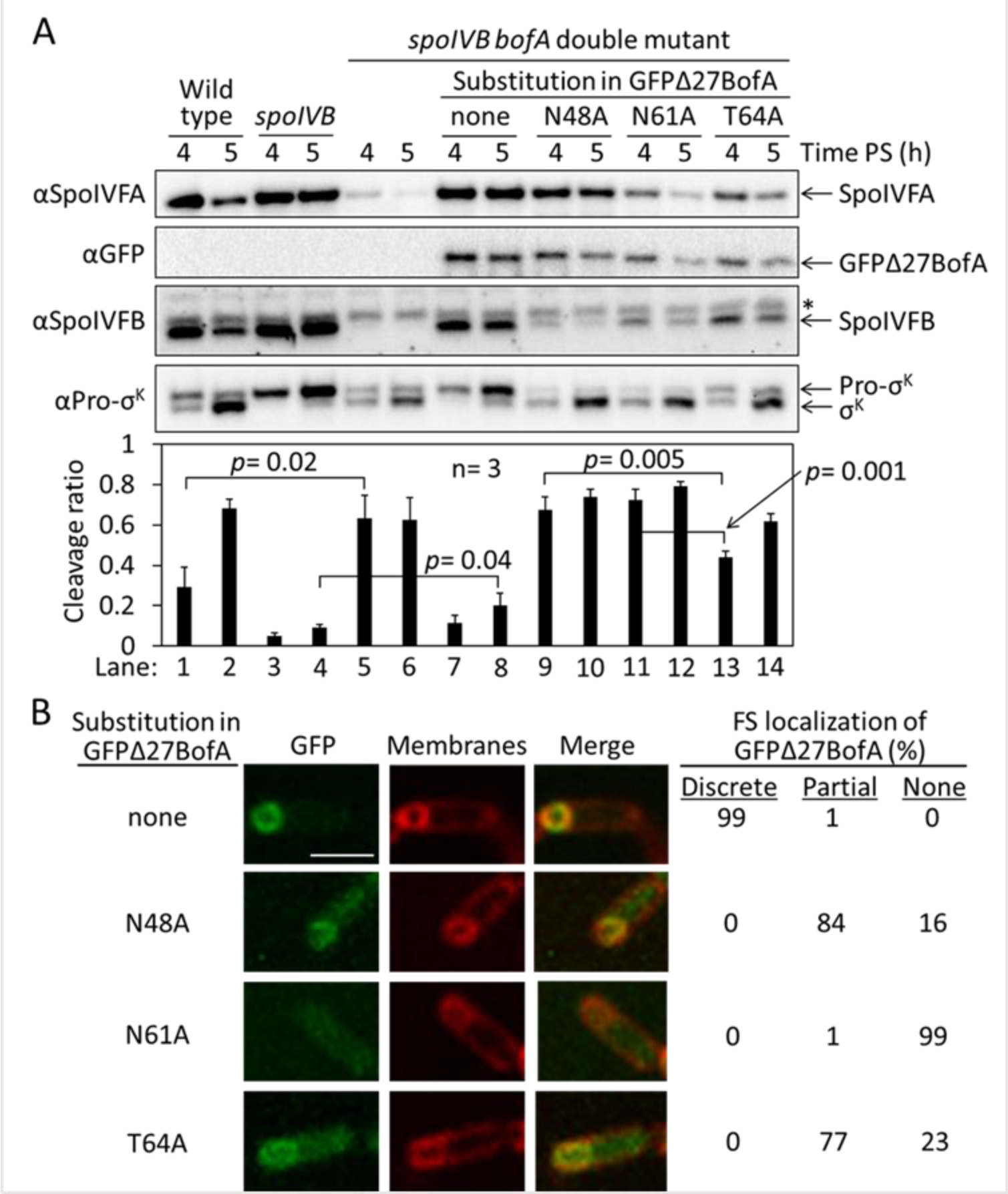
with 1 supplement Effects of alanine substitutions in GFPΔ27BofA during *B. subtilis* sporulation. (**A**) Effects of GFPΔ27BofA variants (N48A, N61A, and T64A) on Pro-σ^K^ cleavage. Wild-type strain PY79, a *spoIVB165* null mutant, a *spoIVB165 bofA::erm* double mutant, and the double mutant with P*bofA*-*gfpΔ27bofA* integrated ectopically at *amyE* to express GFPΔ27BofA with no substitution (none) or the indicated Ala substitution, were starved to induce sporulation. Samples collected at 4 and 5 h poststarvation (PS) were subjected to immunoblot analysis with antibodies against SpoIVFA, GFP, SpoIVFB, and Pro-σ^K^. The graph shows quantification of the cleavage ratio [σ^K^/(Pro-σ^K^ + σ^K^)] for three biological replicates. Error bars, 1 standard deviation. Student’s two-tailed *t* tests were performed to compare certain cleavage ratios (*p* values). (**B**) Localization of GFPΔ27BofA and the three variants. Samples collected at 3 h PS were treated with FM 4-64 to stain membranes. Confocal microscopy images of fluorescence from GFPΔ27BofA, membranes, and merged images are shown for representative sporangia with discrete (no substitution in GFPΔ27BofA, designated “none”), partial (N48A and T64A), or no forespore (FS) localization (N61A). Scale bar, 1 μm. The percentage of sporangia (44-93 counted; non-sporulating cells were not counted) with each localization pattern is shown. Figure 3-source data 1 Immunoblot images (annotated) and quantification of cleavage assays (Figure 3A), and confocal microscopy images (see the readme file) and a table of the numbers of sporangia in which FS localization of GFPΔ27BofA and the three variants was counted (Figure 3B). Figure 3-source data 2 Immunoblot images (raw) (Figure 3A).

Strikingly, ectopic production of the GFPΔ27BofA N48A or N61A variant in the double mutant did not inhibit SpoIVFB, as Pro-σ^K^ was cleaved prematurely at 4 h (Figure 3A, lanes 9 & 11). The level of the N48A variant was comparable with GFPΔ27BofA, but less of the N61A variant was observed, especially at 5 h (lane 12). In comparison with the strain that produced GFPΔ27BofA, the SpoIVFA level was normal in the strain that produced the N48A variant, but less SpoIVFB was observed, and less of both SpoIVFA and SpoIVFB was observed in the strain producing the N61A variant. The reduced levels of SpoIVFB and SpoIVFA may reflect protein instability due to less complex formation with the GFPΔ27BofA variants, as inferred from reduced SpoIVFB and SpoIVFA levels in the absence of BofA during *B. subtilis* sporulation (36).

Production of the T64A variant in the double mutant (Figure 3A, lanes 13 & 14) showed a pattern similar to wild type; Pro-σ^K^ cleavage increased between 4 and 5 h, and the levels of SpoIVFA and SpoIVFB decreased. The levels of all three proteins were slightly lower in the strain producing the T64A variant than in wild type. The level of the T64A variant was similar to the N48A and N61A variants at 4 h, yet a lower percentage of the Pro-σ^K^ was cleaved in the T64A variant (lanes 9, 11 & 13), suggesting that the T64A variant inhibits SpoIVFB, although not as well as GFPΔ27BofA (lane 7). We conclude that the three conserved residues of BofA are important for SpoIVFB inhibition during sporulation.

Since GFPΔ27BofA co-localizes with SpoIVFA and SpoIVFB to the outer FS membrane during sporulation (34–36), we examined the ability of the GFPΔ27BofA variants to localize to the FS. As a control, GFPΔ27BofA produced in the *spoIVB165 bofA::erm* double mutant localized discretely to the FS at 3 h PS (Figure 3B). The N48A and T64A variants localized partially to the FS, but GFP fluorescence was also observed in the MC cytoplasm, suggesting partial mislocalization. The N61A variant appeared to be dispersed evenly throughout the MC cytoplasm. The fusion proteins were intact, demonstrating that the cytoplasmic GFP fluorescence was not attributable to breakdown (Figure 3-figure supplement 1). Inability of the N61A variant to localize to the FS may explain the loss of SpoIVFB inhibition and the abundant cleavage of Pro-σ^K^ at 4 h (Figure 3A, lane 11). Importantly, the similar ability of the N48A and T64A variants to localize to the FS (Figure 3B) does not account for their differential effects on the level of SpoIVFB and its ability to cleave Pro-σ^K^ (Figure 3A). The strain producing GFPΔ27BofA N48A exhibited less SpoIVFB yet more Pro-σ^K^ cleavage at 4 h (lane 9) than the strain producing GFPΔ27BofA T64A (lane 13), so the N48A substitution more severely impairs the ability of GFPΔ27BofA to inhibit SpoIVFB.

### BofA TMS2 Occupies the SpoIVFB Active Site Cleft

A common mechanism of metalloprotease inhibition involves a protein residue side chain coordinating the catalytic metal ion in place of a water molecule necessary for substrate peptide bond hydrolysis (62, 63). Early work suggested that BofA H57 might inhibit SpoIVFB by coordinating zinc (37). Although His is highly conserved at the corresponding position of BofA orthologs (Figure 2-figure supplement 1) and GFPΔ27BofA H57A allowed about 20% cleavage of Pro-σ^K^(1–127) in *E. coli* (Figure 2), H57 is near the predicted C-terminal end of BofA TMS2 (Figure 2-figure supplement 1) and therefore presumably near the membrane surface, whereas a zinc ion at the SpoIVFB active site is predicted to be near the middle of the lipid bilayer (56). The sole Cys of *B. subtilis* BofA, C46, is a potential zinc ligand located near the center of TMS2, but Cys is not conserved at the corresponding position in BofA orthologs (Figure 2-figure supplement 1). However, nearby N48 is highly conserved among BofA orthologs and is also a potential zinc ligand. For example, the latent form of *Staphylococcus aureus* LytM has a prosegment with an Asn residue as a zinc ligand (64). We hypothesized that BofA N48 is a zinc ligand and that BofA TMS2 occupies the SpoIVFB active site cleft in the inhibition complex.

To begin testing our hypothesis, we devised a strategy based on a model of the SpoIVFB membrane domain derived from the crystal structure of an archaeal homolog (7). The model is supported by cross-linking studies of catalytically-inactive SpoIVFB E44Q in complex with Pro-σ^K^(1–127), whose Proregion appears to occupy the SpoIVFB active site cleft (55, 56, 65). A cleft between TMS1 and TMS6 of SpoIVFB is proposed to gate substrate access to the active site (59), which mutational evidence supports is formed by a zinc ion near E44 of the HELGH metalloprotease motif within SpoIVFB TMS2 (46, 47) (Figure 4-figure supplement 1A). If BofA N48 is a zinc ligand and BofA TMS2 occupies the SpoIVFB active site cleft in the inhibition complex, we reasoned that BofA C46 may be near SpoIVFB E44 (Figure 4-figure supplement 1B) and we used a disulfide cross-linking approach to test this possibility.

**Figure 4.**
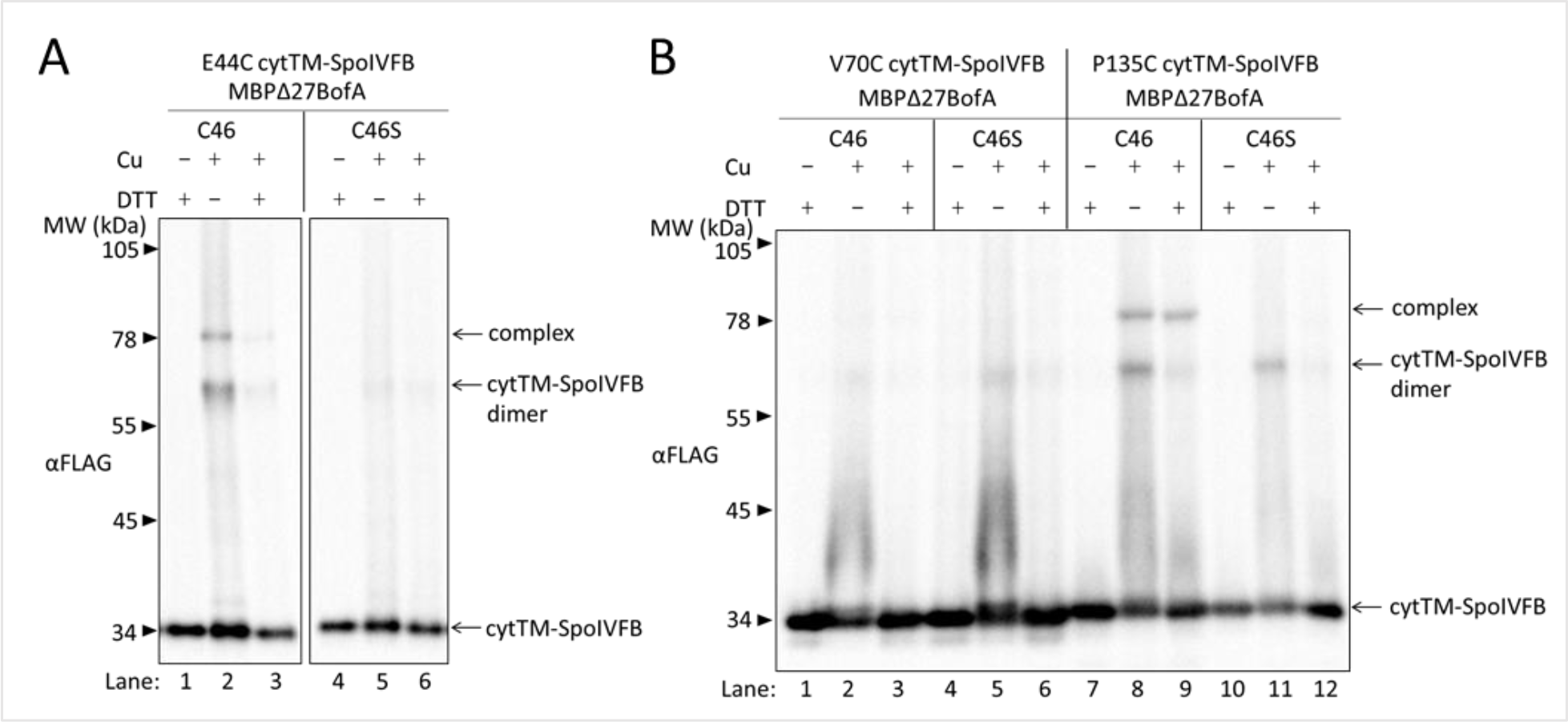
with 5 supplements BofA TMS2 is near the SpoIVFB active site. (**A**) Disulfide cross-linking of E44C at the cytTM-SpoIVFB active site to C46 in TMS2 of MBPΔ27BofA. pET Quartet plasmids were used to produce single-Cys E44C cytTM-SpoIVFB in combination with MBPΔ27BofA C46 (pSO91) or Cys-less MBPΔ27BofA C46S as a negative control (pSO110), and Cys-less variants of SpoIVFA and Pro-σ^K^(1-127) in *E. coli*. Samples collected after 2 h of IPTG induction were treated for 60 min with Cu^2+^(phenanthroline)3 (Cu +) to promote disulfide bond formation or with 2-phenanthroline (Cu –) as a negative control, then treated with TCA to precipitate proteins and resuspended in sample buffer with DTT (+) to reverse cross-links or without (–) to preserve cross-links, and finally subjected to immunoblot analysis with FLAG antibodies to visualize cytTM-SpoIVFB monomer, dimer, and complex with MBPΔ27BofA. (**B**) Disulfide cross-linking of V70C or P135C near the cytTM-SpoIVFB active site to C46 in TMS2 MBPΔ27BofA. pET Quartet plasmids were used to produce single- Cys V70C or P135C cytTM-SpoIVFB E44Q variants in combination with MBPΔ27BofA (pSO92, pSO93) or Cys-less MBPΔ27BofA C46S as a negative control (pSO111, pSO112), and Cys-less variants of SpoIVFA and Pro-σ^K^(1-127) in *E. coli*. Samples collected after 2 h of IPTG induction were treated and subjected to immunoblot analysis as in (**A**). A representative result from at least two biological replicates is shown in (**A**) and (**B**). Figure 4-source data 1 Immunoblot images (raw and annotated).

We tested whether BofA C46 could be cross-linked to SpoIVFB E44C in the inhibition complex formed with pET Quartet plasmids in *E. coli* as in Figure 1B, but with appropriate variants of the four proteins. Single-Cys E44C cytTM-SpoIVFB (which is catalytically-inactive) and Cys-less Pro-σ^K^(1–127) (which can be cleaved by active SpoIVFB) were described previously (55). We created functional variants of Cys-less SpoIVFA and single-Cys MBPΔ27BofA (GFP has Cys residues but MBP lacks them, leaving BofA C46 as the sole Cys residue) (Figure 4-figure supplement 2). As a negative control, we replaced C46 with Ser to obtain functional Cys-less MBPΔ27BofA. *E. coli* induced to co-produce combinations of four proteins were treated with the oxidant Cu^2+^(phenanthroline)3 to promote disulfide bond formation. For MBPΔ27BofA C46 (Figure 4A, lane 2), but not the Cys-less C46S negative control (lane 5), treatment with oxidant caused formation of a species of the expected size for a cross-linked complex with single-Cys E44C cytTM-SpoIVFB, which was detected by immunoblotting with anti-FLAG antibodies. Treatment with the reducing agent DTT greatly diminished the abundance of the apparent complex, consistent with cross-link reversal (lane 3). A species of the expected size for a cross-linked dimer of single-Cys E44C cytTM-SpoIVFB was also observed. Formation of the apparent dimer varies, as reported previously (55). As expected, anti-MBP antibodies detected the presumptive cross-linked complex of MBPΔ27BofA C46 with single-Cys E44C cytTM-SpoIVFB, albeit weakly, and the negative control with E44Q rather than E44C failed to form the complex (Figure 4-figure supplement 3, lanes 2 & 5). Since the signal for the complex was stronger with anti-FLAG antibodies (Figure 4A), we used those antibodies in the cross-linking experiments reported below.

In addition to E44 of SpoIVFB, V70 in a predicted membrane-reentrant loop and P135 in a predicted short loop interrupting TMS4 (Figure 1A and Figure 4-figure supplement 1A) were shown to be in proximity to the Proregion of Pro-σ^K^(1–127) (55). Therefore, we tested whether MBPΔ27BofA C46 could be cross-linked to single-Cys V70C or P135C cytTM-SpoIVFB E44Q variants. We included the inactivating E44Q substitution since the V70C and P135C variants (unlike the E44C variant) could cleave Cys-less Pro-σ^K^(1–127) (55), even though the inhibitory proteins were expected to almost completely inhibit cleavage (Figure 4-figure supplement 2).

MBPΔ27BofA C46 formed a complex with the P135C variant, but not with the V70C variant (Figure 4B, lanes 2 & 8). As expected, Cys-less MBPΔ27BofA C46S failed to form a complex with either variant (lanes 5 & 11). Full-length BofA C46 (lacking MBP) also formed a complex of the expected (smaller) size with the E44C and P135C variants, but not with the V70C variant (Figure 4-figure supplement 4, lanes 2, 14, & 8). As expected, Cys-less BofA C46S failed to form a complex with any of the variants (lanes 5, 11, & 17). Our cross-linking results show that BofA TMS2 occupies the SpoIVFB active site cleft in the inhibition complex, placing BofA C46 in proximity to SpoIVFB E44 and P135, but not V70.

Based on our initial cross-linking results, we modeled BofA TMS2 in the SpoIVFB active site cleft, and tested predictions of the model using additional disulfide cross-linking experiments (Figure 4-figure supplement 5). The results confirmed predictions of the initial model and suggested a preferred orientation of BofA TMS2 in the SpoIVFB active site cleft, which led to the refined model shown in Figure 4-figure supplement 1B.

### BofA and SpoIVFA Do Not Prevent Pro-σ^K^(1-127) from Interacting with SpoIVFB

Since our cross-linking results show that BofA TMS2 occupies the SpoIVFB active site cleft, we tested whether BofA and SpoIVFA prevent Pro-σ^K^(1–127) from interacting with SpoIVFB in *E. coli*. A catalytically-inactive E44C cytTM-SpoIVFB variant with a FLAG2 epitope tag was co-produced with Pro-σ^K^(1–127), SpoIVFA, and GFPΔ27BofA. Cell lysates were prepared and detergent-solubilized proteins were co-immunoprecipitated in pull-down assays with anti-FLAG antibody beads. All four proteins were seen in the bound sample (Figure 5A, lane 4), indicating that GFPΔ27BofA and SpoIVFA did not completely prevent Pro-σ^K^(1–127) from interacting with the cytTM-SpoIVFB variant. However, only the cytTM-SpoIVFB variant was detected when the bound sample was diluted tenfold (lane 3) to match the input sample concentration (lane 1), and portions of the other three proteins were observed in the unbound sample (lane 2), indicating that co-purification was inefficient. A negative control with the cytTM-SpoIVFB variant lacking the FLAG2 epitope tag showed no Pro-σ^K^(1–127) in the bound sample, but small amounts of GFPΔ27BofA and SpoIVFA were detected (lane 8), indicative of weak, nonspecific binding to the beads.

**Figure 5.**
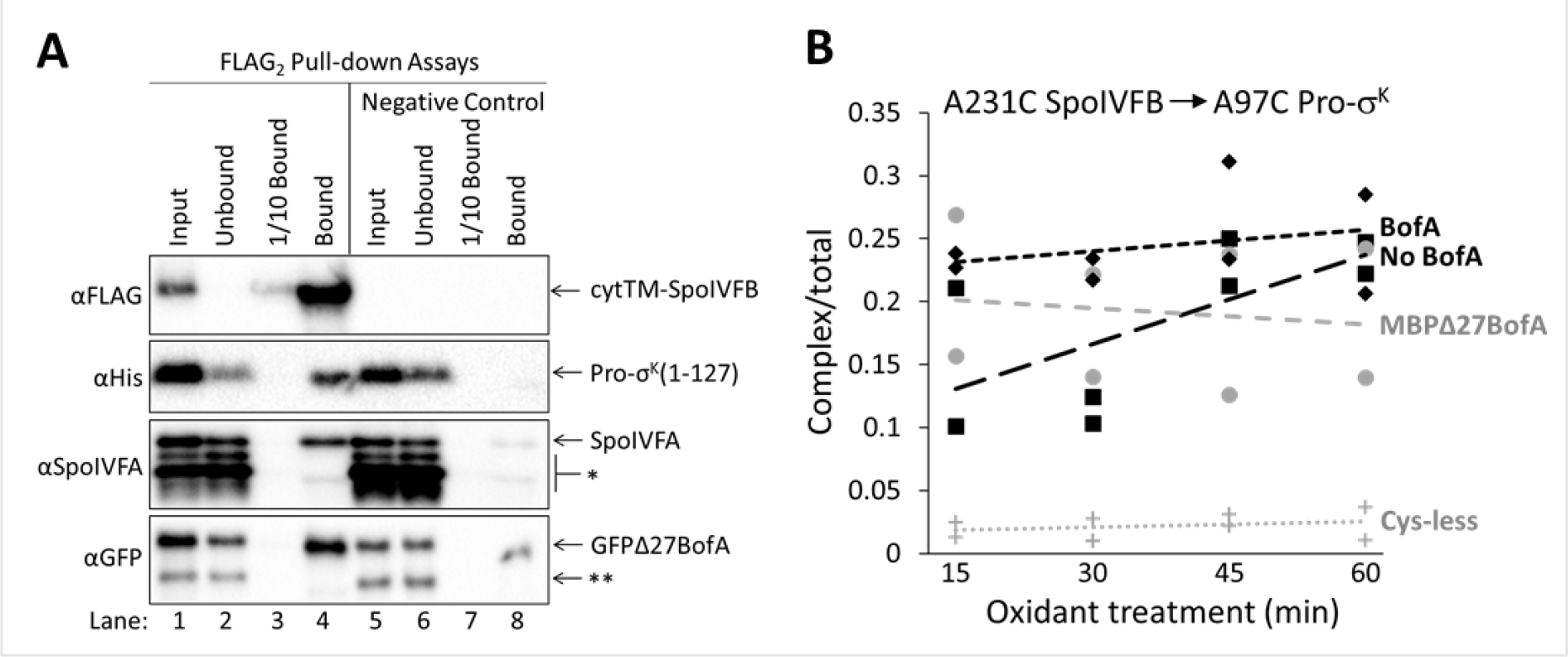
with 3 supplements Inhibitory proteins do not prevent Pro-σ^K^(1-127) from interacting with SpoIVFB. (**A**) Pro-σ^K^(1-127), SpoIVFA, and GFPΔ27BofA co-purify with cytTM-SpoIVFB. pET Quartet plasmids were used to produce a catalytically-inactive E44C cytTM-SpoIVFB variant with a FLAG2 tag (pSO73), or a variant lacking FLAG2 as a negative control (pSO149), in combination with Pro-σ^K^(1-127), SpoIVFA, and GFPΔ27BofA in *E. coli*. Samples collected after 2 h of IPTG induction were subjected to co-immunoprecipitation with anti-FLAG antibody beads. Input, unbound, 1/10 bound (diluted to match input), and (undiluted) bound samples were subjected to immunoblot analysis with FLAG, penta-His, SpoIVFA, and GFP antibodies. The single star (*) indicates cross-reacting proteins below SpoIVFA and the double star (**) indicates a cross-reacting protein or breakdown species of GFPΔ27BofA that fail to co-purify. A representative result from two biological replicates is shown. (**B**) A231C in the cytTM-SpoIVFB CBS domain forms disulfide cross-links with A97C in the C-terminal region of Pro-σ^K^(1-127) in the absence or presence of inhibitory proteins. pET Duet plasmids were used to produce single- Cys A231C cytTM-SpoIVFB E44Q in combination with single-Cys A97C Pro-σ^K^(1-127) (pSO130; squares and line with long dashes labeled “No BofA”, although SpoIVFA is also absent) or with Cys-less Pro-σ^K^(1-127) as a negative control (pSO255; crosses and line labeled “Cys-less”) in *E. coli*. pET Quartet plasmids were used to produce single-Cys A231C cytTM- SpoIVFB E44Q, single-Cys A97C Pro-σ^K^(1-127), and Cys-less SpoIVFA in combination with Cys-less MBPΔ27BofA (pSO133; circles and line labeled “MBPΔ27BofA”) or with Cys-less full-length BofA (pSO246; diamonds and line labeled “BofA”) in *E. coli*. Samples collected after 2 h of IPTG induction were treated for 15, 30, 45, or 60 min with Cu^2+^(phenanthroline)3 oxidant to promote disulfide bond formation and subjected to immunoblot analysis with FLAG antibodies to visualize the cytTM-SpoIVFB monomer, dimer, and complex with Pro-σ^K^(1-127) (Figure 5-figure supplement 3C). Abundance of the complex was divided by the total amount of cytTM-SpoIVFB monomer, dimer, and complex. The ratio over time was plotted (n=2) with a best-fit trend line. Figure 5-source data 1 Immunoblot images (raw and annotated) (Figure 5A) and quantification of cross-linking (Figure 5B).

We also performed pull-down assays with cobalt resin, which binds to the His6 tag on Pro-σ^K^(1–127), and we performed both types of pull-down assays (i.e., anti-FLAG antibody beads and cobalt resin) when full-length BofA rather than GFPΔ27BofA was co-produced with the other three proteins (Figure 5-figure supplement 1). Neither BofA nor GFPΔ27BofA when co-produced with SpoIVFA prevented Pro-σ^K^(1–127) from interacting with the cytTM-SpoIVFB variant. However, GFPΔ27BofA and SpoIVFA reduced co-purification of full-length Pro-σ^K^- His6 with the cytTM-SpoIVFB variant in both types of pull-down assays (Figure 5-figure supplement 2), as compared with Pro-σ^K^(1–127) (Figure 5A and Figure 5-figure supplement 1), suggesting that the C-terminal half of Pro-σ^K^ affects complex formation.

To explain the presence of substrate in the pulled-down protein complexes, but the absence of substrate cleavage (Figure 1B), we hypothesized that inhibitory proteins block substrate access to the SpoIVFB active site cleft, but do not prevent substrate interaction with the soluble C-terminal CBS domain of SpoIVFB (Figure 1A). The model of catalytically-inactive SpoIVFB E44Q in complex with Pro-σ^K^(1–127) predicts extensive interactions between the SpoIVFB CBS domain and the Pro-σ^K^(1–127) C-terminal region (56). We used the model to guide testing of single-Cys variants of the two proteins for disulfide cross-link formation. We discovered that A231C in the cytTM-SpoIVFB CBS domain can be cross-linked to A97C in the Pro-σ^K^(1–127) C-terminal region when *E. coli* co-producing the proteins are treated with oxidant (Figure 5-figure supplement 3A). We showed that the Cys substitutions do not impair cytTM- SpoIVFB activity or Pro-σ^K^(1–127) susceptibility to cleavage (Figure 5-figure supplement 3B). Finally, we measured time-dependent cross-linking in the presence or absence of Cys-less inhibitory proteins. Co-production of inhibitory proteins had little or no effect on formation of cross-linked complex (Figure 5B and Figure 5-figure supplement 3C). These results suggest that neither full-length BofA nor MBPΔ27BofA, when co-produced with SpoIVFA, prevent Pro- σ^K^(1–127) from interacting with the CBS domain of SpoIVFB in *E. coli*, consistent with the results of our pull-down assays (Figure 5A and Figure 5-figure supplement 1).

### Inhibitory Proteins Block Access of the Substrate N-terminal Proregion to the SpoIVFB Active Site

SpoIVFB cleaves Pro-σ^K^ (49) and Pro-σ^K^(1–127) (37) between residues S21 and Y22. In disulfide cross-linking experiments, Cys substitutions for several residues near the cleavage site in otherwise Cys-less Pro-σ^K^(1–127) formed a cross-linked complex with single-Cys (E44C, V70C, or P135C) cytTM-SpoIVFB variants (55). The complex was most abundant with the E44C and V70C variants, so we compared these interactions in the presence or absence of Cys-less inhibitory proteins.

SpoIVFB E44 is presumed to activate a water molecule for substrate peptide bond hydrolysis at the enzyme active site (46, 47). To test access of the substrate Proregion to the enzyme active site, we first measured time-dependent crosslinking between single-Cys E44C cytTM-SpoIVFB and single-Cys (F18C, V20C, S21C, K24C) Pro-σ^K^(1–127) variants co-produced in the absence of inhibitory proteins in *E. coli*. The V20C and K24C Pro-σ^K^(1–127) variants formed abundant complex that increased over time, but the F18C and S21C variants formed much less complex, only slightly more than the Cys-less Pro-σ^K^(1–127) negative control (Figure 6A and Figure 6-figure supplement 1A). Figure 6B shows a representative immunoblot (60-min oxidant treatment). When Cys-less MBPΔ27BofA and SpoIVFA were co-produced, the V20C and K24C Pro-σ^K^(1–127) variants formed much less complex and its abundance did not increase over time (Figure 6C and Figure 6-figure supplement 1B). Figure 6D shows a representative immunoblot (60-min oxidant treatment) for comparison with Figure 6B. When Cys-less full-length BofA and SpoIVFA were co-produced, similar decreases in abundance of the complex were observed (Figure 6-figure supplement 2). Figure 6 E and F summarize the cross-linking time courses with and without inhibitory proteins for the V20C and K24C Pro- σ^K^(1–127) variants. The effects of MBPΔ27BofA (lacking TMS1) and full-length BofA were indistinguishable. These results likely explain why BofA TMS1 is dispensable for most of the inhibitory function of BofA (36, 37) (Figure 1B and 3A). We conclude that inhibitory proteins block access of the substrate N-terminal Proregion to the SpoIVFB active site.

**Figure 6.**
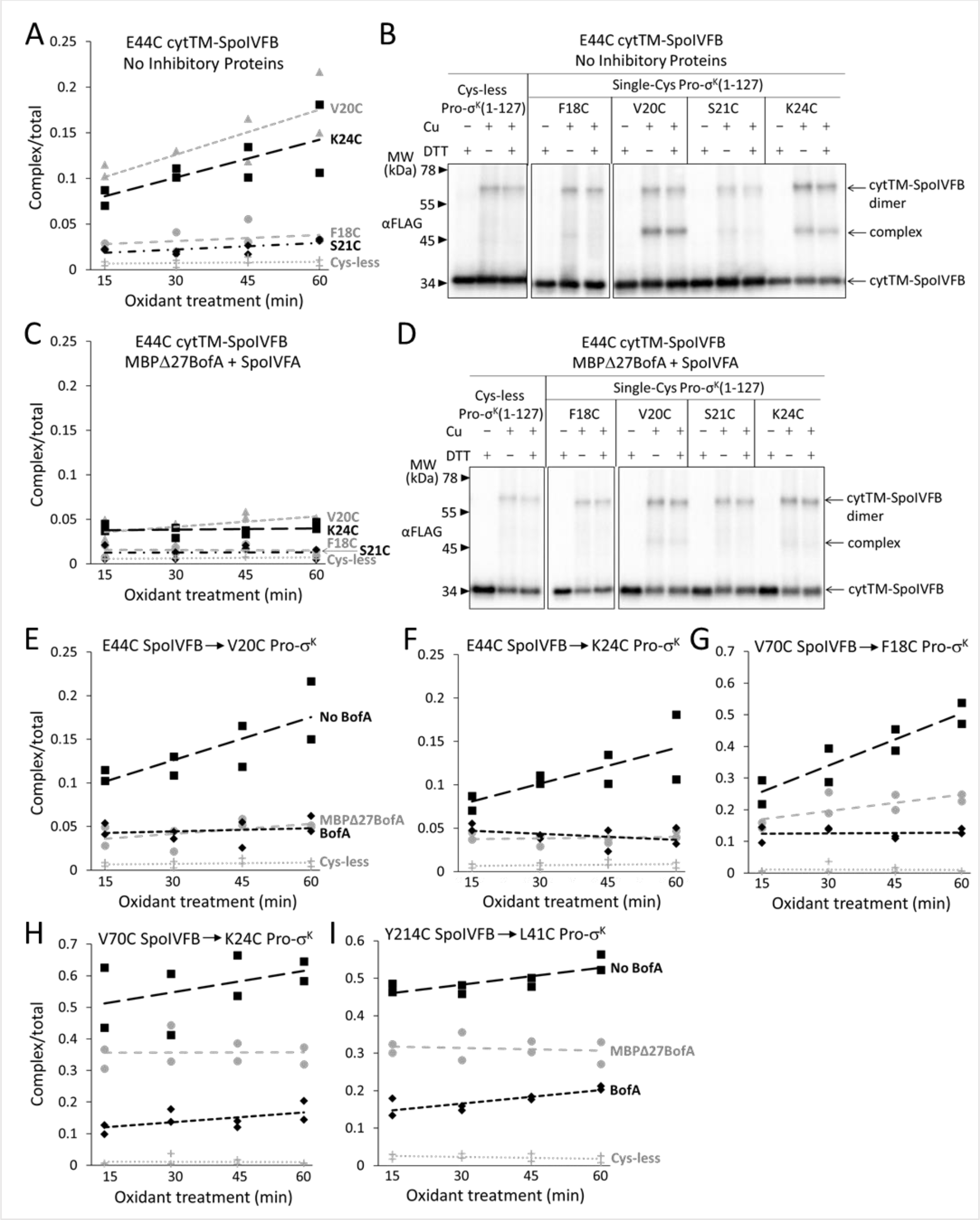
with 5 supplements Inhibitory proteins block access of the Pro-σ^K^(1-127) N-terminal region to the SpoIVFB active site and hinder interaction with the SpoIVFB membrane-reentrant loop and interdomain linker. (**A**) E44C at the cytTM-SpoIVFB active site forms abundant disulfide cross-links with V20C or K24C in the N-terminal region of Pro-σ^K^(1-127) in the absence of inhibitory proteins. pET Duet plasmids were used to produce single-Cys E44C cytTM-SpoIVFB in combination with single-Cys F18C (pSO167), V20C (pSO169), S21C (pSO170), or K24C (pSO128) Pro-σ^K^(1-127), or with Cys-less Pro-σ^K^(1-127) (pSO79) as a negative control, in *E. coli*. Samples collected after 2 h of IPTG induction were treated with Cu^2+^(phenanthroline)3 oxidant for 15, 30, 45, or 60 min to promote disulfide bond formation and subjected to immunoblot analysis with FLAG antibodies to visualize the cytTM-SpoIVFB monomer, dimer, and complex with Pro-σ^K^(1-127) (Figure 6- figure supplement 1A). Abundance of the complex was divided by the total amount of cytTM- SpoIVFB monomer, dimer, and complex. The ratio over time was plotted (n=2) with a best-fit trend line. (**B**) Representative immunoblots of 60-min samples from the experiment described in (**A**). (**C**) MBPΔ27BofA and SpoIVFA decrease cross-linking between E44C cytTM-SpoIVFB and V20C or K24C Pro-σ^K^(1-127). pET Quartet plasmids were used to produce single-Cys E44C cytTM-SpoIVFB in combination with single-Cys F18C (pSO163), V20C (pSO165), S21C (pSO166), or K24C (pSO131) Pro-σ^K^(1-127), or with Cys-less Pro-σ^K^(1-127) (pSO110) as a negative control, and Cys-less variants of MBPΔ27BofA and SpoIVFA in *E. coli*. Samples collected after 2 h of IPTG induction were treated and subjected to immunoblot analysis as in (**A**) (Figure 6-figure supplement 1B). The complex/total ratio was plotted as in (**A**). (**D**) Representative immunoblots of 60-min samples from the experiment described in (**C**). (**E** and **F**) Summaries of the effects of inhibitory proteins on cross-linking between E44C cytTM-SpoIVFB and V20C or K24C Pro-σ^K^(1-127). Data from (**A**) (labeled “No BofA” in (**E**), although SpoIVFA is also absent), (**C**) (labeled “MBPΔ27BofA” in (**E**), although SpoIVFA is also present), and Figure 6-figure supplement 2 (labeled “BofA” in (**E**), although SpoIVFA is also present) are plotted along with Cys-less Pro-σ^K^(1-127) as a negative control. In (**F**), symbols and lines are as in (**E**). (**G** and **H**) Summaries of the effects of inhibitory proteins on cross-linking between V70C in the cytTM-SpoIVFB membrane-reentrant loop and F18C or K24C in the Pro- σ^K^(1-127) N-terminal region. Data from Figure 6-figure supplement 3 are plotted using symbols and lines as in (**E**). (**I**) Summary of the effects of inhibitory proteins on cross-linking between Y214C in the cytTM-SpoIVFB interdomain linker and L41C in the Pro-σ^K^(1-127) N-terminal region. Data from Figure 6-figure supplement 5 are plotted using symbols and lines as in (**E**). Figure 6-source data 1 Quantification of cross-linking (Figure 6 A, C, and E-I).

SpoIVFB V70 is located in a predicted membrane-reentrant loop (Figure 1A), which may bind to the Proregion and present it to the active site for cleavage (56) based on a study *E. coli* RseP (66). Single-Cys V70C cytTM-SpoIVFB E44Q formed abundant cross-linked complex with F18C and K24C Pro-σ^K^(1–127) variants in the absence of inhibitory proteins (55) (Figure 6 G and H; Figure 6-figure supplement 3 A and B). When Cys-less MBPΔ27BofA and SpoIVFA were co-produced, less complex formed (Figure 6 G and H; Figure 6-figure supplement 3 C and D). Cys-less full-length BofA and SpoIVFA further decreased complex formation (Figure 6 G and H; Figure 6-figure supplement 3 E and F) and caused four novel species to be observed, including with the Cys-less Pro-σ^K^(1–127) negative control (Figure 6-figure supplement 3G), perhaps due to cross-linking of the V70C cytTM-SpoIVFB variant to *E. coli* proteins (i.e., BofA may cause the SpoIVFB membrane-reentrant loop to be exposed). Since full-length BofA hindered cross-linking more than MBPΔ27BofA (lacking TMS1) (Figure 6 G and H), both TMSs of BofA appear to interfere with the normal interaction between the SpoIVFB membrane-reentrant loop and the substrate Proregion.

Since full-length BofA also inhibited cleavage of Pro-σ^K^(1–127) in *E. coli* (Figure 1B) and Pro-σ^K^ in *B. subtilis* (Figure 3A) slightly more than GFPΔ27BofA (lacking TMS1), we compared time-dependent cross-linking of C46 of BofA and MBPΔ27BofA to single-Cys E44C and P135C cytTM-SpoIVFB variants. The results suggest that full-length BofA forms slightly more complex over time, whereas complex abundance did not increase over time with MBPΔ27BofA (Figure 6-figure supplement 4). Hence, BofA TMS1 may slightly enhance TMS2 occupancy in the SpoIVFB active site cleft.

To examine the extent to which inhibitory proteins hinder the interaction of the substrate with SpoIVFB, we identified a residue in the linker between the membrane and CBS domains of SpoIVFB (Figure 1A) that forms disulfide cross-linked complexes with Pro-σ^K^(1–127). The model of catalytically-inactive SpoIVFB E44Q in complex with Pro-σ^K^(1–127) (56) was used to guide testing for disulfide cross-link formation and we found that A214C in the cytTM-SpoIVFB linker can be cross-linked to A41C in Pro-σ^K^(1-127) (Figure 6-figure supplement 5A). We showed that the Cys substitutions do not impair cytTM-SpoIVFB activity or Pro-σ^K^(1-127) susceptibility to cleavage (Figure 5-figure supplement 3B). Finally, we measured time-dependent cross-linking in the presence or absence of Cys-less inhibitory proteins. Interestingly, formation of cross-linked complex was hindered more by full-length BofA than by MBPΔ27BofA, when co-produced with SpoIVFA (Figure 6I and Figure 6-figure supplement 5B), similar to cross-linking between V70C in the cytTM-SpoIVFB membrane-reentrant loop and F18C or K24C near the cleavage site in Pro-σ^K^(1-127) (Figure 6 G and H). The similar pattern suggests that in both cases BofA TMS2 partially interferes with the interaction and BofA TMS1 augments the interference.

### A Model of SpoIVFB in Complex with BofA and Parts of SpoIVFA and Pro-σ^K^

Computational models were generated using a similar protocol as described previously (56), but including additional constraints reflecting experimental cross-linking data reported herein, as well as newly predicted intra- and inter-chain contacts based on co-evolutionary couplings. Two final models were generated: 1) Full-length SpoIVFB modeled as a tetramer, with part of one Pro-σ^K^ molecule (residues 1-114), referred to as ‘fb.sigk’; 2) Full-length SpoIVFB, again modeled as a tetramer, with one molecule each of full-length BofA and parts of Pro-σ^K^ (residues 38-114) and SpoIVFA (residues 65-111), referred to as ‘fb.sigk.bofa.fa’. The omitted residues of Pro-σ^K^ and SpoIVFA could not be placed with sufficient confidence. The first model is described in Figure 7-figure supplement 1.

**Figure 7.**
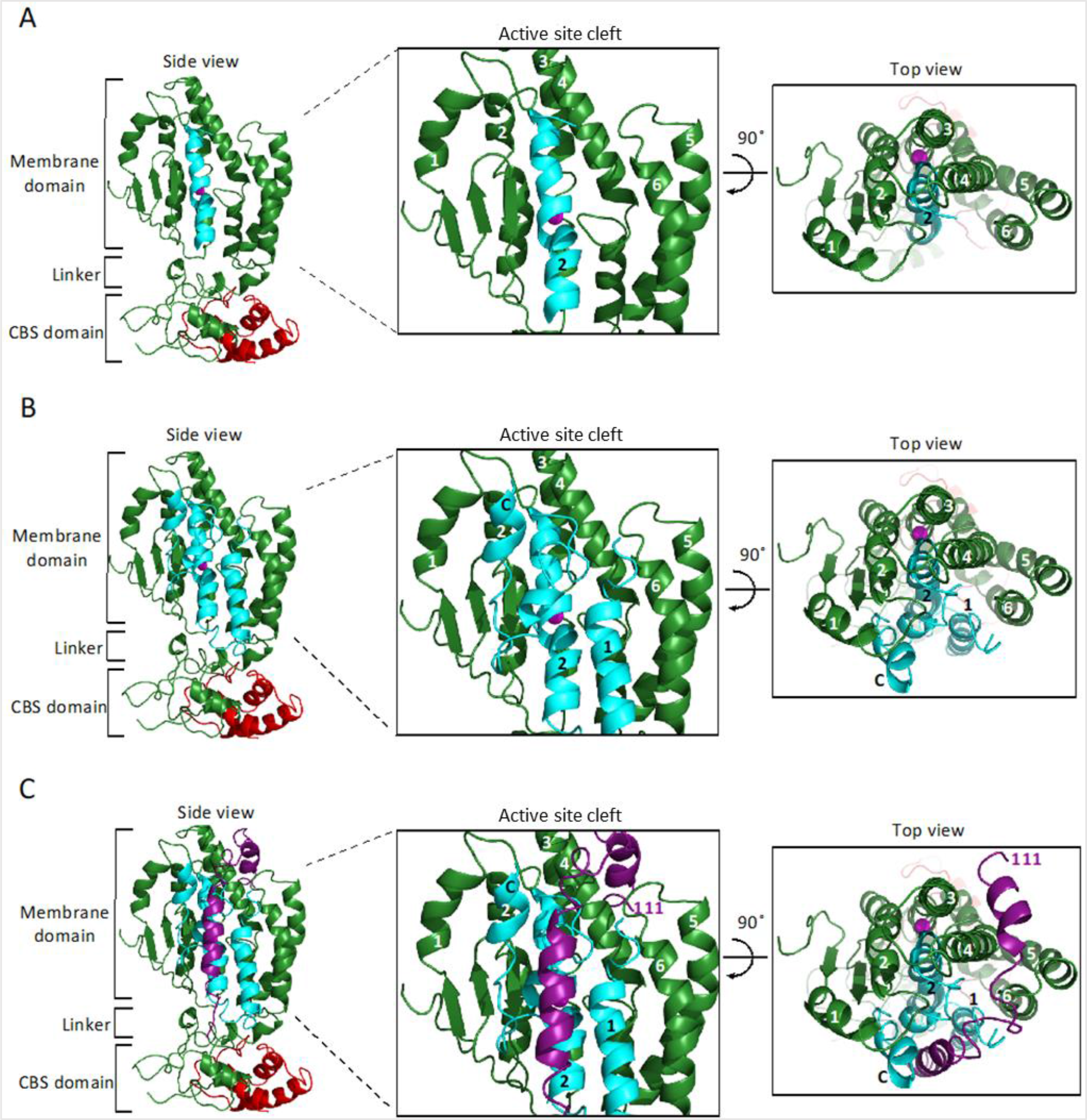
with 1 supplement Model of SpoIVFB monomer with BofA and parts of SpoIVFA and Pro-σ^K^. (**A**) Model of SpoIVFB, BofA TMS2, and the C-terminal part of Pro-σ^K^(1-127). At *Left*, a side view of the complex, showing the six TMSs of the SpoIVFB membrane domain with the active site zinc ion (magenta), the interdomain linker, and the CBS domain (green), BofA TMS2 (cyan), and Pro-σ^K^(38-114) (red). In the enlarged view of the active site cleft (*Center*), TMSs 1– 6 of SpoIVFB and TMS2 of BofA are numbered. At *Right*, a top view is shown. (**B**) Model of SpoIVFB with full-length BofA and Pro-σ^K^(38-114). Predicted TMSs 1 and 2 of BofA are numbered and its C-terminal region is labeled “C” near the C-terminus in the views shown in the *Center* and at *Right*. (**C**) Model of SpoIVFB with full-length BofA, SpoIVFA(65-111) (purple, residue 111 is numbered), and Pro-σ^K^(38-114). Figure 7 -source data 1 PyMOL session file used to produce the images and PDB file of the model of a SpoIVFB tetramer with BofA and parts of SpoIVFA and Pro-σ^K^.

Relevant features of the second model are illustrated in Figure 7 (only one molecule of SpoIVFB is shown). The first side view shows SpoIVFB with BofA TMS2 and the C-terminal part of Pro-σ^K^(1–127) (Figure 7A). The membrane domain of SpoIVFB (green) interacts with BofA TMS2 (cyan) while the interdomain linker and CBS domain of SpoIVFB interact with the modeled portion of Pro-σ^K^(1–127) (red). The enlarged view of the SpoIVFB active site cleft shows BofA TMS2 surrounded by SpoIVFB (TMSs labeled 1-6). The top view of the membrane domain emphasizes proximity between BofA TMS2, SpoIVFB TMSs 2-4, and the zinc ion (magenta) involved in catalysis.

The second side view shows SpoIVFB with full-length BofA and the C-terminal part of Pro-σ^K^(1–127) (Figure 7B). Our model predicts that BofA contains 2 TMSs and a membrane-embedded C-terminal region (labeled C near the C-terminus in the enlarged view of the SpoIVFB active site cleft) that forms two short α-helices connected by a turn. The enlarged and top views show that BofA interacts extensively with SpoIVFB and occupies its active site cleft, which would sterically hinder access of the Proregion of Pro-σ^K^. The side chain of conserved residue N48 in BofA TMS2 is predicted to interact with the side chain of T64, which is located in a loop that also includes N61 and precedes the first short α-helix of the C-terminal region.

The N61 side chain points away from the N48 and T64 side chains in the model, but the orientation of the N61 side chain is uncertain given the predicted loop structure, so an interaction with the side chains of N48 and/or T64 remains possible. Alternatively, the three conserved residues may contact other residues within BofA, based on co-evolutionary couplings, to stabilize the BofA structure and promote interactions with SpoIVFA and SpoIVFB. The three proteins form a heterotrimeric membrane complex that improves accumulation of each protein likely by inhibiting proteolytic degradation (36). Ala substitutions for N48, N61, and T64 in GFPΔ27BofA reduced its level as well as the levels of SpoIVFA and SpoIVFB, both upon heterologous expression in *E. coli* (Figure 2) and during *B. subtilis* sporulation (Figure 3A), suggesting that assembly of the inhibition complex was impaired. In agreement, the Ala substitutions caused partial or complete mislocalization of GFPΔ27BofA during sporulation (Figure 3B), which normally relies on SpoIVFA (36). Impaired assembly of the inhibition complex can explain the observed cleavage of Pro-σ^K^(1–127) in *E. coli* (Figure 2) and premature cleavage of Pro-σ^K^ in sporulating *B. subtilis* (Figure 3A).

Figure 7C shows the addition of the modeled portion of SpoIVFA (purple), which is predicted by co-evolutionary couplings to contact the BofA C-terminal region and SpoIVFB TMS4. Hence, the model predicts that SpoIVFA stabilizes the interaction of BofA with SpoIVFB. Since GFPΔ27BofA has the C-terminal region predicted to interact with SpoIVFA, the interaction of GFPΔ27BofA with SpoIVFB is likewise predicted to be stabilized by SpoIVFA, which may explain the dependence of SpoIVFB inhibition on both GFPΔ27BofA and SpoIVFA (Figure 1C), and why GFPΔ27BofA inhibits substrate cleavage (Figure 1B and 3A) and MBPΔ27BofA occupies the SpoIVFB active site cleft (Figure 6-figure supplement 4) nearly as well as full-length BofA.

## Discussion

Our results provide evidence that BofA TMS2 occupies the SpoIVFB active site cleft. Both inhibitory proteins block access of the substrate N-terminal Proregion to the SpoIVFB active site, but do not prevent interaction between the C-terminal region of Pro-σ^K^(1–127) and the SpoIVFB CBS domain. The mechanism of SpoIVFB inhibition is novel in comparison with previously known mechanisms of IP regulation. Structural modeling predicts that conserved BofA residues N48 and T64 interact to stabilize TMS2 and a membrane-embedded C-terminal region.

SpoIVFA is predicted to contact the BofA C-terminal region and SpoIVFB TMS4, bridging the two proteins to stabilize the inhibition complex. The model has clear implications for relief of SpoIVFB and its orthologs from inhibition during sporulation, as well as for IP inhibitor design.

### A Novel Mechanism of Intramembrane Protease Regulation

IPs were known to be regulated by three mechanisms – substrate localization, substrate extramembrane domain cleavage, and substrate interaction with an adapter protein. Substrate localization to a different organelle than its cognate IP is common in eukaryotic cells (3, 51). An early example of this regulatory mechanism emerged from studies of S2P. SREBP substrates are retained in the endoplasmic reticulum by SCAP and Insig proteins, then transported to the Golgi when more cholesterol is needed (67) (Figure 8).

**Figure 8.**
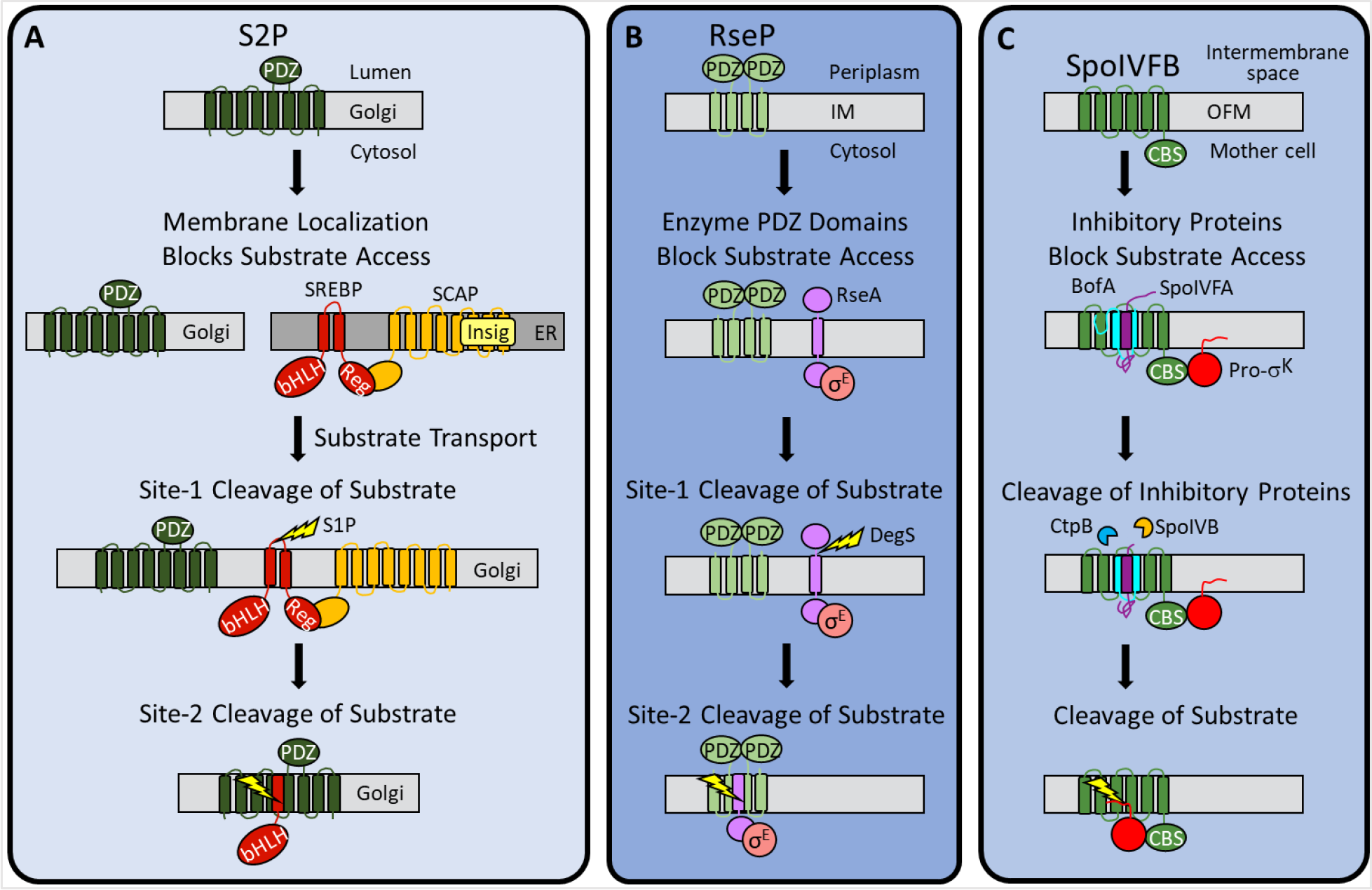
Regulation of intramembrane proteolysis by SpoIVFB differs from that of S2P and RseP. (**A**) S2P regulation. Human S2P localizes to membranes of the Golgi apparatus and has eight predicted TMSs and a PDZ domain (20, 101, 102). SREBP substrates localize to endoplasmic reticulum (ER) membranes *via* interaction of their regulatory domain (Reg) with SCAP, which is bound by Insig proteins when the cholesterol level is high (15). Cholesterol depletion releases SREBP-SCAP complexes from Insig and allows vesicular substrate transport to the Golgi membranes (103-105). Cleavage (lightning bolt) by S1P at site-1 in the luminal loop of SREBPs promotes separation of the TMSs and site-2 cleavage by S2P (106, 107). S2P cleavage releases the basic helix-loop-helix (bHLH) domain of SREBP substrates into the cytosol, for subsequent entry into the nucleus and transcriptional activation of genes required for fatty acid and cholesterol synthesis and uptake (108). (**B**) RseP regulation. RseP localizes to the *E. coli*inner membrane (IM) and has four predicted TMSs and tandem PDZ domains (20, 109, 110). The substrate RseA also localizes to the inner membrane and sequesters σ^E^ in the absence of envelope stress (111). The PDZ domains of RseP block access of the RseA/σ^E^ complex to the RseP active site (69). Stress causes misfolding of outer membrane proteins and accumulation of lipopolysaccharide intermediates, which trigger site-1 cleavage by DegS in the RseA periplasmic domain (68, 112). Removal of the RseA extramembrane domain permits entry into the RseP active site cleft for site-2 cleavage (113, 114). Release of RseA/σ^E^ into the cytosol allows ClpXP protease to degrade RseA, liberating σ^E^ to direct transcription of stress-responsive genes (115, 116). (**C**) SpoIVFB regulation. SpoIVFB localizes to the outer forespore membrane (OFM) during *B. subtilis* sporulation and has six predicted TMSs and a CBS domain (20, 32, 34, 54). Our results support a novel regulatory mechanism in which the inhibitory proteins BofA and SpoIVFA occupy the SpoIVFB active site cleft and block access of the Pro-σ^K^ Proregion. The inhibitory proteins did not prevent interaction of Pro-σ^K^ with the SpoIVFB CBS domain in our *E. coli* system. Proteases SpoIVB and CtpB are secreted from the forespore into the intermembrane space and cleave the inhibitory proteins (41–43, 45), allowing the Pro-σ^K^ Proregion to enter the SpoIVFB active site cleft for cleavage (46–48), which releases σ^K^ into the mother cell to direct transcription of sporulation genes (49, 50).

RIP of SREBPs also involves extramembrane loop cleavage, referred to as site-1 cleavage, prior to site-2 cleavage by S2P (1) (Figure 8). Substrates of bacterial metallo IPs (with the exception of SpoIVFB) typically also require initial regulated site-1 cleavage by another protease (19). A well-studied example in *E. coli* involves envelope stress regulating site-1 cleavage of the RseA extramembrane domain by DegS, allowing site-2 cleavage by RseP (20, 68) (Figure 8). Substrates of aspartyl IPs likewise require an initial cleavage by another protease, referred to as ectodomain shedding (52, 53). Shedding and site-1 cleavage of RseA allow substrate entry to the IP active site cleft. Without the initial cleavage, the substrate extramembrane domain prevents entry, due to size exclusion by the IP. In RseP, tandem PDZ domains act as a size-exclusion filter (69) (Figure 8). The nicastrin subunit plays a similar role in the metazoan aspartyl IP γ-secretase (70).

Only two cases of IP regulation by substrate interaction with an adapter protein have been reported. In the bacterium *Mycobacterium tuberculosis*, the adapter protein Ppr1 bridges the metallo IP Rip1 to one of its substrates, but not to others (21). The γ-secretase subunits nicastrin and PEN-2 also appear to interact with substrate before passage to the active site (71).

Our results support a novel mechanism of IP regulation – inhibitory proteins block substrate access to the SpoIVFB active site (Figure 8). Many soluble proteases are regulated by inhibitory proteins, prodomains, or prosegments that sterically hinder substrate access to their active site (62), but analogous regulation of an IP has not been reported previously. In addition to steric hindrance, inhibition of soluble metalloproteases often involves a protein residue side chain coordinating the catalytic metal ion (62, 63). We hypothesized that BofA N48 coordinates the predicted zinc ion at the SpoIVFB active site since Asn is highly conserved at the corresponding position in BofA orthologs (Figure 2-figure supplement 1) and BofA N48 is crucial for SpoIVFB inhibition (Figure 2 and 3). Consistent with our hypothesis, BofA TMS2 with N48 near its center occupies the SpoIVFB active site cleft, based on disulfide cross-linking experiments (Figure 4 and Figure 4-figure supplement 3-5). However, the N48 side chain points away from the zinc ion in our structural model, which instead predicts that the N48 side chain interacts with the T64 side chain, another conserved BofA residue shown to be important for SpoIVFB inhibition (Figure 2 and 3). Therefore, we favor a simple steric hindrance mechanism of inhibition not involving a BofA residue acting as a zinc ligand.

In addition to BofA TMS2 sterically hindering substrate access to the SpoIVFB active site, what are the roles of other parts of BofA in the novel inhibition mechanism? It was known that deletion of three residues from the C-terminal end of BofA caused a loss of function or stability in *B. subtilis* (33, 60). We found that deleting the three residues or changing them to Ala in GFPΔ27BofA caused loss of SpoIVFB inhibition in our *E. coli* system (Figure 2-figure supplement 2A). Our structural model predicts that the three BofA residues interact with SpoIVFA and SpoIVFB (Figure 7C), which can be tested by cross-linking experiments.

BofA TMS1 appears to sterically hinder some interactions between the Pro-σ^K^ N-terminal region and SpoIVFB, based on comparison between full-length BofA and MBPΔ27BofA (lacking TMS1) in cross-linking time courses. The interactions hindered more by full-length BofA included cross-links between the Pro-σ^K^ Proregion and the SpoIVFB membrane-reentrant loop (Figure 6G and H), which may bind to the Proregion and present it to the SpoIVFB active site for cleavage (56) based on a study of the homologous membrane-reentrant loop of *E. coli* RseP (66). Full-length BofA also hindered cross-links between Pro-σ^K^ L41C and the SpoIVFB interdomain linker (Y214C) more than MBPΔ27BofA (Figure 6I), suggesting that BofA TMS1 weakens the interaction between the interdomain linker and Pro-σ^K^, which is crucial for cleavage (56, 59). Our model predicts that both TMSs of BofA, as well as its C-terminal region, occupy the SpoIVFB active site cleft (Figure 7B), plausibly explaining the additional steric hindrance that BofA TMS1 appears to provide. Yet, TMS1 plays a minor role in SpoIVFB inhibition compared with BofA TMS2 and its C-terminal region, since full-length BofA inhibited cleavage of Pro-σ^K^(1–127) in *E. coli* (Figure 1B) and Pro-σ^K^ in *B. subtilis* (Figure 3A) only slightly more than GFPΔ27BofA. Nevertheless, the minor role of BofA TMS1 may be important as premature σ^K^ production can reduce sporulation efficiency (31).

### Implications for Activation of SpoIVFB and Its Orthologs in Other Endospore Formers

Inhibition of SpoIVFB is relieved during *B. subtilis* sporulation when SpoIVB cleaves the C-terminal end of SpoIVFA (41–44) and CtpB cleaves the C-terminal ends of both SpoIVFA and BofA (42, 43, 45) (Figure 1-figure supplement 1). The levels of inhibitory proteins decrease coincident with increasing Pro-σ^K^ cleavage (38, 41, 42, 45), suggesting that degradation of SpoIVFA and BofA permits SpoIVFB activity. Alternatively, a conformational change in the inhibition complex may suffice to trigger SpoIVFB activity, based on *in vitro* experiments indicating that SpoIVB cleavage of SpoIVFA alters susceptibility of the complex to general proteolysis but does not disrupt the complex (42).

Our results imply that activation of SpoIVFB requires removal or substantial movement of inhibitory proteins to allow substrate access to the active site. Foremost, steric hindrance by BofA TMS2 must be relieved. Additionally, the critical roles of the BofA C-terminal region and SpoIVFA must be explained. Our structural model predicts that SpoIVFA bridges the BofA C-terminal region and SpoIVFB TMS4 to stabilize the inhibition complex (Figure 7C). We propose that cleavage of SpoIVFA by SpoIVB initiates destabilization of the inhibition complex. The initial cleavage, perhaps followed by further proteolysis of SpoIVFA, may destabilize the membrane-embedded BofA C-terminal region and expose it to cleavage by CtpB, which hastens SpoIVFB activation but is not essential (40, 42, 43, 45). Finally, degradation of BofA or a conformational change would relieve steric hindrance by BofA TMS2 and allow substrate access to the SpoIVFB active site.

Whether BofA and SpoIVFA completely prevent Pro-σ^K^ from interacting with SpoIVFB prior to activation during sporulation is an open question. Pull-down and localization experiments suggested a complete block, but involved fusion proteins that may weaken the interaction between SpoIVFB and Pro-σ^K^ (59). Our experiments employed cytTM-SpoIVFB variants and heterologous expression in growing *E. coli.* Pull-down and cross-linking results showed that inhibitory proteins do not Pro-σ^K^(1–127) from interacting with cytTM-SpoIVFB (Figure 5 and Figure 5-figure supplement 1). Neither did GFPΔ27BofA and SpoIVFA prevent full-length Pro-σ^K^ from interacting with cytTM-SpoIVFB (Figure 5-figure supplement 2), although the interaction appeared to be weakened by the presence of the C-terminal half of Pro- sigK. Perhaps full-length BofA and SpoIVFA would completely prevent full-length Pro-σ^K^ from interacting with cytTM-SpoIVFB.

The mechanisms of inhibition and activation of *B. subtilis* SpoIVFB likely apply to its orthologs in spore-forming bacilli since these bacteria encode BofA, SpoIVFA, SpoIVB, and CtpB (23, 25). This group includes well-known human or plant pathogens such as *Bacillus anthracis*, *Bacillus cereus*, and *Bacillus thuringiensis*. In contrast, clostridia that form endospores do not have a recognizable gene for SpoIVFA (22, 23). In some clostridia, a non-orthologous gene has been proposed to code for a protein that performs the same function as SpoIVFA (23), so similar mechanisms of SpoIVFB regulation may apply. The *E. coli* system described herein will facilitate further testing of heterologous protein function. Many spore-forming clostridia also lack a recognizable gene for BofA (25). Our results imply that SpoIVFB activity would be unregulated in these bacteria (i.e., coordination between forespore and mother cell gene expression would be lost). A precedence for less precise control of σ^K^-dependent gene expression is known from studies of the human pathogen *Clostridioides difficile* (72, 73), in which the gene for σ^K^ does not encode a Proregion and a gene for SpoIVFB is absent (74). Based on phylogenomic analysis, the precise control of σ^K^-dependent gene expression observed in *B. subtilis* emerged early in evolution and has been maintained in bacilli which propagate primarily in aerobic environments, but has been modified or lost in many clostridia which are found mainly in anaerobic habitats (25).

### Design of IP Inhibitors

Efforts toward development of therapeutic peptidyl inhibitors of serine IPs (rhomboids) are well-advanced (11, 12, 75). Translational work on aspartyl IPs has focused on γ-secretase owing to its processing of the amyloid precursor protein (APP) associated with Alzheimer’s disease (76). Wide-spectrum inhibitors of γ-secretase exhibit toxicity in clinical trials, mainly due to inhibition of signaling *via* Notch receptors, which are also substrates of γ- secretase (77). The structures of γ-secretase complexes with APP and Notch reveal differences in binding that may allow substrate-specific inhibitors to be developed as therapeutics (13, 14). Stabilization of complexes in which γ-secretase progressively cleaves APP is another promising approach toward development of drugs to treat Alzheimer’s disease (78).

In comparison with rhomboids and γ-secretase, translational work on metallo IP inhibitors is in its infancy. Recently, batimastat, a peptidic hydroxamate known to inhibit eukaryotic matrix metalloproteases (MMPs), was shown to selectively inhibit *E. coli* RseP *in vivo* (79). A lack of selectivity has been a problem in efforts toward using peptidic hydroxamates as MMP inhibitors for treatment of cancer, arthritis, and other diseases (80–82). The hydroxamate strongly coordinates the catalytic metal ion and the peptidic portions have not provided enough specificity to prevent off-target effects. Even so, the side effects may not preclude using peptidic hydroxamates to treat bacterial infections topically, and systemically if administered briefly. Other small-molecule MMP inhibitors that have already been developed (80) should be screened for activity against bacteria and fungi with metallo IPs known to be involved in pathogenesis (18–21).

Our findings reveal concepts that may inform efforts to design selective inhibitors of IPs. BofA TMS2 appears to block the SpoIVFB active site (Figure 7A), similar to the prosegment of many latent proteases (62). Like BofA, some prosegments can act in *trans*, an observation that has led to the demonstration of prodomains as selective inhibitors of A Disintegrin And Metalloproteinase (ADAM) family enzymes (83, 84). Selectivity relies on the extensive interaction surface between the prodomain and the enzyme, including features specific to the pair, which can boost efficacy and diminish off-target effects in therapeutic strategies (82). A peptide encompassing residues 48 to 64 of BofA may inhibit SpoIVFB, but our model also predicts extensive interactions between the BofA C-terminal region (residues 65 to 87) and SpoIVFB (Figure 7B), so a longer BofA peptide, perhaps flexibly linked to SpoIVFA residues 96 to 109, which are also predicted to interact with SpoIVFB (Figure 7C), may exhibit improved inhibition and selectivity. Since cleavage of Pro-σ^K^(1–127) by cytTM-SpoIVFB has been reconstituted *in vitro* (48), it will be possible to test peptides for inhibition. Structure determination of SpoIVFB in complex with an inhibitory peptide, GFPΔ27BofA, or full-length BofA is an important goal to facilitate design of metallo IP inhibitors. In particular, it may inform efforts to design inhibitors of SpoIVFB orthologs in pathogenic bacilli and clostridia, which persist by forming endospores (27–29). Such efforts could lead to new strategies to control endosporulation.

Our findings also suggest that BofA TMS1 interferes with interactions between the Pro- σ^K^ N-terminal region and SpoIVFB. Therefore, a cyclic peptide inhibitor may prove to be more effective than a linear one. The desirable characteristics of cyclic peptides as therapeutics, and new methods of producing and screening cyclic peptide libraries, make this an attractive possibility (85, 86). Although we favor a simple steric hindrance mechanism of inhibition not involving a BofA residue acting as a zinc ligand, it is possible that inclusion of such a residue in a BofA TMS2 peptide mimic would improve SpoIVFB inhibition. A similar strategy could be used to design inhibitors of other metallo IPs.

## Materials and Methods

### Plasmids, Primers, and Strains

Plasmids used in this study are described in Supplementary File 1, as are primers used in plasmid construction. Plasmids were cloned in *E. coli* strain DH5α (87). Relevant parts of plasmids were verified by DNA sequencing with primers listed in Supplementary File 1 and *B. subtilis* strains used in this study are also described therein.

### Pro-σ^K^(1–127) Cleavage in *E. coli*

Strain BL21(DE3) (Novagen) was used to produce proteins in *E. coli*. Two plasmids with different antibiotic resistance genes were cotransformed (37) or a single plasmid was transformed, with selection on Luria-Bertani (LB) agar supplemented with kanamycin sulfate (50 μg/mL) and/or ampicillin (100 μg/mL). Transformants (4-5 colonies) were grown in LB medium with 50 μg/mL kanamycin sulfate and/or 200 μg/mL ampicillin at 37°C with shaking (200 rpm). Typically, overnight culture (200 μL) was transferred to 10 mL of LB medium with antibiotics, cultures were grown at 37°C with shaking (250 rpm) to an optical density of 60-80 Klett units, and isopropyl β-D-thiogalactopyranoside (IPTG) (0.5 mM) was added to induce protein production for 2 h. For transformants with either pET Quintet or full-length Pro-σ^K^-His6, overnight growth was avoided. Transformants were transferred directly to 10 mL of LB medium with antibiotic, and cultures were grown and induced as described above. Equivalent amounts of cells (based on optical density in Klett units) were collected (12,000 × *g* for 1 min) and extracts were prepared (37), then subjected to immunoblot analysis.

### Immunoblot Analysis

Samples were subjected to immunoblot analysis as described (38). Briefly, proteins were separated by SDS-PAGE using 14% Prosieve (Lonza) polyacrylamide gels and electroblotted to Immobilon-P membranes (Millipore). Protein migration was monitored using SeeBlue Plus2 Prestained Standard (Invitrogen) and blots were blocked with 5% nonfat dry milk (Meijer) in TBST (20 mM Tris-HCl pH 7.5, 0.5 M NaCl, 0.1% Tween 20) for 1 h at 25°C with shaking. Blots were probed with antibodies against His6 (penta-His Qiagen catalog #34460; 1:10,000), FLAG2 (Sigma catalog #A8592; 1:10,000), GFP (38) (1:10,000), MBP (NEB catalog #E8030S; 1:10,000), SpoIVFA (38) (1:3,000), Pro-σ^K^ (88) (1:3,000), and/or SpoIVFB (47, 56) (1:5,000) diluted in TBST with 2% milk, overnight at 4°C with shaking. Since the GFP, MBP, SpoIVFA, Pro-σ^K^, and SpoIVFB antibodies were not HRP-conjugated, they were detected with goat anti-rabbit-HRP antibody (Bio-Rad catalog #170-6515; 1:10,000) diluted in TBST with 2% milk, 1 h at 25°C with shaking. Signals were generated using the Western Lightning Plus ECL reagent (PerkinElmer) and detected using a ChemiDoc MP imaging system (Bio-Rad). Unsaturated signals were quantified using the Image Lab 5.1 software (Bio-Rad) lane and bands tool in order to determine the Pro-σ^K^(1–127) cleavage ratio or the ratio of disulfide cross-linked complex to the total intensity of the cytTM-SpoIVFB variant monomer, dimer, and complex.

### BofA Sequence Analysis

Orthologs of *B. subtilis bofA*, which are present in the genomes of most endospore-forming bacteria (23), were collected from the NCBI and Uniprot databases.

The protein sequences of BofA orthologs were aligned using the T-Coffee multiple sequence alignment package (89). Residues identical in at least 70% of the sequences were considered conserved.

### *B. subtilis* Sporulation and GFPΔBofA Localization

GFPΔ27BofA or its variants were expressed under control of the *bofA* promoter (P*bofA*) after ectopic chromosomal integration. Plasmids bearing P*bofA*-*gfpΔ27bofA* or its variant, bordered by regions of homology to *B. subtilis amyE*, were transformed into strain ZR264. Transformants with a gene replacement at *amyE* were selected on LB agar with spectinomycin sulfate (100 μg/mL) and identified by loss of amylase activity (90). Sporulation was induced by growing cells in the absence of antibiotics followed by the resuspension of cells in SM medium (90). At indicated times PS, samples (50 μL) were centrifuged (12,000 × *g* for 1 min), supernatants were removed, and cell pellets were stored at -80°C. Whole-cell extracts were prepared as described for *E. coli* (37), except samples were incubated at 50°C for 3 min instead of boiling for 3 min (56), and proteins were subjected to immunoblot analysis.

To image GFPΔ27BofA localization, samples collected at 3 h PS were examined by fluorescence microscopy using an Olympus FluoView FV-1000 filter-based confocal microscope. GFPΔ27BofA (ex/em ∼488/507 nm) was excited using a 458 nM argon laser and fluorescence was captured using a BA465-495 nm band pass filter. The lipophilic dye FM 4-64 (1 μg/mL) (AAT Bioquest) was used to stain membranes. FM 4-64 (ex/em ∼515/640 nm) was excited using a 515 nm argon laser and fluorescence was captured using a BA560IF band pass filter (91).

### Disulfide Cross-Linking

A method described previously (92) was used with slight modifications (55). As described above for Pro-σ^K^(1–127) cleavage, *E. coli* BL21(DE3) was transformed with a plasmid, grown in LB (10 mL), induced with IPTG, and equivalent amounts of cells were collected. Cells were mixed with chloramphenicol (200 μg/mL) and 2- phenanthroline (3 mM), collected by centrifugation (12,000 × *g* for 1 min), washed with 10 mM Tris-HCl pH 8.1 containing 3 mM 2-phenanthroline, and suspended in 10 mM Tris-HCl pH 8.1. Samples were treated with 1 mM Cu^2+^(phenanthroline)3 or 3 mM 2-phenanthroline (as a negative control) for 15, 30, 45 or 60 min at 37°C, followed by incubation with neocuproine (12.5 mM) for 5 min at 37°C. Cells were lysed and proteins were precipitated by addition of trichloroacetic acid (5%) and inversion every 5 min for 30 min on ice. Proteins were sedimented by centrifugation (12,000 × *g*) for 15 min at 4°C, the supernatant was removed, and the pellet was washed with cold acetone. The pellets were sedimented by centrifugation (12,000 × *g*) for 5 min at 4°C and the supernatants were discarded. The pellets were dried for 5 min at 25°C and resuspended in buffer (100 mM Tris-HCl pH 7.5, 1.5% SDS, 5 mM EDTA, 25 mM *N*- ethylmaleimide) for 30 min at 25°C. Portions were mixed with an equal volume of sample buffer (25 mM Tris-HCl pH 6.8, 2% SDS, 10% glycerol, 0.015% bromophenol blue) with or without 100 mM DTT, and were typically incubated at 37°C for 10 min, prior to immunoblot analysis. In experiments with single-Cys V70C cytTM-SpoIVFB E44Q, single-Cys Pro-σ^K^(1–127) variants, and Cys-less variants of BofA and SpoIVFA, samples were boiled 3 min prior to immunoblot analysis, which helped to resolve species (Figure 6-figure supplement 3 E and G).

### Co-Immunoprecipitation (FLAG2 Pull-Down Assays)

*E. coli* BL21(DE3) was transformed with a plasmid, grown in LB (1 L), and induced with IPTG as described above. The culture was split, cells were harvested, and cell pellets were stored at -80°C. Cell lysates were prepared as described (65), except that each cell pellet was resuspended in 20 mL of lysis buffer containing 50 mM Tris-HCl pH 7.1 rather than PBS. Cell lysates were centrifuged (15,000 × *g* for 15 min at 4 °C) to sediment cell debris and protein inclusion bodies. The supernatant was treated with 1% *n*-dodecyl-β-D-maltoside (DDM) (Anatrace) for 1 h at 4 °C to solubilize membrane proteins, then centrifuged at 150,000 × *g* for 1 h at 4 °C. The supernatant was designated the input sample and 1 mL was mixed with 50 uL anti-DYKDDDDK magnetic agarose (Pierce) that had been equilibrated with buffer (50 mM Tris-HCl pH 7.1, 0.1% DDM, 5 mM 2-mercaptoethanol, 10% glycerol) and the mixture was rotated for 1 h at 25°C. The magnetic agarose was removed with a DynaMag-2 magnet (Invitrogen) and the supernatant was saved (unbound sample). The magnetic agarose was washed three times by gently vortexing with 500 μL wash buffer (50 mM Tris-HCl pH 7.1, 150 mM NaCl, 10% glycerol, 0.1% DDM), then washed once with 500 μL water. The magnetic agarose was mixed with 50 μL of 2× sample buffer (50 mM Tris·HCl pH 6.8, 4% SDS, 20% glycerol, 200 mM DTT 0.03% bromophenol blue) and boiled for 3 min (bound sample). A portion of the bound sample was diluted tenfold (1/10 bound sample) with 1× sample buffer to match the concentration of the input sample. Samples were subjected to immunoblot analysis.

### Colbalt Affinity Purification (His6 Pull-Down Assays)

Input sample (15 mL) prepared as described above was mixed with imidazole (5 mM) and 0.5 mL of Talon superflow metal affinity resin (Clontech) that had been equilibrated with buffer (as above for magnetic agarose). The mixture was rotated for 1 h at 4°C. The cobalt resin was sedimented by centrifugation at 708 × *g* for 2 min at 4 °C and the supernatant was saved (unbound sample). The resin was washed three times with 5 mL wash buffer (as above plus 5 mM imidazole), each time rotating the mixture for 10 min at 4 °C and sedimenting resin as above. The resin was mixed with 0.5 mL 2× sample buffer and boiled for 3 min (bound sample). A portion of the bound sample was diluted fifteenfold (1/15 bound sample) with 1× sample buffer to match the concentration of the input sample. Samples were subjected to immunoblot analysis.

### Modeling of Complexes Containing SpoIVFB, BofA, and Parts of SpoIVFA and Pro-σ^K^

The modeling proceeded through stages where initial monomeric models were assembled step-by-step into multimeric complexes guided primarily by the restraints from cross-linking experiments and predicted contacts from co-evolutionary coupling analysis. More specifically, a SpoIVFB monomer was first assembled from the membrane and CBS domains. Two monomers were combined into a dimer and the dimer was assembled into a plausible tetramer. Part of one molecule of Pro-σ^K^ (residues 1-114) was subsequently added, which resulted in ‘fb.sigk’. The ‘fb.sigk.bofa.fa’ model was developed by starting from ‘fb.sigk’, truncating the Pro-σ^K^ N-terminus (residues 1-37), and subsequently adding first BofA and finally SpoIVFA (residues 65- 111).

The membrane domain of a SpoIVFB monomer was modeled initially based on the structure for the site-2 protease from *Methanocaldococcus jannaschii* (PDB code: 3B4R) (7). The structure was combined with a CBS domain modeled based on the CBS-domain protein TM0935 from *Thermotoga martima* (PDB code: 1O50) (93). The CBS domain provides a dimerization interface for the C-terminal part of SpoIVFB. The full-length dimer including the membrane domain was completed by considering predicted contacts from co-evolutionary coupling analysis. A tetramer was then built guided by the arrangement of transmembrane helices and orientation of CBS domains in the recently published full-length structure of the chloride proton exchanger CLC-7 (PDB code: 7JM6) (94) and again by considering contacts from co-evolutionary couplings. We note that the resulting dimer and tetramer models are different from our previous model for the SpoIVFB as we could take advantage of the new structural template and additional information from the co-evolutionary couplings. The initial model for the well-folded C-terminal part of Pro-σ^K^ (residues 40-114) was built based on the structure of RNA polymerase sigma subunit domain 2 (PDB code: 3UGO) (95). Suitable templates are not available for BofA and the part of SpoIVFA modeled here. For these components, initial models were obtained based on the predicted intra-chain contacts from the co-evolutionary couplings.

The modeling under restraints was initially carried out using Cα-based coarse-grained models that were allowed to relax under restraints *via* cycles of energy minimization and short molecular dynamics simulations at elevated temperature as described in more detail previously (56). Restraints from cross-linking were applied as described previously (56). Different from our previous work, we also applied extensive intra- and inter-chain restraints based on predicted contacts from co-evolutionary coupling analysis. In addition, we applied positional restraints to keep the structures of the SpoIVFB membrane and CBS domains and the folded C-terminus of Pro-σ^K^ close to their initial models while allowing subunits to move relative to each other as guided by the restraints. For BofA and SpoIVFA, individual helices were restrained internally but allowed to move relative to each other to find the optimal arrangement in complex with SpoIVFB, again guided by the predicted contact restraints. Contact restraints with respect to SpoIVFB were implemented as minimum distance restraints to the closest residue in any of the four subunits since contact predictions cannot distinguish between chemically equivalent oligomer units. All available experimental cross-links were applied but the list of predicted contacts was edited to exclude certain contacts incompatible with cross-linking data and previous biochemical data on the overall topology of the SpoIVFB-BofA-SpoIVFA complex when embedded into the membrane. Excluded contacts may reflect uncertainties in the prediction method as inter-chain contacts are more difficult to predict reliably. They may also indicate alternate biologically relevant interactions between SpoIVFB, BofA, and SpoIVFA that were not probed in the experiments *via* cross-linking. Detailed lists of all applied restraints are given in Supplementary Files 2-5.

Inter-residue contacts were predicted by trRosetta (96). A multiple sequence alignment (MSA), the input of trRosetta, for a protein was generated using HHblits (97) to search against the UniClust30 database (98). It was further curated to only include sequences that are related to spore formation and are from organisms which have both SpoIVFB and SpoIVFA sequences.

To predict inter-protein contacts using trRosetta, a hybrid MSA was generated by pasting two interacting proteins’ sequences that are from the same strain. Twenty glycines were inserted between the target sequences to prevent irrelevant predictions near the pasted regions due to the proximity in the hybrid sequence. Similarly, gaps were inserted for the homologous sequences to preserve alignment with the *B. subtilis* proteins. The contact predictions for the inserted residues were ignored for further modeling.

Finally, the Cα models for the complexes were converted to an all-atom model using PyRosetta (99) with the curated predicted intra- and inter-protein contacts and the cross-linking constraints. First, the model was locally minimized in the Rosetta centroid representation. The cross-linking constraints and inter-protein contacts were used for the Cα model building.

Predicted intra-protein contacts were also used only if their contact probabilities were higher than 0.15 and if they were not violated severely in the Cα model; a predicted contact was considered as a severe violation if its score was greater than 10 Rosetta energy units (REUs). The relative weights for each scoring term were 25, 0.25, and 0.1 for the cross-linking constraints, inter-protein contacts, and intra-protein contacts, respectively. To prevent large deviation from the Cα model, harmonic restraints were applied on every Cα atom with a force constant of 0.1 REU/Å^2^. In addition, Rosetta centroid energy terms were also used: Ramachandran energy (1.0), omega angle potential (0.5), backbone hydrogen bond energy (5.0), and van der Waals energy (1.0) with weights in the parentheses. The minimization was performed for 100 steps. Then, the minimized structure was converted to an all-atom model, and the FastRelax protocol was applied to the model with the Rosetta scoring function (ref2015) (100) and the same cross-linking constraints, the contact predictions, and the harmonic restraints on Cα atoms that were used for the minimization. Eight all-atom models were generated from the Cα model, and a model with the least cross-linking constraint violations was selected. To generate final models, Zn^2+^ ions were added at the active sites of each subunit of SpoIVFB to be coordinated with D137, H43, and H47 followed by another brief minimization using CHARMM under harmonic restraints on heavy atoms to accommodate the Zn^2+^ ions without clashes.

## Acknowledgments

We thank Daniel Parrell for assistance with fluorescence microscopy, Jon Kaguni for helpful comments on the manuscript, and David Rudner, Simon Cutting, Ruanbao Zhou, and Yang Zhang for plasmids. This research was supported by National Institutes of Health Grants R01 GM43585 (to L.K.) and R35 GM126948 (to M.F.), and by Michigan State University AgBioResearch.

## Figure Supplements

**Figure 1-figure supplement 1**

**Morphological changes during endosporulation and regulated intramembrane proteolysis of Pro-σ^K^ in *B. subtilis*.**

**Figure 1-figure supplement 2**

**Full-length BofA alone fails to inhibit Pro-σ^K^(1-127) cleavage in *E. coli* and full-length Pro- σ^K^ is similar to Pro-σ^K^(1-127) in terms of requirements for cleavage inhibition.**

**Figure 1-figure supplement 3**

**SpoIVB partially relieves inhibition of SpoIVFB by BofA and SpoIVFA in *E. coli*.**

**Figure 1-figure supplement 4**

**An F66A substitution in cytTM-SpoIVFB partially overcomes inhibition by GFPΔ27BofA and SpoIVFA in *E. coli*.**

**Figure 2-figure supplement 1**

**Sequence alignments of BofA orthologs to determine conserved residues.**

**Figure 2-figure supplement 2**

**BofA C-terminal residues and residues preceding predicted TMS2 contribute to inhibition of SpoIVFB in *E. coli*.**

**Figure 3-figure supplement 1**

**GFPΔ27BofA variants are intact during *B. subtilis* sporulation.**

**Figure 4-figure supplement 1**

**Models of SpoIVFB and BofA TMS2.**

**Figure 4-figure supplement 2**

**Cys-less variants of SpoIVFA and MBPΔ27BofA inhibit cleavage of Pro-σ^K^(1-127) by cytTM-SpoIVFB in *E. coli*.**

**Figure 4-figure supplement 3**

**BofA TMS2 is in proximity to the active site of SpoIVFB.**

**Figure 4-figure supplement 4**

**Full-length BofA is in proximity to the active site of SpoIVFB.**

**Figure 4-figure supplement 5**

**BofA TMS2 has a preferred orientation in the active site cleft of SpoIVFB.**

**Figure 5-figure supplement 1**

**Neither GFPΔ27BofA nor full-length BofA when co-produced with SpoIVFA prevent Pro- σ^K^(1-127) from interacting with SpoIVFB.**

**Figure 5-figure supplement 2**

**GFPΔ27BofA and SpoIVFA do not prevent full-length Pro-σ^K^ from interacting with SpoIVFB.**

**Figure 5-figure supplement 3**

**Disulfide cross-linking between the cytTM-SpoIVFB CBS domain and the Pro-σ^K^(1-127) C-terminal region.**

**Figure 6-figure supplement 1**

**Disulfide cross-linking between E44C at the active site of cytTM-SpoIVFB and the Proregion of Pro-σ^K^(1-127) in the absence of inhibitory proteins or in the presence of MBPΔ27BofA and SpoIVFA.**

**Figure 6-figure supplement 2**

**Full-length BofA and SpoIVFA decrease cross-linking between E44C cytTM-SpoIVFB and V20C or K24C Pro-σ^K^(1-127).**

**Figure 6-figure supplement 3**

**Disulfide cross-linking between V70C in the cytTM-SpoIVFB membrane-reentrant loop and F18C or K24C in the Pro-σ^K^(1-127) N-terminal region is decreased more by full-length BofA than by MBPΔ27BofA (lacking TMS1).**

**Figure 6-figure supplement 4**

**Comparison of disulfide cross-linking between C46 in TMS2 of full-length BofA or MBPΔ27BofA (lacking TMS1) and E44C at or P135C near the active site of cytTM- SpoIVFB.**

**Figure 6-figure supplement 5**

**Disulfide cross-linking between Y214C in the cytTM-SpoIVFB interdomain linker and L41C in the Pro-σ^K^(1-127) N-terminal region is decreased more by full-length BofA than by MBPΔ27BofA (lacking TMS1).**

**Figure 7-figure supplement 1**

**Model of SpoIVFB tetramer with part of one Pro-σ^K^ molecule.**

**Source Data Files**

**Figure 1-source data 1**

**Immunoblot images (raw and annotated) and quantification of cleavage assays (Figure 1 B and C).**

**Figure 1-figure supplement 2-source data 1**

**Immunoblot images (raw and annotated) and quantification of cleavage assays.**

**Figure 1-figure supplement 3-source data 1**

**Immunoblot images (raw and annotated) and quantification of cleavage assays.**

**Figure 1-figure supplement 4-source data 1**

**Immunoblot images (raw and annotated) and quantification of cleavage assays.**

**Figure 2-source data 1**

**Immunoblot images (raw and annotated) and quantification of cleavage assays.**

**Figure 2-figure supplement 2-source data 1**

**Immunoblot images (raw and annotated) and quantification of cleavage assays.**

**Figure 3-source data 1**

**Immunoblot images (annotated) and quantification of cleavage assays (Figure 3A), and confocal microscopy images (see the readme file) and a table of the numbers of sporangia in which FS localization of GFPΔ27BofA and the three variants was counted (Figure 3B).**

**Figure 3-source data 2**

**Immunoblot images (raw) (Figure 3A).**

**Figure 3-figure supplement 1-source data 1 Immunoblot images (raw and annotated).**

**Figure 4-source data 1**

**Immunoblot images (raw and annotated).**

**Figure 4-figure supplement 1-source data 1 PyMOL session file used to produce the images.**

**Figure 4-figure supplement 2-source data 1**

**Immunoblot images (raw and annotated) and quantification of cleavage assays.**

**Figure 4-figure supplement 4-source data 1 Immunoblot images (raw and annotated).**

**Figure 4-figure supplement 5-source data 1**

**Immunoblot images (raw and annotated) (Figure 4-figure supplement 5 B and C) and quantification of cleavage assays (Figure 4-figure supplement 5B).**

**Figure 5-source data 1**

**Immunoblot images (raw and annotated) (Figure 5A) and quantification of cross-linking (Figure 5B).**

**Figure 5-figure supplement 1-source data 1 Immunoblot images (raw and annotated).**

**Figure 5-figure supplement 2-source data 1 Immunoblot images (raw and annotated).**

**Figure 5-figure supplement 3-source data 1**

**Immunoblot images (raw and annotated) and quantification of cleavage assays (Figure 5- figure supplement 3B).**

**Figure 6-source data 1**

**Quantification of cross-linking (Figure 6 A, C, and E-I).**

**Figure 6-figure supplement 1-source data 1**

**Immunoblot images (raw and annotated) (Figure 6-figure supplement 1A).**

**Figure 6-figure supplement 1-source data 2**

**Immunoblot images (raw and annotated) (Figure 6-figure supplement 1B).**

**Figure 6-figure supplement 2-source data 1**

**Immunoblot images (raw and annotated) (Figure 6-figure supplement 2A) and quantification of cross-linking (Figure 6-figure supplement 2B).**

**Figure 6-figure supplement 3-source data 1**

**Immunoblot images (raw and annotated) (Figure 6-figure supplement 3 A, C, E, and G) and quantification of cross-linking (Figure 6-figure supplement 3 B, D, and F).**

**Figure 6-figure supplement 4-source data 1**

**Immunoblot images (raw and annotated) (Figure 6-figure supplement 4 A, B, D, and E) and quantification of cross-linking (Figure 6-figure supplement 4 C and F).**

**Figure 6-figure supplement 5-source data 1 Immunoblot images (raw and annotated).**

**Figure 7-source data 1**

**PyMOL session file used to produce the images and PDB file of the model of a SpoIVFB tetramer with BofA and parts of SpoIVFA and Pro-σ^K^.**

**Figure 7-figure supplement 1-source data 1**

**PyMOL session file used to produce the images and PDB file of the model of a SpoIVFB tetramer with part of one molecule of Pro-σ^K^.**

**Supplementary Files Supplementary File 1**

**Plasmids, primers, and *B. subtilis* strains used in this study.**

**Supplementary File 2**

**Applied restraints from cross-linking, and distance evaluation, for the model of SpoIVFB in complex with part of Pro-σ^K^.**

**Supplementary File 3**

**Applied restraints from co-evolutionary coupling analysis, and distance evaluation, for the model of SpoIVFB in complex with part of Pro-σ^K^.**

**Supplementary File 4**

**Applied restraints from cross-linking, and distance evaluation, for the model of SpoIVFB in complex with BofA and parts of SpoIVFA and Pro-σ^K^.**

**Supplementary File 5**

**Applied restraints from co-evolutionary coupling analysis, and distance evaluation, for the model of SpoIVFB in complex with BofA and parts of SpoIVFA and Pro-σ^K^.**

## References

1. Brown MS, Ye J, Rawson RB, & Goldstein JL (2000) Regulated intramembrane proteolysis: a control mechanism conserved from bacteria to humans. Cell 100(4):391–398.

2. Urban S (2013) Mechanisms and cellular functions of intramembrane proteases. Biochim. Biophys. Acta - Biomembr. 1828:2797–2800.

3. Kuhnle N, Dederer V, & Lemberg MK (2019) Intramembrane proteolysis at a glance: from signalling to protein degradation. J. Cell Sci. 132(16):jcs217745.

4. Manolaridis I, et al. (2013) Mechanism of farnesylated CAAX protein processing by the intramembrane protease Rce1. Nature 504(7479):301–305.

5. Hu J, Xue Y, Lee S, & Ha Y (2011) The crystal structure of GXGD membrane protease FlaK. Nature 475(7357):528–531.

6. Li X, et al. (2013) Structure of a presenilin family intramembrane aspartate protease. Nature 493(7430):56–61.

7. Feng L, et al. (2007) Structure of a site-2 protease family intramembrane metalloprotease. Science 318(5856):1608–1612.

8. Wang Y, Zhang Y, & Ha Y (2006) Crystal structure of a rhomboid family intramembrane protease. Nature 444(7116):179–183.

9. Wu Z, et al. (2006) Structural analysis of a rhomboid family intramembrane protease reveals a gating mechanism for substrate entry. Nat. Struct. Mol. Biol. 13(12):1084–1091.

10. Cho S, Dickey SW, & Urban S (2016) Crystal structures and inhibition kinetics reveal a two-stage catalytic mechanism with drug design implications for rhomboid proteolysis. Mol. Cell 61(3):329–340.

11. Cho S, Baker RP, Ji M, & Urban S (2019) Ten catalytic snapshots of rhomboid intramembrane proteolysis from gate opening to peptide release. Nat. Struct. Mol. Biol. 26(10):910–918.

12. Ticha A, et al. (2017) General and modular strategy for designing potent, selective, and pharmacologically compliant inhibitors of rhomboid proteases. Cell Chem. Biol. 24(12):1523–1536 e1524.

13. Yang G, et al. (2019) Structural basis of Notch recognition by human γ-secretase. Nature 565(7738):192–197.

14. Zhou R, et al. (2019) Recognition of the amyloid precursor protein by human γ-secretase. Science 363(6428):eaaw0930.

15. Rawson RB (2013) The site-2 protease. Biochim. Biophys. Acta - Biomembr. 1828(12):2801–2807.

16. Ye J (2013) Roles of regulated intramembrane proteolysis in virus infection and antiviral immunity. Biochim. Biophys. Acta - Biomembr. 1828(12):2926–2932.

17. Adam Z (2013) Emerging roles for diverse intramembrane proteases in plant biology. Biochim. Biophys. Acta - Biomembr. 1828(12):2933–2936.

18. Chang YC, Bien CM, Lee H, Espenshade PJ, & Kwon-Chung KJ (2007) Sre1p, a regulator of oxygen sensing and sterol homeostasis, is required for virulence in *Cryptococcus neoformans*. Mol. Microbiol. 64(3):614–629.

19. Urban S (2009) Making the cut: central roles of intramembrane proteolysis in pathogenic microorganisms. Nat. Rev. Microbiol. 7:411–423.

20. Kroos L & Akiyama Y (2013) Biochemical and structural insights into intramembrane metalloprotease mechanisms. Biochim. Biophys. Acta - Biomembr. 1828(12):2873–2885.

21. Schneider JS & Glickman MS (2013) Function of site-2 proteases in bacteria and bacterial pathogens. Biochim. Biophys. Acta - Biomembr. 1828(12):2808–2814.

22. de Hoon MJ, Eichenberger P, & Vitkup D (2010) Hierarchical evolution of the bacterial sporulation network. Curr. Biol. 20(17):R735–R745.

23. Galperin MY, et al. (2012) Genomic determinants of sporulation in *Bacilli* and *Clostridia*: towards the minimal set of sporulation-specific genes. Environ. Microbiol. 14(11):2870–2890.

24. Abecasis AB, et al. (2013) A genomic signature and the identification of new sporulation genes. J. Bacteriol. 195(9):2101–2115.

25. Ramos-Silva P, Serrano M, & Henriques AO (2019) From root to tips: sporulation evolution and specialization in *Bacillus subtilis* and the intestinal pathogen *Clostridioides difficile*. Mol. Biol. Evol. 36(12):2714–2736.

26. McKenney PT, Driks A, & Eichenberger P (2013) The *Bacillus subtilis* endospore: assembly and functions of the multilayered coat. Nat. Rev. Microbiol. 11(1):33–44.

27. Al-Hinai MA, Jones SW, & Papoutsakis ET (2015) The *Clostridium* sporulation programs: diversity and preservation of endospore differentiation. Microbiol. Mol. Biol. Rev. 79(1):19–37.

28. Checinska A, Paszczynski A, & Burbank M (2015) *Bacillus* and other spore-forming genera: variations in responses and mechanisms for survival. Annu. Rev. Food Sci. Technol. 6:351–369.

29. Browne HP, et al. (2016) Culturing of ’unculturable’ human microbiota reveals novel taxa and extensive sporulation. Nature 533(7604):543–546.

30. Tan IS & Ramamurthi KS (2014) Spore formation in *Bacillus subtilis*. Environ. Microbiol. Rep. 6(3):212–225.

31. Cutting S, et al. (1990) A forespore checkpoint for mother-cell gene expression during development in *Bacillus subtilis*. Cell 62:239–250.

32. Cutting S, Roels S, & Losick R (1991) Sporulation operon *spoIVF* and the characterization of mutations that uncouple mother-cell from forespore gene expression in *Bacillus subtilis*. J. Mol. Biol. 221:1237–1256.

33. Ricca E, Cutting S, & Losick R (1992) Characterization of *bofA*, a gene involved in intercompartmental regulation of pro-σ^K^ processing during sporulation in *Bacillus subtilis*. J. Bacteriol. 174:3177–3184.

34. Resnekov O, Alper S, & Losick R (1996) Subcellular localization of proteins governing the proteolytic activation of a developmental transcription factor in *Bacillus subtilis*. Genes Cells 1(6):529–542.

35. Rudner DZ, Pan Q, & Losick RM (2002) Evidence that subcellular localization of a bacterial membrane protein is achieved by diffusion and capture. Proc. Natl. Acad. Sci. USA 99(13):8701–8706.

36. Rudner DZ & Losick R (2002) A sporulation membrane protein tethers the pro-σ^K^ processing enzyme to its inhibitor and dictates its subcellular localization. Genes Dev. 16(8):1007–1018.

37. Zhou R & Kroos L (2004) BofA protein inhibits intramembrane proteolysis of pro-σ^K^ in an intercompartmental signaling pathway during *Bacillus subtilis* sporulation. Proc. Natl. Acad. Sci. USA 101(17):6385–6390.

38. Kroos L, Yu YT, Mills D, & Ferguson-Miller S (2002) Forespore signaling is necessary for pro-σ^K^ processing during *Bacillus subtilis* sporulation despite the loss of SpoIVFA upon translational arrest. J. Bacteriol. 184(19):5393–5401.

39. Cutting S, Driks A, Schmidt R, Kunkel B, & Losick R (1991) Forespore-specific transcription of a gene in the signal transduction pathway that governs pro-σ^K^ processing in *Bacillus subtilis*. Genes Dev. 5:456–466.

40. Pan Q, Losick R, & Rudner DZ (2003) A second PDZ-containing serine protease contributes to activation of the sporulation transcription factor σ^K^ in *Bacillus subtilis*. J. Bacteriol. 185(20):6051–6056.

41. Dong TC & Cutting SM (2003) SpoIVB-mediated cleavage of SpoIVFA could provide the intercellular signal to activate processing of Pro-σ^K^ in *Bacillus subtilis*. Mol. Microbiol. 49(5):1425–1434.

42. Campo N & Rudner DZ (2006) A branched pathway governing the activation of a developmental transcription factor by regulated intramembrane proteolysis. Mol. Cell 23(1):25–35.

43. Campo N & Rudner DZ (2007) SpoIVB and CtpB are both forespore signals in the activation of the sporulation transcription factor σ^K^ in *Bacillus subtilis*. J. Bacteriol. 189(16):6021–6027.

44. Mastny M, et al. (2013) CtpB assembles a gated protease tunnel regulating cell-cell signaling during spore formation in *Bacillus subtilis*. Cell 155(3):647–658.

45. Zhou R & Kroos L (2005) Serine proteases from two cell types target different components of a complex that governs regulated intramembrane proteolysis of pro-σ^K^ during *Bacillus subtilis* development. Mol. Microbiol. 58(3):835–846.

46. Rudner D, Fawcett P, & Losick R (1999) A family of membrane-embedded metalloproteases involved in regulated proteolysis of membrane-associated transcription factors. Proc. Natl. Acad. Sci. USA 96:14765–14770.

47. Yu Y-TN & Kroos L (2000) Evidence that SpoIVFB is a novel type of membrane metalloprotease governing intercompartmental communication during *Bacillus subtilis* sporulation. J. Bacteriol. 182:3305–3309.

48. Zhou R, Cusumano C, Sui D, Garavito RM, & Kroos L (2009) Intramembrane proteolytic cleavage of a membrane-tethered transcription factor by a metalloprotease depends on ATP. Proc. Natl. Acad. Sci. USA 106:16174–16179.

49. Kroos L, Kunkel B, & Losick R (1989) Switch protein alters specificity of RNA polymerase containing a compartment-specific sigma factor. Science 243:526–529.

50. Eichenberger P, et al. (2004) The program of gene transcription for a single differentiating cell type during sporulation in *Bacillus subtilis*. PLoS Biol. 2(10):e328.

51. Morohashi Y & Tomita T (2013) Protein trafficking and maturation regulate intramembrane proteolysis. Biochim. Biophys. Acta 1828(12):2855–2861.

52. Lichtenthaler SF, Lemberg MK, & Fluhrer R (2018) Proteolytic ectodomain shedding of membrane proteins in mammals-hardware, concepts, and recent developments. EMBO J. 37(15):e99456.

53. Beard HA, Barniol-Xicota M, Yang J, & Verhelst SHL (2019) Discovery of cellular roles of intramembrane proteases. ACS Chem. Biol. 14(11):2372–2388.

54. Kinch LN, Ginalski K, & Grishin NV (2006) Site-2 protease regulated intramembrane proteolysis: sequence homologs suggest an ancient signaling cascade. Protein Sci. 15(1):84–93.

55. Zhang Y, Luethy PM, Zhou R, & Kroos L (2013) Residues in conserved loops of intramembrane metalloprotease SpoIVFB interact with residues near the cleavage site in Pro-σ^K^. J. Bacteriol. 195(21):4936–4946.

56. Halder S, Parrell D, Whitten D, Feig M, & Kroos L (2017) Interaction of intramembrane metalloprotease SpoIVFB with substrate Pro-σ^K^. Proc. Natl. Acad. Sci. USA 114:E10677–E10686.

57. Saribas AS, Gruenke L, & Waskell L (2001) Overexpression and purification of the membrane-bound cytochrome P450 2B4. Protein Expr. Purif. 21(2):303–309.

58. Prince H, Zhou R, & Kroos L (2005) Substrate requirements for regulated intramembrane proteolysis of *Bacillus subtilis* pro-σ^K^. J. Bacteriol. 187:961–971.

59. Ramirez-Guadiana FH, et al. (2018) Evidence that regulation of intramembrane proteolysis is mediated by substrate gating during sporulation in *Bacillus subtilis*. PLoS Genet. 14(11):e1007753.

60. Varcamonti M, Marasco R, De Felice M, & Sacco M (1997) Membrane topology analysis of the *Bacillus subtilis* BofA protein involved in pro-σ^K^ processing. Microbiol. 143(Pt 4):1053–1058.

61. Resnekov O (1999) Role of the sporulation protein BofA in regulating activation of the *Bacillus subtilis* developmental transcription factor σ^K^. J. Bacteriol. 181:5384–5388.

62. Arolas JL, Goulas T, Cuppari A, & Gomis-Ruth FX (2018) Multiple architectures and mechanisms of latency in metallopeptidase zymogens. Chem. Rev. 118(11):5581–5597.

63. Van Wart HE & Birkedal-Hansen H (1990) The cysteine switch: a principle of regulation of metalloproteinase activity with potential applicability to the entire matrix metalloproteinase gene family. Proc. Natl. Acad. Sci. USA 87(14):5578–5582.

64. Odintsov SG, Sabala I, Marcyjaniak M, & Bochtler M (2004) Latent LytM at 1.3 A resolution. J. Mol. Biol. 335(3):775–785.

65. Zhang Y, et al. (2016) Complex formed between intramembrane metalloprotease SpoIVFB and its substrate, Pro-σ^K^. J. Biol. Chem. 291:10347–10362.

66. Akiyama K, et al. (2015) Roles of the membrane-reentrant β-hairpin-like loop of RseP protease in selective substrate cleavage. eLife 4:e08928

67. Rawson RB (2003) The SREBP pathway--insights from Insigs and insects. Nat. Rev. Mol. Cell Biol. 4(8):631–640.

68. Lima S, Guo MS, Chaba R, Gross CA, & Sauer RT (2013) Dual molecular signals mediate the bacterial response to outer-membrane stress. Science 340(6134):837–841.

69. Hizukuri Y, et al. (2014) A structure-based model of substrate discrimination by a noncanonical PDZ tandem in the intramembrane-cleaving protease RseP. Structure 22(2):326–336.

70. Bolduc DM, Montagna DR, Gu YL, Selkoe DJ, & Wolfe MS (2016) Nicastrin functions to sterically hinder γ-secretase-substrate interactions driven by substrate transmembrane domain. Proc. Natl. Acad. Sci. USA 113(5):E509–E518.

71. Fukumori A & Steiner H (2016) Substrate recruitment of γ-secretase and mechanism of clinical presenilin mutations revealed by photoaffinity mapping. EMBO J. 35(15):1628–1643.

72. Pereira FC, et al. (2013) The spore differentiation pathway in the enteric pathogen *Clostridium difficile*. PLoS Genet. 9(10):e1003782.

73. Saujet L, et al. (2013) Genome-wide analysis of cell type-specific gene transcription during spore formation in *Clostridium difficile*. PLoS Genet. 9(10):e1003756.

74. Haraldsen JD & Sonenshein AL (2003) Efficient sporulation in *Clostridium difficile* requires disruption of the σ^K^ gene. Mol. Microbiol. 48(3):811–821.

75. Ticha A, Collis B, & Strisovsky K (2018) The rhomboid superfamily: structural mechanisms and chemical biology opportunities. Trends Biochem. Sci. 43(9):726–739.

76. Wolfe MS (2019) Dysfunctional γ-secretase in familial Alzheimer’s disease. Neurochem Res 44(1):5–11.

77. De Strooper B & Chavez Gutierrez L (2015) Learning by failing: ideas and concepts to tackle γ-secretases in Alzheimer’s disease and beyond. Annu. Rev. Pharmacol. Toxicol. 55:419–437.

78. Szaruga M, et al. (2017) Alzheimer’s-causing mutations shift Aβ length by destabilizing γ-secretase-Aβn interactions. Cell 170(3):443–456 e414.

79. Konovalova A, et al. (2018) Inhibitor of intramembrane protease RseP blocks the σ^E^ response causing lethal accumulation of unfolded outer membrane proteins. Proc. Natl. Acad. Sci. USA 115(28):E6614–E6621.

80. Jacobsen JA, Major Jourden JL, Miller MT, & Cohen SM (2010) To bind zinc or not to bind zinc: an examination of innovative approaches to improved metalloproteinase inhibition. Biochim. Biophys. Acta 1803(1):72–94.

81. Fields GB (2015) New strategies for targeting matrix metalloproteinases. Matrix Biol. 44–46:239-246.

82. Gomis-Ruth FX (2017) Third time lucky? Getting a grip on matrix metalloproteinases. J. Biol. Chem. 292(43):17975–17976.

83. Moss ML, et al. (2007) The ADAM10 prodomain is a specific inhibitor of ADAM10 proteolytic activity and inhibits cellular shedding events. J. Biol. Chem. 282(49):35712–35721.

84. Wong E, et al. (2016) Harnessing the natural inhibitory domain to control TNFα Converting Enzyme (TACE) activity *in vivo*. Sci. Rep. 6:35598.

85. Zorzi A, Deyle K, & Heinis C (2017) Cyclic peptide therapeutics: past, present and future. Curr. Opin. Chem. Biol. 38:24–29.

86. Sohrabi C, Foster A, & Tavassoli A (2020) Methods for generating and screening libraries of genetically encoded cyclic peptides in drug discovery. Nat. Rev. Chem. 4(2):90–101.

87. Hanahan D (1983) Studies on transformation of *Escherichia coli* with plasmids. J. Mol. Biol. 166:557–580.

88. Lu S, Halberg R, & Kroos L (1990) Processing of the mother-cell σ factor, σ^K^, may depend on events occurring in the forespore during *Bacillus subtilis* development. Proc. Natl. Acad. Sci. USA 87:9722–9726.

89. Notredame C, Higgins DG, & Heringa J (2000) T-Coffee: A novel method for fast and accurate multiple sequence alignment. J. Mol. Biol. 302(1):205–217.

90. Harwood CR & Cutting SM (1990) Molecular Biological Methods for Bacillus (John Wiley & Sons, Chichester, England) p 581.

91. Parrell D & Kroos L (2020) Channels modestly impact compartment-specific ATP levels during *Bacillus subtilis* sporulation and a rise in the mother cell ATP level is not necessary for Pro-σ^K^ cleavage. Mol. Microbiol. 114:563–581.

92. Koide K, Ito K, & Akiyama Y (2008) Substrate recognition and binding by RseP, an *Escherichia coli* intramembrane protease. J. Biol. Chem. 283(15):9562–9570.

93. Miller MD, et al. (2004) Crystal structure of a tandem cystathionine-beta-synthase (CBS) domain protein (TM0935) from *Thermotoga maritima* at 1.87 Angstrom resolution. Proteins 57(1):213–217.

94. Schrecker M, Korobenko J, & Hite RK (2020) Cryo-EM structure of the lysosomal chloride-proton exchanger CLC-7 in complex with OSTM1. eLife 9:e59555.

95. Feklistov A & Darst SA (2011) Structural basis for promoter -10 element recognition by the bacterial RNA polymerase sigma subunit. Cell 147(6):1257–1269.

96. Yang J, et al. (2020) Improved protein structure prediction using predicted interresidue orientations. Proc. Natl. Acad. Sci. USA 117(3):1496–1503.

97. Remmert M, Biegert A, Hauser A, & Soding J (2011) HHblits: lightning-fast iterative protein sequence searching by HMM-HMM alignment. Nat. Methods 9(2):173–175.

98. Mirdita M, et al. (2017) Uniclust databases of clustered and deeply annotated protein sequences and alignments. Nucleic Acids Res. 45(D1):D170–D176.

99. Chaudhury S, Lyskov S, & Gray JJ (2010) PyRosetta: a script-based interface for implementing molecular modeling algorithms using Rosetta. Bioinformatics 26(5):689–691.

100. Park H, et al. (2016) Simultaneous optimization of biomolecular energy functions on features from small molecules and macromolecules. J. Chem. Theory Comput. 12(12):6201–6212.

101. Ha Y (2009) Structure and mechanism of intramembrane protease. Semin. Cell Dev. Biol. 20(2):240–250.

102. Zelenski N, Rawson R, Brown M, & Goldstein J (1999) Membrane topology of S2P, a protein required for intramembranous cleavage of sterol regulatory element-binding proteins. J. Biol. Chem. 274:21973–21980.

103. Nohturfft A, DeBose-Boyd RA, Scheek S, Goldstein JL, & Brown MS (1999) Sterols regulate cycling of SREBP cleavage-activating protein (SCAP) between endoplasmic reticulum and Golgi. Proc. Natl. Acad. Sci. USA 96(20):11235–11240.

104. DeBose-Boyd RA, et al. (1999) Transport-dependent proteolysis of SREBP: relocation of site-1 protease from Golgi to ER obviates the need for SREBP transport to Golgi. Cell 99(7):703–712.

105. Chen G & Zhang X (2010) New insights into S2P signaling cascades: regulation, variation, and conservation. Protein Sci. 19(11):2015–2030.

106. Sakai J, et al. (1996) Sterol-regulated release of SREBP-2 from cell membrane requires two sequential cleavages, one within a transmembrane domain. Cell 85:1037–1048.

107. Sakai J, et al. (1997) Identification of complexes between the COOH-terminal domains of sterol regulatory element-binding proteins (SREBPs) and SREBP cleavage-activating protein. J. Biol. Chem. 272(32):20213–20221.

108. Horton JD, et al. (2003) Combined analysis of oligonucleotide microarray data from transgenic and knockout mice identifies direct SREBP target genes. Proc. Natl. Acad. Sci. USA 100(21):12027–12032.

109. Kanehara K, Akiyama Y, & Ito K (2001) Characterization of the *yaeL* gene product and its S2P-protease motifs in *Escherichia coli*. Gene 281(1-2):71–79.

110. Drew D, et al. (2002) Rapid topology mapping of *Escherichia coli* inner-membrane proteins by prediction and PhoA/GFP fusion analysis. Proc. Natl. Acad. Sci. USA 99(5):2690–2695.

111. Ades SE, Connolly LE, Alba BM, & Gross CA (1999) The *Escherichia coli* σ^E^- dependent extracytoplasmic stress response is controlled by the regulated proteolysis of an anti-σ factor. Genes Dev. 13(18):2449–2461.

112. Walsh NP, Alba BM, Bose B, Gross CA, & Sauer RT (2003) OMP peptide signals initiate the envelope-stress response by activating DegS protease via relief of inhibition mediated by its PDZ domain. Cell 113(1):61–71.

113. Kanehara K, Ito K, & Akiyama Y (2002) YaeL (EcfE) activates the σ^E^ pathway of stress response through a site-2 cleavage of anti-σ^E^, RseA. Genes Dev. 16(16):2147–2155.

114. Alba BM, Leeds JA, Onufryk C, Lu CZ, & Gross CA (2002) DegS and YaeL participate sequentially in the cleavage of RseA to activate the σ^E^-dependent extracytoplasmic stress response. Genes Dev. 16(16):2156–2168.

115. Flynn JM, Neher SB, Kim YI, Sauer RT, & Baker TA (2003) Proteomic discovery of cellular substrates of the ClpXP protease reveals five classes of ClpX-recognition signals. Mol. Cell 11(3):671–683.

116. Chaba R, Grigorova IL, Flynn JM, Baker TA, & Gross CA (2007) Design principles of the proteolytic cascade governing the σ^E^-mediated envelope stress response in *Escherichia coli*: keys to graded, buffered, and rapid signal transduction. Genes Dev. 21(1):124–136.

