## Supplementary material for "Inhibitory proteins block substrate access by occupying the active site cleft of *Bacillus subtilis* intramembrane protease SpoIVFB": Figure supplements

**Figure 1-figure supplement 1**

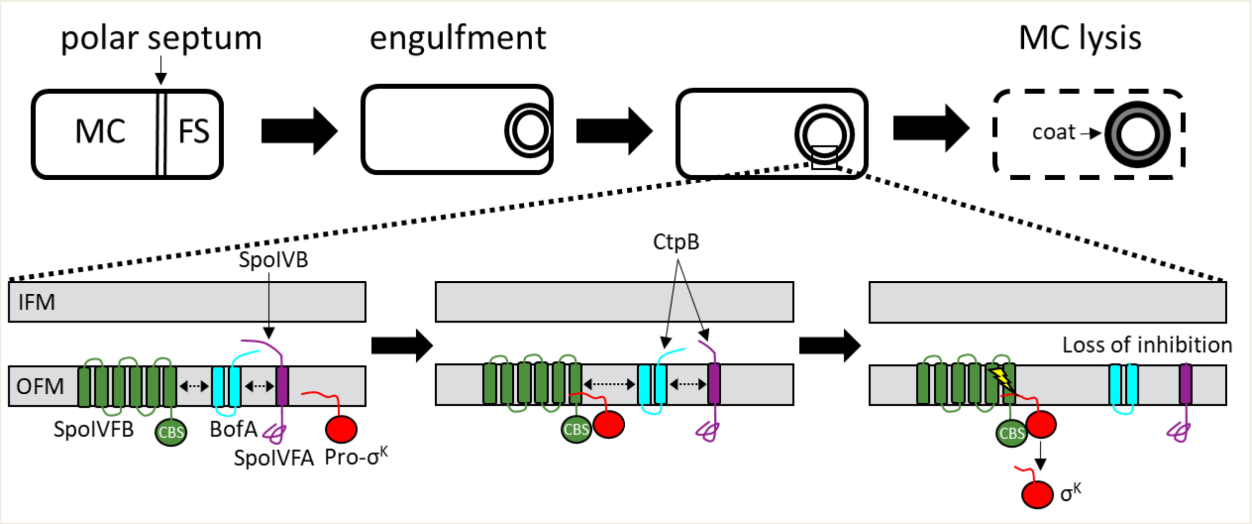


**Morphological changes during endosporulation and regulated intramembrane proteolysis of Pro-$\sigma^K$ in *B. subtilis*.**

Upon starvation, a polar septum forms that divides the cell into MC (MC) and FS (FS) compartments (*Top*). The MC engulfs the FS, resulting in two membranes surrounding the FS. Upon completion of engulfment, regulated intramembrane proteolysis releases $\sigma^K$ into the MC (*Bottom*), where it directs RNA polymerase to transcribe genes whose products form the spore coat and cause MC lysis. The proteolytic cascade begins with SpoIVB, which is exported from the FS into the intermembrane space between the inner FS membrane (IFM) and the outer FS membrane (OFM). SpoIVB cleaves the C-terminal region of SpoIVFA. A second protease, CtpB, is also exported from the FS, and further cleaves the C-terminal region of SpoIVFA, and can cleave the C-terminal end of BofA. Inhibition of SpoIVFB is lost, allowing it to cleave Pro-$\sigma^K$ (lightning bolt), releasing $\sigma^K$ into the MC. The dashed double-headed arrows indicate that SpoIVFB, BofA, and SpoIVFA initially form a complex. See the text for references. Neither the structure of the complex nor how it changes during the proteolytic cascade are fully known.

**Figure 1-figure supplement 2**

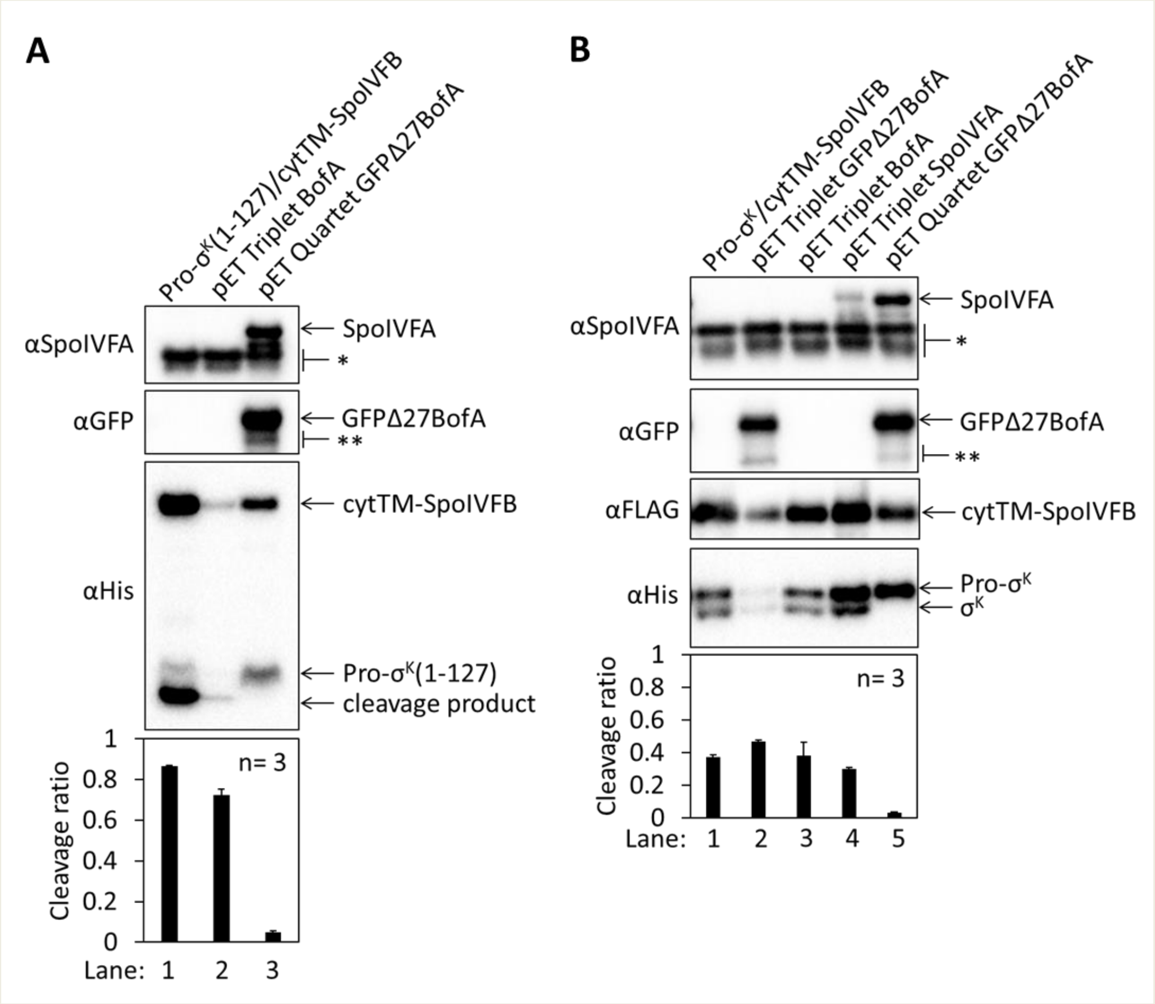


**Full-length BofA alone fails to inhibit Pro-$\sigma^K$(1-127) cleavage in *E. coli* and full-length Pro-$\sigma^K$ is similar to Pro-$\sigma^K$(1-127) in terms of requirements for cleavage inhibition.**

(**A**) Cleavage assays examining inhibition by full-length BofA alone. A pET Triplet plasmid was used to produce Pro-$\sigma^K$(1-127), cytTM-SpoIVFB, and BofA (lane 2, pSO312), and compared with a pET Duet plasmid that produces only Pro-$\sigma^K$(1-127) and cytTM-SpoIVFB (lane 1, pYZ2) or with a pET Quartet plasmid that produces Pro-$\sigma^K$(1-127), cytTM-SpoIVFB, SpoIVFA, and GFPΔ27BofA (lane 3, pSO40). Samples collected after 2 h of IPTG induction were subjected to immunoblot analysis, and the graph shows quantification of the cleavage ratio, as explained in the Figure 1B legend. (**B**) Cleavage assays with full-length Pro-$\sigma^K$ as substrate. Pro-$\sigma^K$ and cytTM-SpoIVFB were produced alone (lane 1, pSO290) or with GFPΔ27BofA (lane 2, pSO313), full-length BofA (lane 3, pSO314), SpoIVFA (lane 4, pSO315), or both GFPΔ27BofA and SpoIVFA (lane 5, pSO289) in *E. coli*. Samples collected after 2 h of IPTG induction were subjected to immunoblot analysis with SpoIVFA, GFP, FLAG, or penta-His antibodies as indicated. Quantification was as in (**A**).

**Figure 1-figure supplement 2-source data 1**
**Immunoblot images (raw and annotated) and quantification of cleavage assays.**

**Figure 1-figure supplement 3**

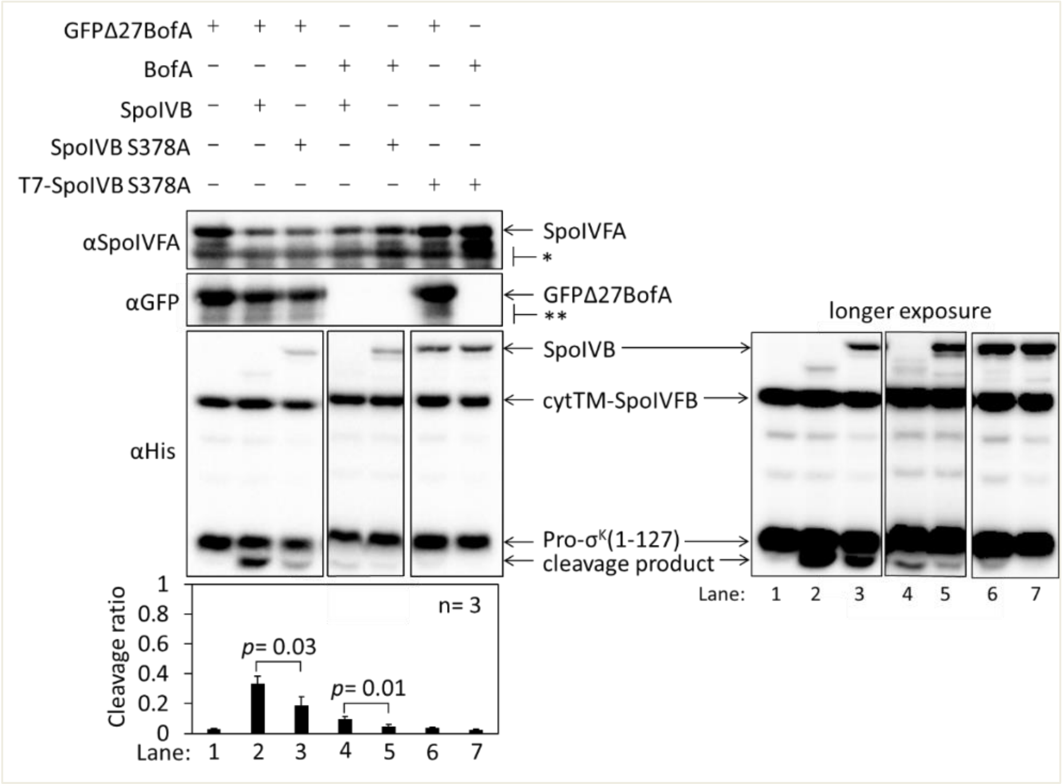


**SpoIVB partially relieves inhibition of SpoIVFB by BofA and SpoIVFA in *E. coli*.**
Cleavage assays examining the effects of SpoIVB production on BofA and SpoIVFA inhibition of SpoIVFB in *E. coli*. pET Quintet plasmids were used to produce Pro-$\sigma^K$(1-127), cytTM-SpoIVFB, SpoIVFA, GFPΔ27BofA or full-length BofA, and SpoIVB or SpoIVB S378A from pSO240, pSO241 and pSO251-pSO254. For comparison, a pET Quartet plasmid was used to produce Pro-$\sigma^K$(1-127), cytTM-SpoIVFB, SpoIVFA, and GFPΔ27BofA (lane 1, pSO40). Samples collected after 2 h of IPTG induction were subjected to immunoblot analysis, and the graph shows quantification of the cleavage ratio, as explained in the Figure 1B legend. For the penta-His antibodies, 2 and 30 s exposures are shown. Student's two-tailed *t* tests were performed to compare certain cleavage ratios (*p* values are indicated). With pET Quintet plasmids that produced active SpoIVB and either GFPΔ27BofA or BofA (lanes 2 & 4), the Pro-$\sigma^K$(1-127) cleavage ratios were greater than with corresponding pET Quintet plasmids that produced inactive SpoIVB S378A (lanes 3 & 5), indicating partial relief from inhibition of SpoIVFB. When *spoIVB* was expressed under the same T7 promoter as inhibitory proteins (i.e., *gfpΔ27bofA/spoIVFA* or *bofA/spoIVFA*), accumulation of GFPΔ27BofA and SpoIVFA was consistently less (lanes 2-5) compared to pET Quartet (lane 1), which exhibited a very low cleavage ratio, as expected. To try to increase protein production, we attempted to engineer pET Quintet plasmids with *spoIVB* under control of an additional T7 promoter. Samples with *spoIVB S378A* under control of an additional T7 promoter (T7-SpoIVB S378A) (lanes 6 & 7) accumulated more SpoIVB S378A (compared with lanes 3 & 5), and more GFPΔ27BofA and SpoIVFA (similar to that observed for pET Quartet in lane 1), and exhibited a very low cleavage ratio (similar to lane 1). Attempts at engineering pET Quintet plasmids to express active *spoIVB* from an additional T7 promoter were unsuccessful. Cell lysis occurred in overnight cultures of strains containing such plasmids, suggesting that T7-SpoIVB is toxic to *E. coli*.

**Figure 1-figure supplement 3-source data 1**
**Immunoblot images (raw and annotated) and quantification of cleavage assays.**

**Figure 1-figure supplement 4**

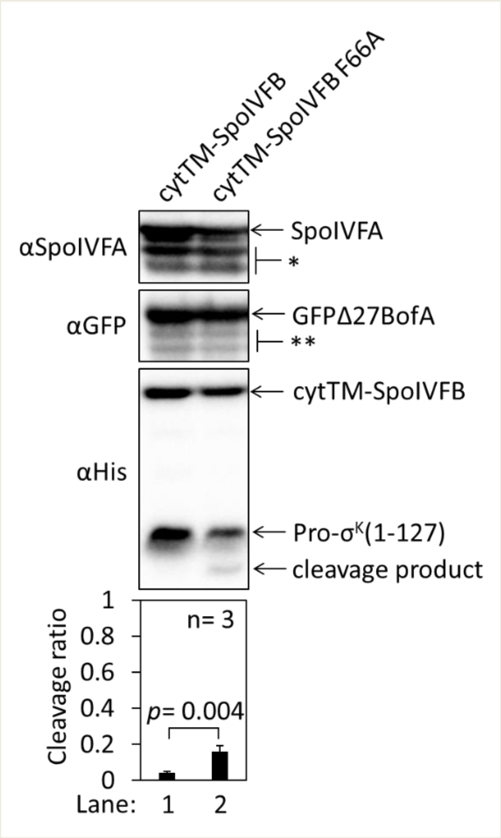


**An F66A substitution in cytTM-SpoIVFB partially overcomes inhibition by GFPΔ27BofA and SpoIVFA in *E. coli*.**

The phenylalanine residue at position 66 of SpoIVFB was predicted to help stabilize a closed conformation that would prevent Pro-$\sigma^K$ access to the active site, and BofA and SpoIVFA were envisioned to further stabilize SpoIVFB in a closed conformation (59). SpoIVB-dependent cleavage of SpoIVFA would presumably favor an open conformation of SpoIVFB capable of cleaving Pro-$\sigma^K$. In the absence of SpoIVB, it was found that an F66A substitution in SpoIVFB-YFP allowed Pro-$\sigma^K$ cleavage during *B. subtilis* sporulation, albeit with reduced efficiency (59). To test whether SpoIVFB inhibition could be relieved by the F66A substitution in *E. coli*, pET Quartet plasmids were used to produce Pro-$\sigma^K$(1-127), SpoIVFA, GFPΔ27BofA, and cytTM-SpoIVFB from pSO40 as a control (lane 1) or cytTM-SpoIVFB F66A from pSO193 (lane 2). Samples collected after 2 h of IPTG induction were subjected to immunoblot analysis, and the graph shows quantification of the cleavage ratio, as explained in the Figure 1B legend. A Student's two-tailed *t* test was performed to compare the cleavage ratios (*p* value is indicated). The Pro-$\sigma^K$(1-127) cleavage ratio was greater when cytTM-SpoIVFB F66A was produced (lane 2) than when cytTM-SpoIVFB was produced (lane 1), but not as great as in the absence of inhibitory proteins (Figure 1B, lane 1), indicating that cytTM-SpoIVFB F66A partially overcomes inhibition by GFPΔ27BofA and SpoIVFA in *E. coli*.

**Figure 1-figure supplement 4-source data 1**
**Immunoblot images (raw and annotated) and quantification of cleavage assays.**

### Figure 2-figure supplement 1

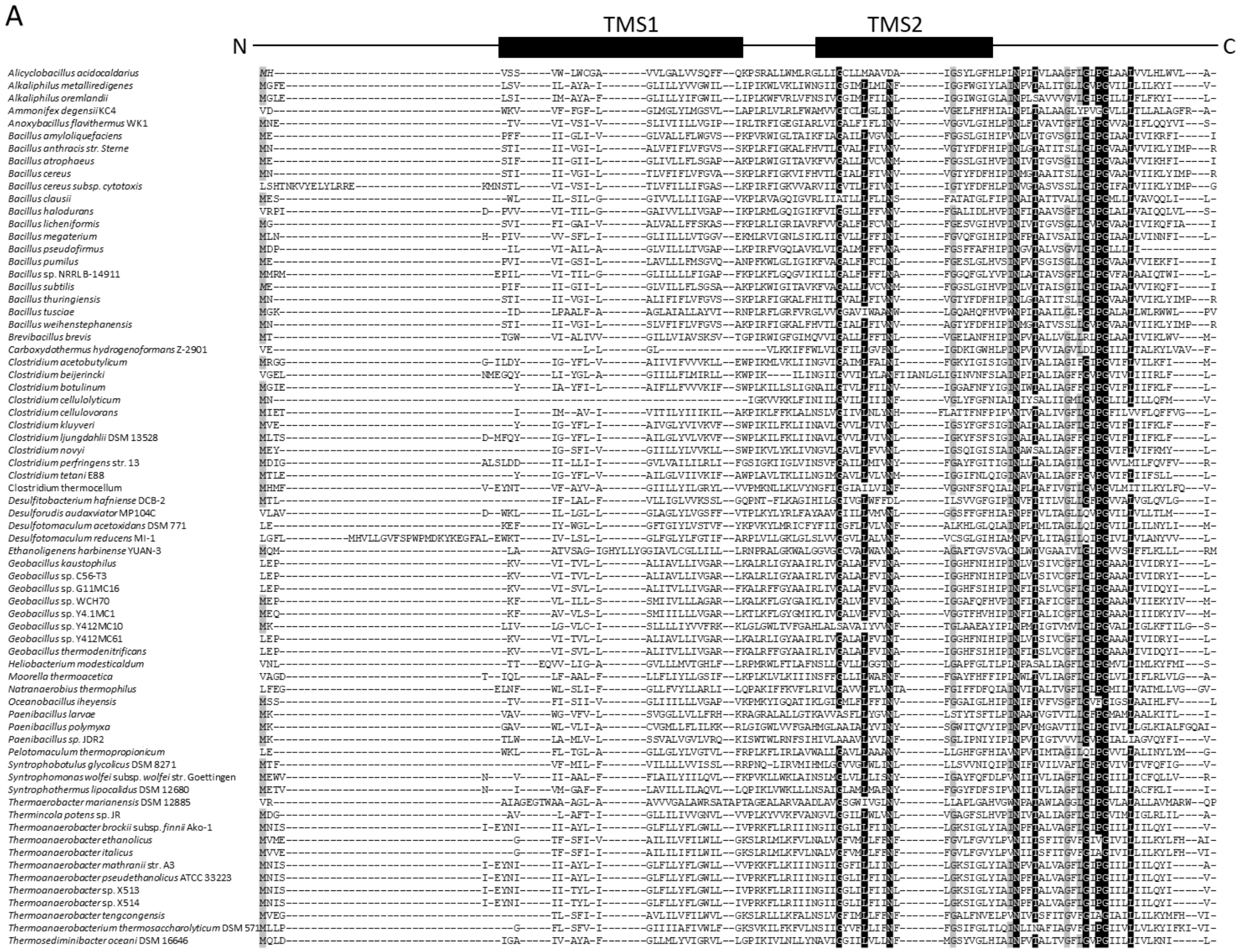

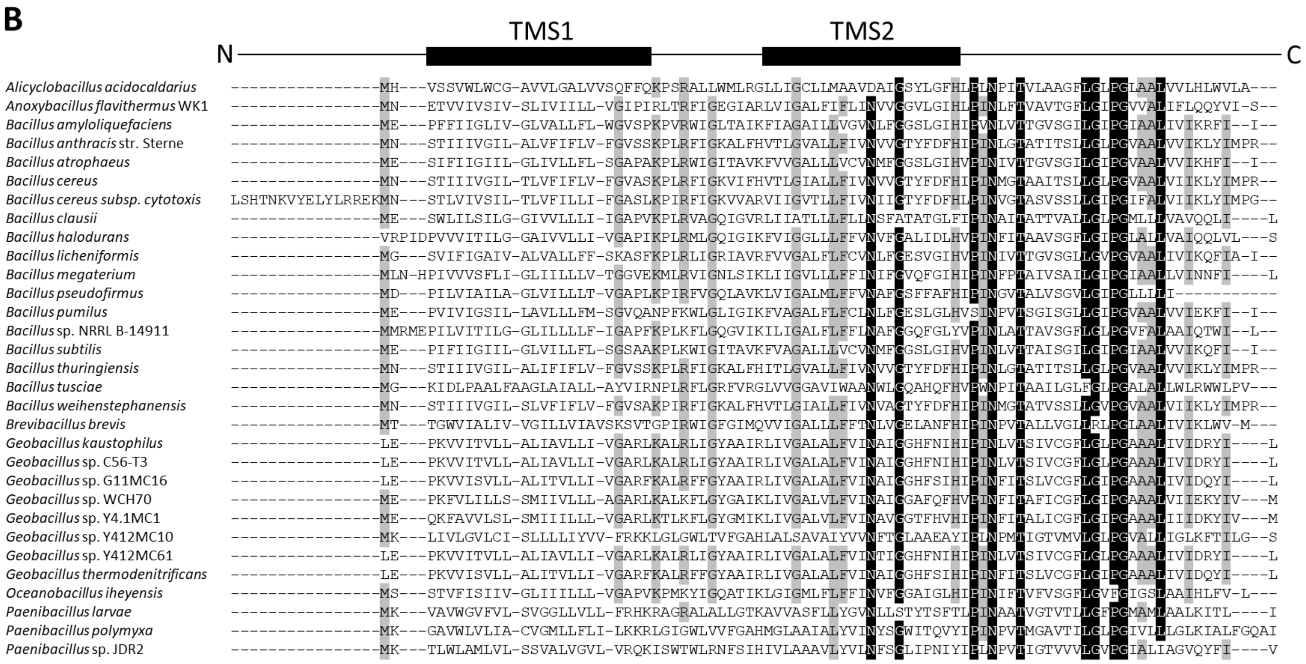


**Sequence alignments of BofA orthologs to determine conserved residues.**
(**A**) Sequence alignment of *B. subtilis* BofA with 69 orthologs. Five residues (gray: M1, G51, I60, G69, L71) are at least 70% conserved and nine residues (black: G40, L44, N48, N61, T64, G72, P74, G75, L79) are at least 90% conserved, all within transmembrane segment 2 (TMS2) and the C-terminal region, except M1. (**B**) Sequence alignment of *B. subtilis* BofA with 30 orthologs that contain *spoIVFA* in their genome. In comparison to *A*, four additional residues in predicted TMS2 and the C-terminal region were deemed of interest for Ala substitutions (H57, I82, and I86 are at least 70% conserved, and P59 is at least 90% conserved). A77 and A78 are also at least 70% conserved, but ineligible for Ala substitutions. V45 of *B. subtilis* does not match F, which is at least 70% conserved at the corresponding position of orthologs.

**Figure 2-figure supplement 2**

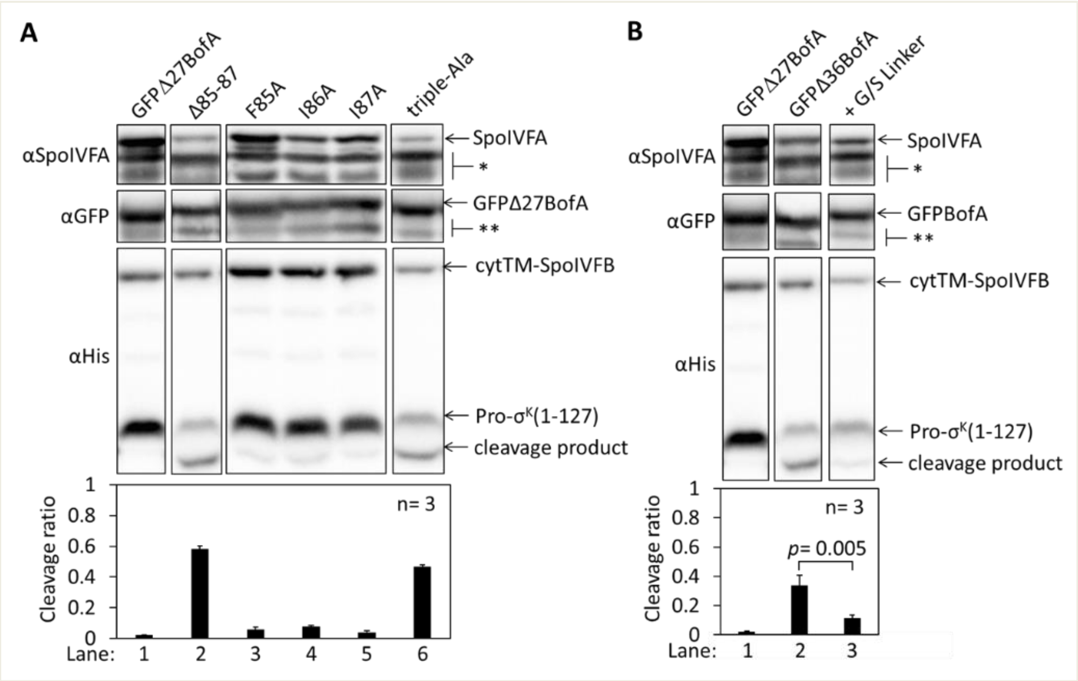


**BofA C-terminal residues and residues preceding predicted TMS2 contribute to inhibition of SpoIVFB in *E. coli*.**

(**A**) Cleavage assays examining the effects of a GFPΔ27BofA truncation and a triple-Ala substitution for the last three residues of GFPΔ27BofA. pET Quartet plasmids were used to produce Pro-$\sigma^K$(1-127), cytTM-SpoIVFB, SpoIVFA, and GFPΔ27BofA from pSO40 as a control (lane 1), GFPΔ27BofA lacking the last three residues (Δ85-87) from pSO43 (lane 2), or GFPΔ27BofA with a triple-Ala substitution for the last three residues from pSO67. Samples collected after 2 h of IPTG induction were subjected to immunoblot analysis, and the graph shows quantification of the cleavage ratio, as explained in the Figure 1B legend. For comparison, lanes 3-5 show data from Figure 2 for single-Ala substitutions in GFPΔ27BofA. The triple-Ala variant increased the cleavage ratio (lane 6), as did the variant lacking the three residues (lane 2). Both variants accumulated normally, but less SpoIVFA accumulated compared to the controls, indicating that residues near the C-terminal end of GFPΔ27BofA affect the synthesis and/or stability of SpoIVFA, and contribute to inhibition of cytTM-SpoIVFB in *E. coli*. (**B**) GFPΔ27BofA lacks TMS1 and all but nine residues preceding predicted TMS2 (36). GFPΔ36BofA additionally lacks the nine residues. Cleavage assays were used to compare inhibition by GFPΔ27BofA, GFPΔ36BofA, or GFPΔ36BofA with a nine-residue glycine/serine (G/S) linker added between GFP and Δ36BofA. pET Quartet plasmids were used to produce Pro-$\sigma^K$(1-127), cytTM-SpoIVFB, SpoIVFA, and GFPΔ27BofA from pSO40 as a control [lane 1, same data as in (**A**)], GFPΔ36BofA from pSO42 (lane 2), or GFPΔ36BofA with the nine-residue G/S linker from pSO69 (lane 3). Samples were subjected to immunoblot analysis and quantification as in (**A**). Samples containing GFPΔ36BofA have a much greater cleavage ratio than samples containing GFPΔ27BofA. Since all four proteins accumulated well in both cases, the nine residues appeared to contribute to the inhibitory function of GFPΔ27BofA. Replacement of the nine residues with the G/S linker decreased the cleavage ratio, based on a Student's two-tailed *t* test (*p* value is indicated), suggesting that moving the GFP tag away from the membrane restored inhibitory function almost completely.

**Figure 2-figure supplement 2-source data 1**
**Immunoblot images (raw and annotated) and quantification of cleavage assays.**

**Figure 3-figure supplement 1**

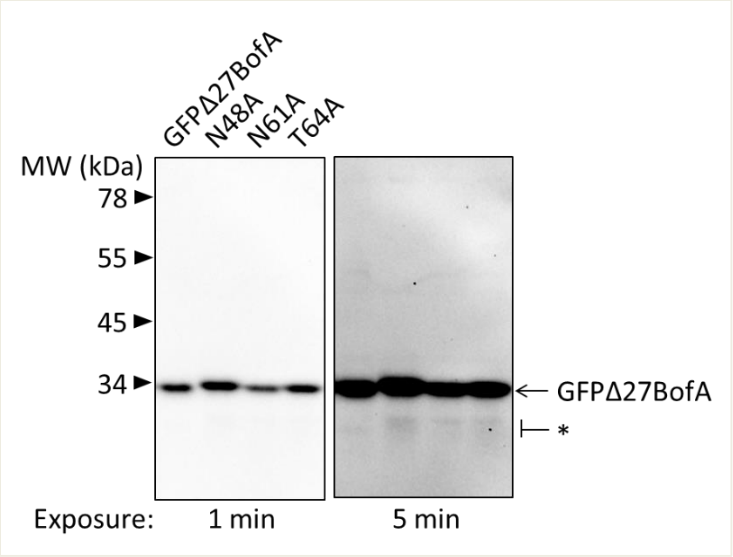


**GFPΔ27BofA variants are intact during *B. subtilis* sporulation.**
A *spoIVB165 bofA::erm* double mutant with $P_{bofA}$-*gfpΔ27bofA* or the indicated mutant version integrated ectopically at *amyE,* were starved to induce sporulation. Samples collected at 3 h poststarvation were subjected to immunoblot analysis with antibodies against GFP. The star (*) indicates very small amounts of potential breakdown species of GFPΔ27BofA and the variants detectable in the long exposure.

**Figure 3-figure supplement 1-source data 1**
**Immunoblot images (raw and annotated).**

**Figure 4-figure supplement 1**

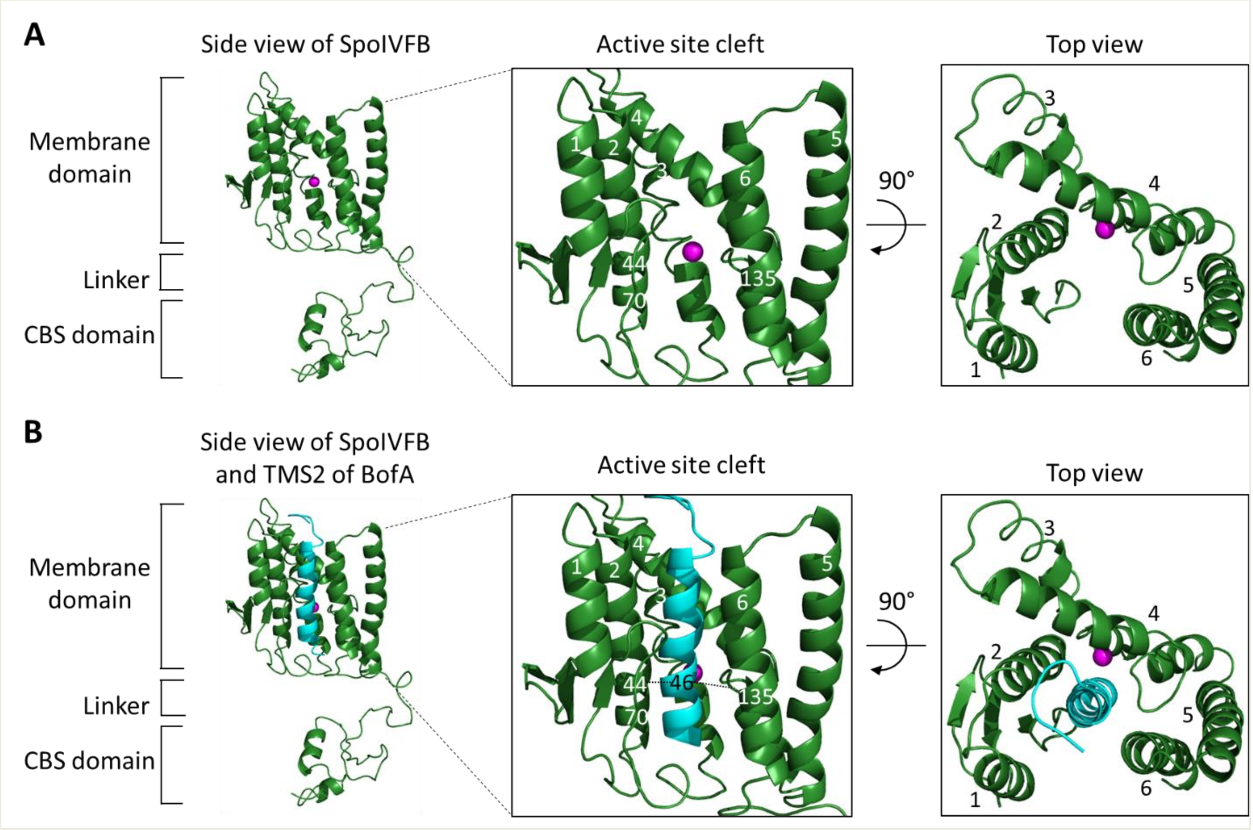


**Models of SpoIVFB and BofA TMS2.**
(**A**) Model of SpoIVFB. At *Left*, a side view of a SpoIVFB monomer. The model shows the six TMSs of the SpoIVFB membrane domain, the zinc ion (magenta) involved in catalysis, the interdomain linker, and the CBS domain. In the enlarged view of the active site cleft (*Center*), TMSs 1– 6 and residues 44, 70, and 135 of SpoIVFB are labeled. At *Right*, a top view is shown. (**B**) Model of SpoIVFB with BofA TMS2. Labeling is as in (**A**) and BofA TMS2 (cyan) is modeled in the SpoIVFB active site cleft. The enlarged view of the active site cleft depicts experimentally observed disulfide cross-links (dashed lines) between BofA C46 and both E44C (in TMS2) and P135C (in a short loop in TMS4) of single-Cys cytTM-SpoIVFB variants (Figure 4).

**Figure 4-figure supplement 1-source data 1**
**PyMOL session file used to produce the images.**

**Figure 4-figure supplement 2**

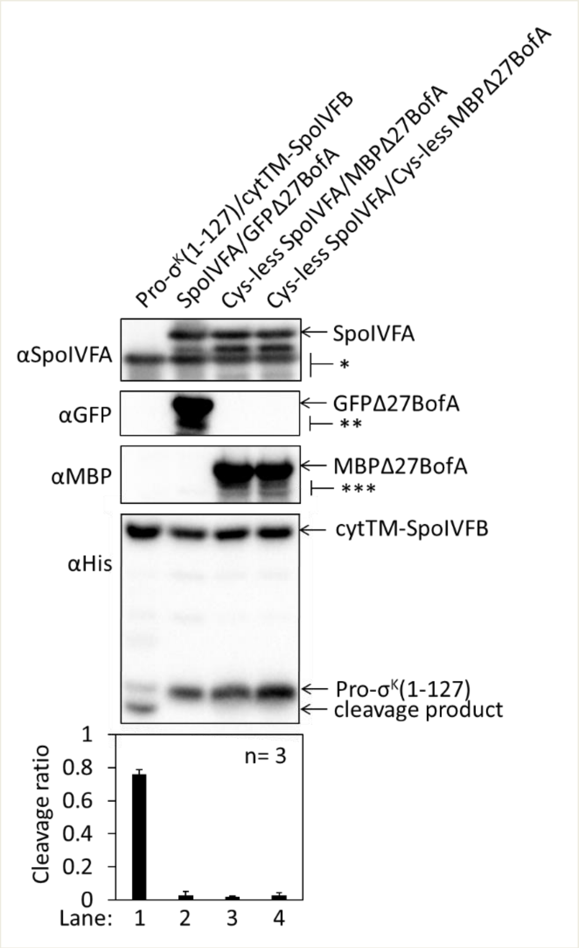


**Cys-less variants of SpoIVFA and MBPΔ27BofA inhibit cleavage of Pro-σ$^K$(1-127) by cytTM-SpoIVFB in *E. coli*.**
Pro-σ$^K$(1-127) and cytTM-SpoIVFB were produced from pYZ2 as a control (lane 1), or pET Quartet plasmids were used to produce Pro-σ$^K$(1-127), cytTM-SpoIVFB, and either SpoIVFA and GFPΔ27BofA from pSO40 as another control (lane 2), Cys-less SpoIVFA and MBPΔ27BofA from pSO90 (lane 3), or Cys-less SpoIVFA and Cys-less MBPΔ27BofA from pSO97 (lane 4). Samples collected after 2 h of IPTG induction were subjected to immunoblot analysis with SpoIVFA, GFP, or penta-His antibodies as indicated. The single star (*) indicates cross-reacting proteins below SpoIVFA. The double (**) and triple (***) star indicate breakdown species of GFPΔ27BofA and MBPΔ27BofA, respectively. A breakdown species below SpoIVFA (not indicated) is observed in some samples. The graph shows quantification of the cleavage ratio, as explained in the Figure 1B legend.

**Figure 4-figure supplement 2-source data 1**
**Immunoblot images (raw and annotated) and quantification of cleavage assays.**

**Figure 4-figure supplement 3**

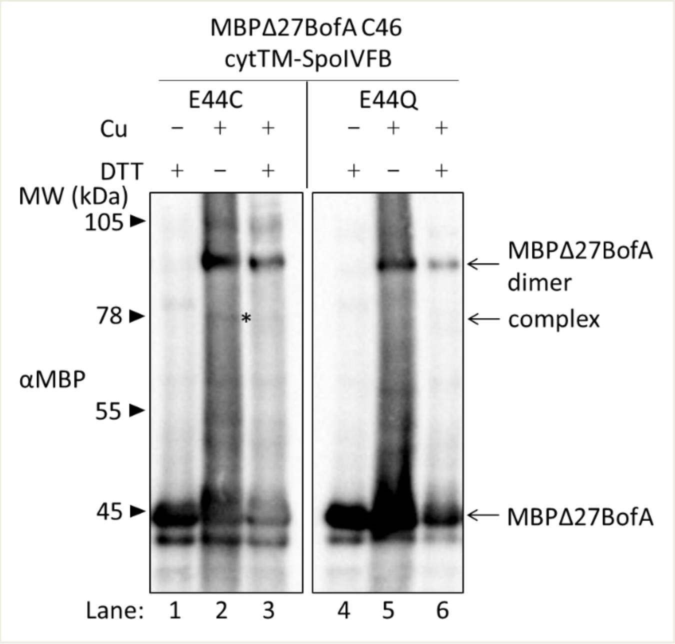


**BofA TMS2 is in proximity to the active site of SpoIVFB.**
Disulfide cross-linking of single-Cys E44C cytTM-SpoIVFB to MBPΔ27BofA C46. pET Quartet plasmids were used to produce single-Cys E44C cytTM-SpoIVFB (pSO91) or Cys-less cytTM-SpoIVFB E44Q as a negative control (pSO94) in combination with MBPΔ27BofA, and Cys-less variants of SpoIVFA and Pro-$\sigma^K$(1-127) in *E. coli*. Samples collected after 2 h of IPTG induction were treated as explained in the Figure 4 legend and subjected to immunoblot analysis with MBP antibodies to visualize MBPΔ27BofA monomer, dimer, and complex with cytTM-SpoIVFB. A representative result from at least two biological replicates is shown. The star indicates the complex in lane 2. The immunoblot images (raw and annotated) are in the Figure 4-source data 1 folder, with explanation on the annotated Figure 4A image.

**Figure 4-figure supplement 4**

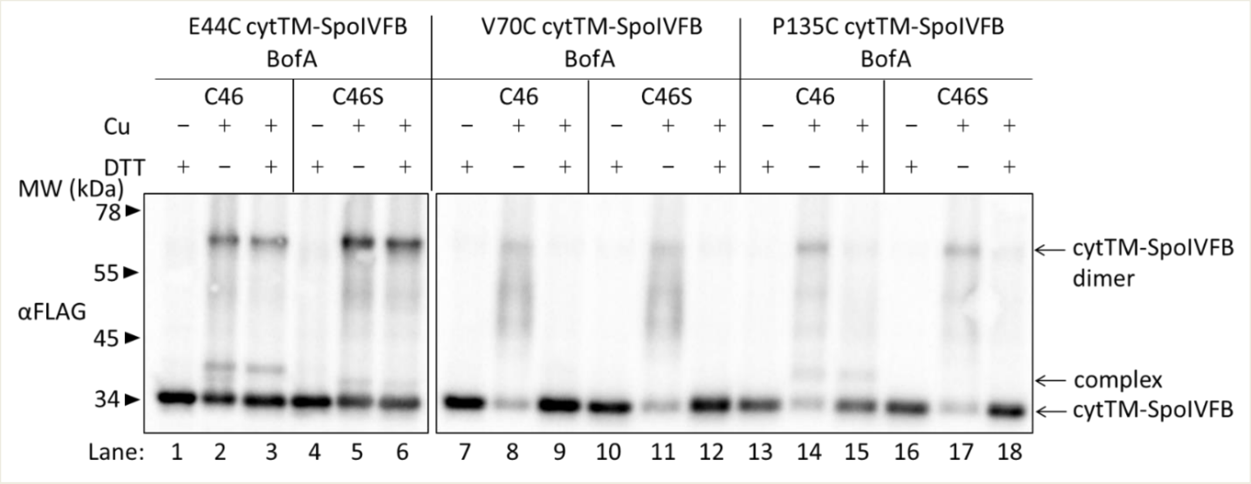


**Full-length BofA is in proximity to the active site of SpoIVFB.**
Disulfide cross-linking of single-Cys cytTM-SpoIVFB variants to BofA C46. pET Quartet plasmids (pSO226-pSO231) were used to produce single-Cys E44C cytTM-SpoIVFB, or single-Cys V70C or P135C cytTM-SpoIVFB E44Q variants, in combination with BofA or BofA C46S, and Cys-less variants of SpoIVFA and Pro-$\sigma^K$(1-127) in *E. coli*. Samples collected after 2 h of IPTG induction were treated as explained in the Figure 4 legend and subjected to immunoblot analysis with FLAG antibodies to visualize cytTM-SpoIVFB monomer, dimer, and complex with full-length BofA. A representative result from at least two biological replicates is shown.

**Figure 4-figure supplement 4-source data 1**
**Immunoblot images (raw and annotated).**

**Figure 4-figure supplement 5**

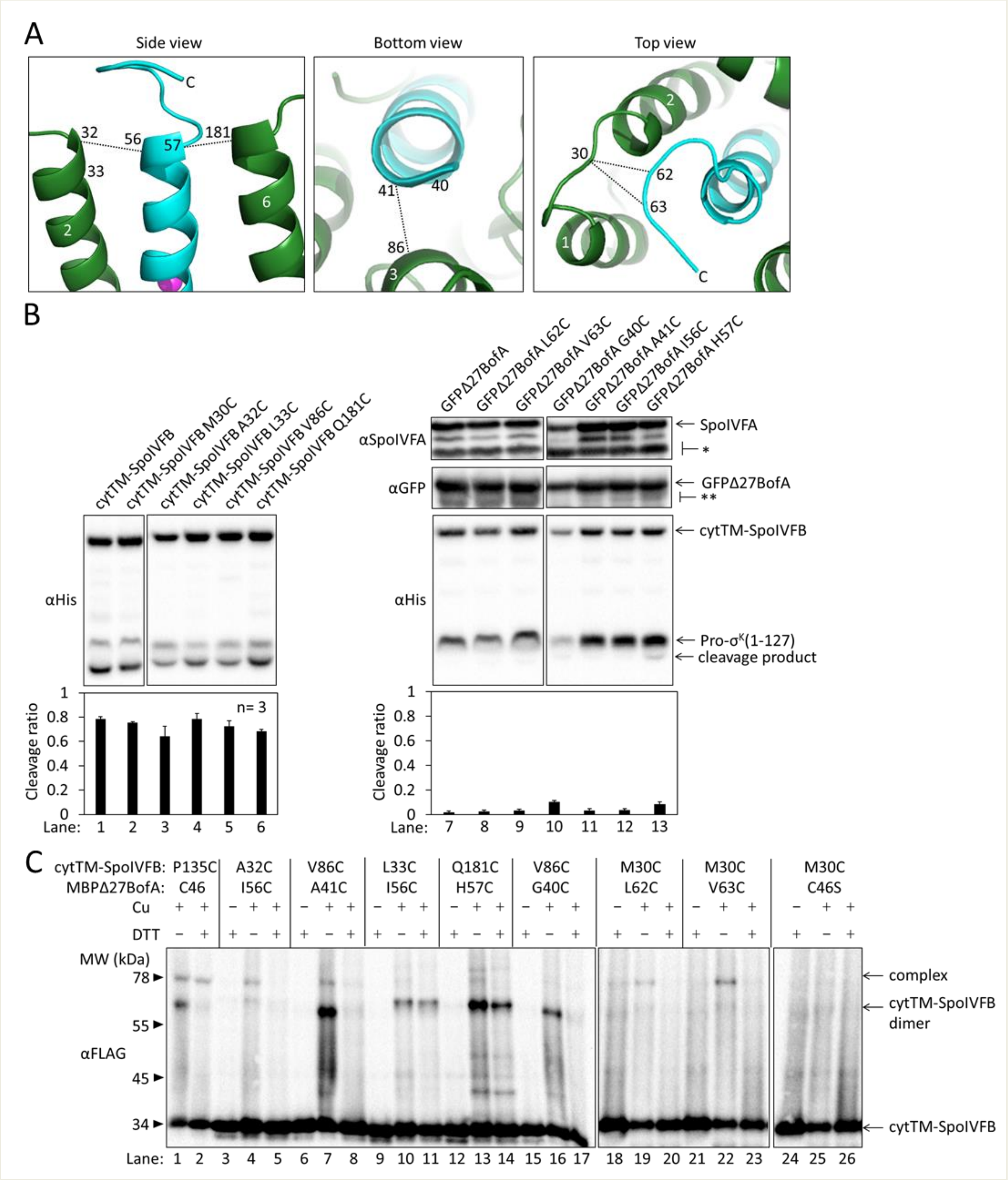


**BofA TMS2 has a preferred orientation in the active site cleft of SpoIVFB.**
(**A**) Enlarged view of the model of SpoIVFB and BofA TMS2 shown in Figure 4-figure supplement 1B, which is based in part on disulfide cross-linked complexes shown in (**C**). At *Left*, the side view of

SpoIVFB TMS2 and TMS6 (green) is shown with the zinc ion (magenta) involved in catalysis and BofA TMS2 (cyan). This view depicts experimentally observed cross-links (dashed lines) between BofA I56C and SpoIVFB A32C, and between BofA H57C and SpoIVFB Q181C. In the bottom view (*Center*), BofA TMS2 is shown with SpoIVFB TMS3. The dashed line indicates a cross-link between BofA A41C and SpoIVFB V86C that was observed. At *Right*, the top view of the model is shown. The dashed lines indicate observed cross-links between SpoIVFB residue M30C (located in the loop connecting TMS1 and TMS2) and BofA L62C and V63C (in a loop near the C-terminal end of TMS2). (**B**) Cleavage assays examining the effects of Cys substitutions for residues of interest in cytTM-SpoIVFB or GFPΔ27BofA. BofA TMS2 was modeled in the SpoIVFB active site cleft based on our initial cross-linking results (Figure 4). The initial model predicted proximity between residues at or near the ends of BofA TMS2 and residues of SpoIVFB, thus identifying residues of interest for cross-linking experiments. First, we examined the effects of Cys substitutions for the residues of interest using cleavage assays. pET Duet plasmids were used to produce Pro-$\sigma^K$(1-127) in combination with cytTM-SpoIVFB from pYZ2 as a control (lane 1) or with the indicated Cys-substituted cytTM-SpoIVFB from pSO141 or pSO256-pSO259 in *E. coli* (*Left*). pET Quartet plasmids were used to produce Pro-$\sigma^K$(1-127), cytTM-SpoIVFB, and SpoIVFA in combination with GFPΔ27BofA from pSO40 as a control (lane 7) or with the indicated Cys-substituted GFPΔ27BofA from pSO142, pSO143, or pSO260-pSO263 in *E. coli* (*Right*). Samples collected after 2 h of IPTG induction were subjected to immunoblot analysis, and the graph shows quantification of the cleavage ratio, as explained in the Figure 1B legend. Since cytTM-SpoIVFB with Cys substitutions for residues of interest cleaved Pro-$\sigma^K$(1-127) (lanes 2-6), we included the inactivating E44Q substitution in the single-Cys cytTM-SpoIVFB variants created for cross-linking. GFPΔ27BofA with Cys substitutions for residues of interest inhibited Pro-$\sigma^K$(1-127) cleavage by cytTM-SpoIVFB, although the G40C and H57C substitutions caused partial loss of inhibition, and the G40C substitution resulted in less accumulation of all four proteins (lanes 8-13). Ala substitutions at these positions had similar effects (Figure 2, lanes 2 & 15). (**C**) Disulfide cross-linking of single-Cys cytTM-SpoIVFB variants to single-Cys MBPΔ27BofA variants. pET Quartet plasmids (pSO93 as a positive control in lanes 1 & 2, pSO147, pSO148, and pSO186-pSO190) were used to produce single-Cys cytTM-SpoIVFB E44Q variants in combination with single-Cys MBPΔ27BofA variants or Cys-less MBPΔ27BofA from pSO144 as a negative control, and Cys-less variants of SpoIVFA and Pro-$\sigma^K$(1-127) in *E. coli*. Samples collected after 2 h of IPTG induction were treated and subjected to immunoblot analysis as explained in the Figure 4A legend. A representative result from two biological replicates is shown. In agreement with the model shown in (**A**), I56C and H57C MBPΔ27BofA variants formed a cross-linked complex with A32C and Q181C cytTM-SpoIVFB variants, respectively, (lanes 4 & 13). The I56C MBPΔ27BofA variant formed very little complex with the L33C cytTM-SpoIVFB variant (lane 10), suggesting that a preferred orientation of BofA TMS2 places I56C farther from L33C than from A32C in TMS2 of SpoIVFB. Similarly, comparison of complex formation by G40C and A41C MBPΔ27BofA variants with the V86C cytTM-SpoIVFB variant (lanes 7 & 16) suggested that a preferred orientation of BofA TMS2 places G40C farther than A41C from V86C near the C-terminal end of SpoIVFB TMS3. Likewise, L62C and V63C MBPΔ27BofA variants formed a cross-linked complex with the M30C cytTM-SpoIVFB variant (lanes 19 & 22), suggesting a loop near the C-terminal end of BofA TMS2 is in proximity to a loop between SpoIVFB TMS1 and TMS2.

**Figure 4-figure supplement 5-source data 1**
**Immunoblot images (raw and annotated) (Figure 4-figure supplement 5 B and C) and quantification of cleavage assays (Figure 4-figure supplement 5B).**

**Figure 5-figure supplement 1**

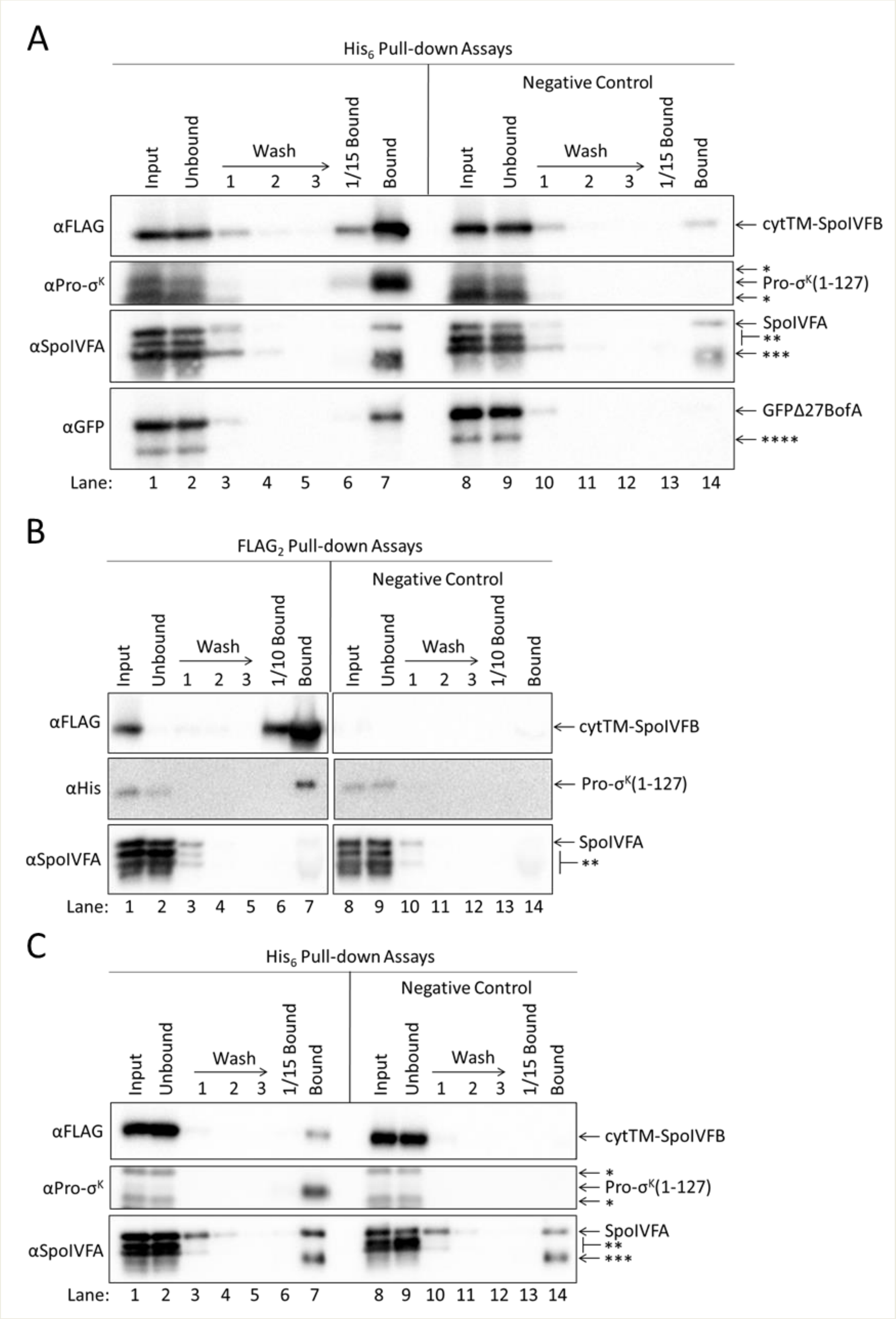

**Neither GFPΔ27BofA nor full-length BofA when co-produced with SpoIVFA prevent Pro-$\sigma^K$(1-127) from interacting with SpoIVFB.**

(**A**) GFPΔ27BofA and SpoIVFA do not prevent SpoIVFB from co-purifying with Pro-$\sigma^K$(1-127). pET Quartet plasmids were used to produce Pro-$\sigma^K$(1-127) (pSO73), or a variant lacking $His_6$ as a negative control (pSO82), in combination with a catalytically-inactive E44C cytTM-SpoIVFB variant containing $FLAG_2$ but lacking $His_6$, GFPΔ27BofA, and SpoIVFA in *E. coli*. Samples collected after 2 h of IPTG induction were subjected to co-purification with cobalt resin. Input, unbound, wash, 1/15 bound (diluted to match input), and (undiluted) bound samples were subjected to immunoblot analysis with FLAG, Pro-$\sigma^K$, SpoIVFA, and GFP antibodies as indicated. The single star (*) indicates cross-reacting proteins above and below Pro-$\sigma^K$(1-127) that fail to co-purify. The double star (**) indicates cross-reacting proteins below SpoIVFA that fail to co-purify. The triple star (***) indicates a putative breakdown species of SpoIVFA that appears to co-purify, but also binds non-specifically. The quadruple star (****) indicates a cross-reacting protein or breakdown species of GFPΔ27BofA that fails to co-purify. All four proteins were seen in the bound sample (lane 7). Only Pro-$\sigma^K$(1-127) and the cytTM-SpoIVFB variant were detected in the diluted bound sample (lane 6). Most of the cytTM-SpoIVFB variant, GFPΔ27BofA, and SpoIVFA were observed in the unbound sample (lane 2), indicating inefficient co-purification. A negative control with a Pro-$\sigma^K$(1-127) variant lacking the $His_6$ tag showed none of the Pro-$\sigma^K$(1-127) variant or GFPΔ27BofA in the bound sample, but a small amount of the cytTM-SpoIVFB variant and considerable SpoIVFA were detected (lane 14), indicative of nonspecific binding to the resin. In the case of SpoIVFA, nonspecific binding rather than co-purification with Pro-$\sigma^K$(1-127) appears to account for most of the signal in lane 7. A putative SpoIVFA breakdown species (indicated by ***) exhibited a similar pattern of abundance in samples as intact SpoIVFA. (**B**) Full-length BofA and SpoIVFA do not prevent Pro-$\sigma^K$(1-127) from co-purifying with SpoIVFB. pET Quartet plasmids were used to produce a catalytically-inactive E44C cytTM-SpoIVFB variant containing $FLAG_2$ but lacking $His_6$ (pSO215), or a variant lacking $FLAG_2$ as a negative control (pSO217), in combination with Pro-$\sigma^K$(1-127), BofA, and SpoIVFA in *E. coli*. Samples collected after 2 h of IPTG induction were subjected to co-immunoprecipitation with anti-FLAG antibody beads. Input, unbound, wash, 1/10 bound (diluted to match input), and (undiluted) bound samples were subjected to immunoblot analysis with FLAG, penta-His, and SpoIVFA antibodies as indicated. Stars indicate proteins as in (**A**). (**C**) Full-length BofA and SpoIVFA do not prevent SpoIVFB from co-purifying with Pro-$\sigma^K$(1-127). pET Quartet plasmids were used to produce Pro-$\sigma^K$(1-127) (pSO215), or a variant lacking $His_6$ as a negative control (pSO216), in combination with a catalytically-inactive E44C cytTM-SpoIVFB variant containing $FLAG_2$ but lacking $His_6$, BofA, and SpoIVFA in *E. coli*. Samples collected after 2 h of IPTG induction were subjected to co-purification with cobalt resin. Input, unbound, wash, 1/15 bound (diluted to match input), and (undiluted) bound samples were subjected to immunoblot analysis with FLAG, Pro-$\sigma^K$, and SpoIVFA antibodies as indicated. Stars indicate proteins as in (**A**). A representative result from two biological replicates is shown in each panel. BofA and SpoIVFA did not completely prevent Pro-$\sigma^K$(1-127) from interacting with the cytTM-SpoIVFB variant in (**B**) or (**C**) (lane 7 in each panel). We note that co-production of BofA decreased the accumulation of Pro-$\sigma^K$(1-127) in the input samples (lane 1 in each panel) compared to co-production of GFPΔ27BofA (lane 1 in Figure 5A and in **A**). We also note that SpoIVFA failed to co-purify with the cytTM-SpoIVFB variant when BofA was co-produced (**B**, lane 7), in contrast to the result when GFPΔ27BofA was co-produced (Figure 5, lane 4). Perhaps BofA decreased Pro-$\sigma^K$(1-127) accumulation and SpoIVFA co-purification more than GFPΔ27BofA because TMS1 in full-length BofA hinders the interaction between Pro-$\sigma^K$(1-127) and the cytTM-SpoIVFB variant, making Pro-$\sigma^K$(1-127) more susceptible to degradation, which may impair SpoIVFA co-purification.

**Figure 5-figure supplement 1-source data 1**
**Immunoblot images (raw and annotated).**

**Figure 5-figure supplement 2**

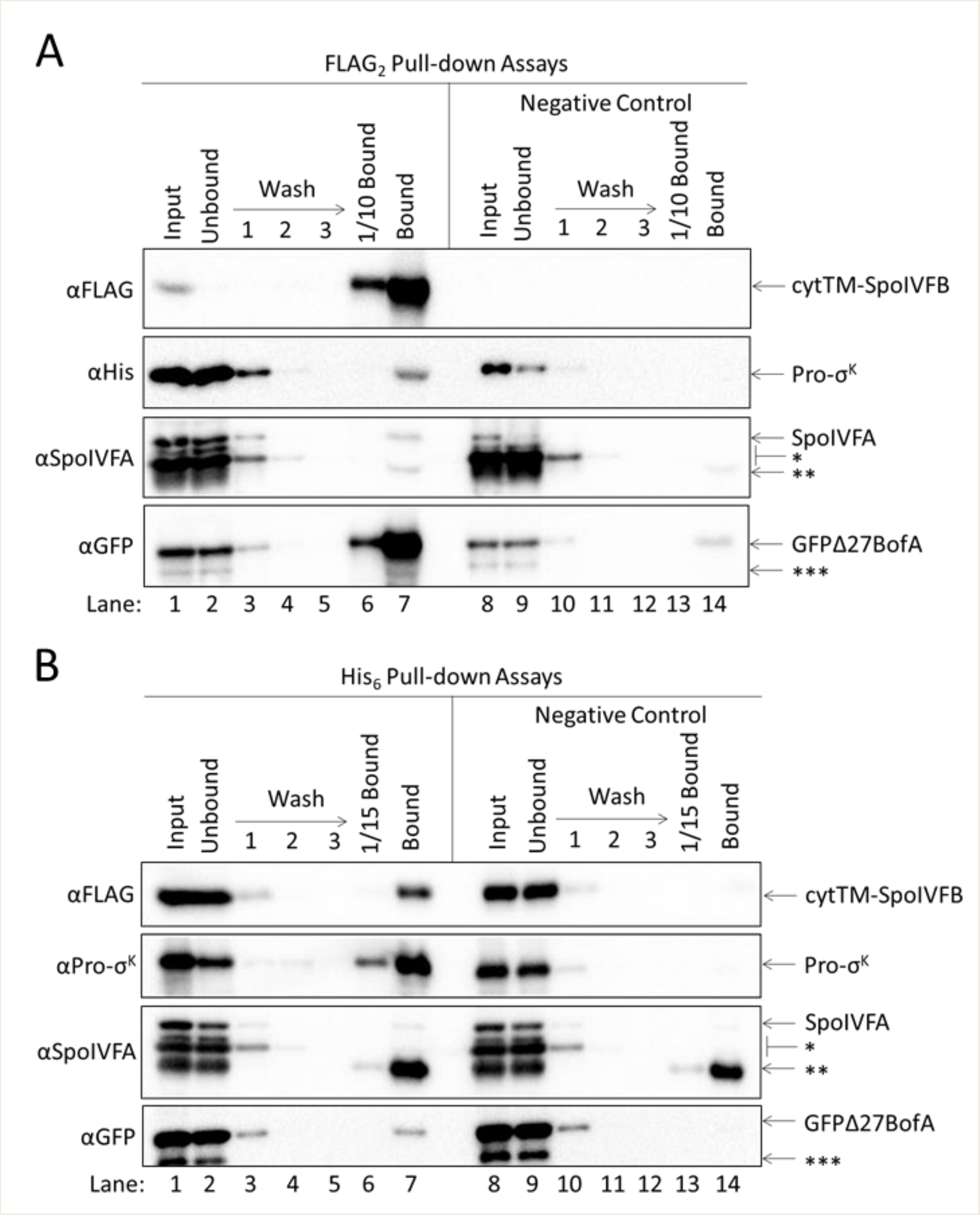


**GFPΔ27BofA and SpoIVFA do not prevent full-length Pro-σ$^K$ from interacting with SpoIVFB.** (**A**) GFPΔ27BofA and SpoIVFA do not prevent Pro-σ$^K$ from co-purifying with SpoIVFB. pET Quartet plasmids were used to produce a catalytically-inactive E44C cytTM-SpoIVFB variant containing FLAG$_2$ but lacking His$_6$ (pSO211), or a variant lacking FLAG$_2$ as a negative control (pSO221), in combination with Pro-σ$^K$-His$_6$, GFPΔ27BofA, and SpoIVFA in *E. coli*. Samples collected after 2 h of IPTG induction were subjected to co-immunoprecipitation with anti-FLAG antibody beads. Input, unbound, wash, 1/10 bound (diluted to match input), and (undiluted) bound samples were subjected to immunoblot analysis with FLAG, penta-His, SpoIVFA, and GFP antibodies as indicated. The single star (*) indicates cross-reacting proteins below SpoIVFA that fail to co-purify. The double star (**) indicates a putative breakdown species of SpoIVFA that appears to co-purify, but also binds non-specifically. The triple star (***) indicates a cross-reacting protein or breakdown species of GFPΔ27BofA that fails to co-purify.

(**B**) GFPΔ27BofA and SpoIVFA do not prevent SpoIVFB from co-purifying with Pro-$\sigma^K$. pET Quartet plasmids were used to produce Pro-$\sigma^K$-$His_6$ (pSO211), or a variant lacking $His_6$ as a negative control (pSO220), in combination with a catalytically-inactive E44C cytTM-SpoIVFB variant containing $FLAG_2$ but lacking $His_6$, GFPΔ27BofA, and SpoIVFA in *E. coli*. Samples collected after 2 h of IPTG induction were subjected to co-purification with cobalt resin. Input, unbound, wash, 1/15 bound (diluted to match input), and (undiluted) bound samples were subjected to immunoblot analysis with FLAG, Pro-$\sigma^K$, SpoIVFA, and GFP antibodies as indicated. Stars indicate proteins as in (**A**). A representative result from two biological replicates is shown in each panel. We note that when Pro-$\sigma^K$-$His_6$ was co-produced rather than Pro-$\sigma^K$(1-127), less SpoIVFA and more GFPΔ27BofA co-purified with the cytTM-SpoIVFB variant (compare **A** and Figure 5A), and less of both inhibitory proteins co-purified with Pro-$\sigma^K$-$His_6$ (compare **B** and Figure 5-figure supplement 1A), consistent with the notion that the C-terminal half of full-length Pro-$\sigma^K$ affects complex formation.

**Figure 5-figure supplement 2-source data 1**
**Immunoblot images (raw and annotated).**

**Figure 5-figure supplement 3**

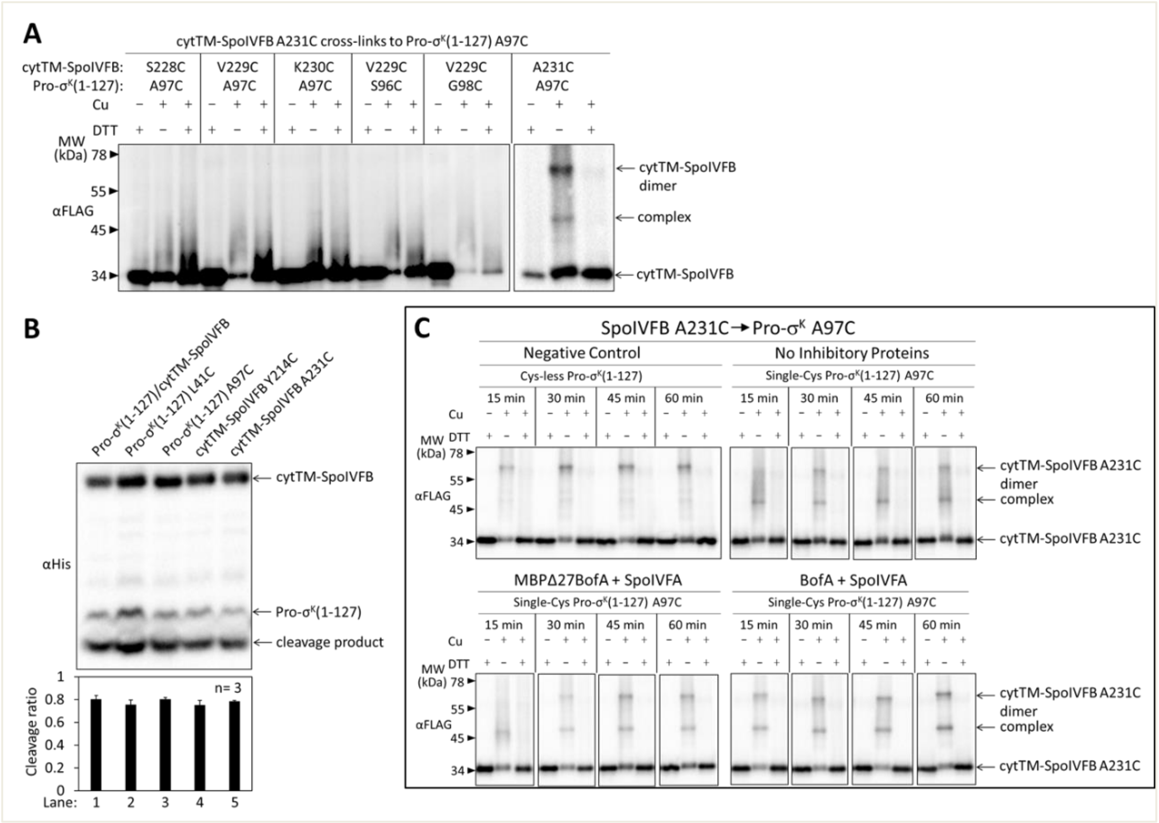


**Disulfide cross-linking between the cytTM-SpoIVFB CBS domain and the Pro-σ$^K$(1-127) C-terminal region.**

(**A**) Cross-linking between single-Cys cytTM-SpoIVFB variants and single-Cys Pro-σ$^K$(1-127) variants. pET Duet plasmids (pSO122-pSO126, pSO130) were used to produce single-Cys S228C, V229C, K230C or A231C cytTM-SpoIVFB E44Q variants in combination with single-Cys S96C, A97C, or G98C Pro-σ$^K$(1-127) variants in *E. coli*. Samples collected after 2 h of IPTG induction were treated and subjected to immunoblot analysis as explained in the Figure 4A legend. A representative result from two biological replicates is shown. (**B**) Cleavage assays examining the effects of Cys substitutions in cytTM-SpoIVFB or Pro-σ$^K$(1-127). pET Duet plasmids were used to produce Pro-σ$^K$(1-127) and cytTM-SpoIVFB from pYZ2 as a control (lane 1), cytTM-SpoIVFB and the indicated Cys-substituted Pro-σ$^K$(1-127) from pSO157 or pSO158 (lanes 2 & 3), or Pro-σ$^K$(1-127) and the indicated Cys-substituted cytTM-SpoIVFB from pSO159 or pSO160 (lanes 4 & 5) in *E. coli*. Samples collected after 2 h of IPTG induction were subjected to immunoblot analysis, and the graph shows quantification of the cleavage ratio, as explained in the Figure 1B legend. (**C**) Time course of cross-linking between the single-Cys A231C CBS domain variant of cytTM-SpoIVFB E44Q and single-Cys A97C Pro-σ$^K$(1-127) in the absence or presence of inhibitory proteins. See the Figure 5B legend for explanation of the experiment. A representative result from two biological replicates is shown.

**Figure 5-figure supplement 3-source data 1**
**Immunoblot images (raw and annotated) and quantification of cleavage assays (Figure 5-figure supplement 3B).**

**Figure 6-figure supplement 1**

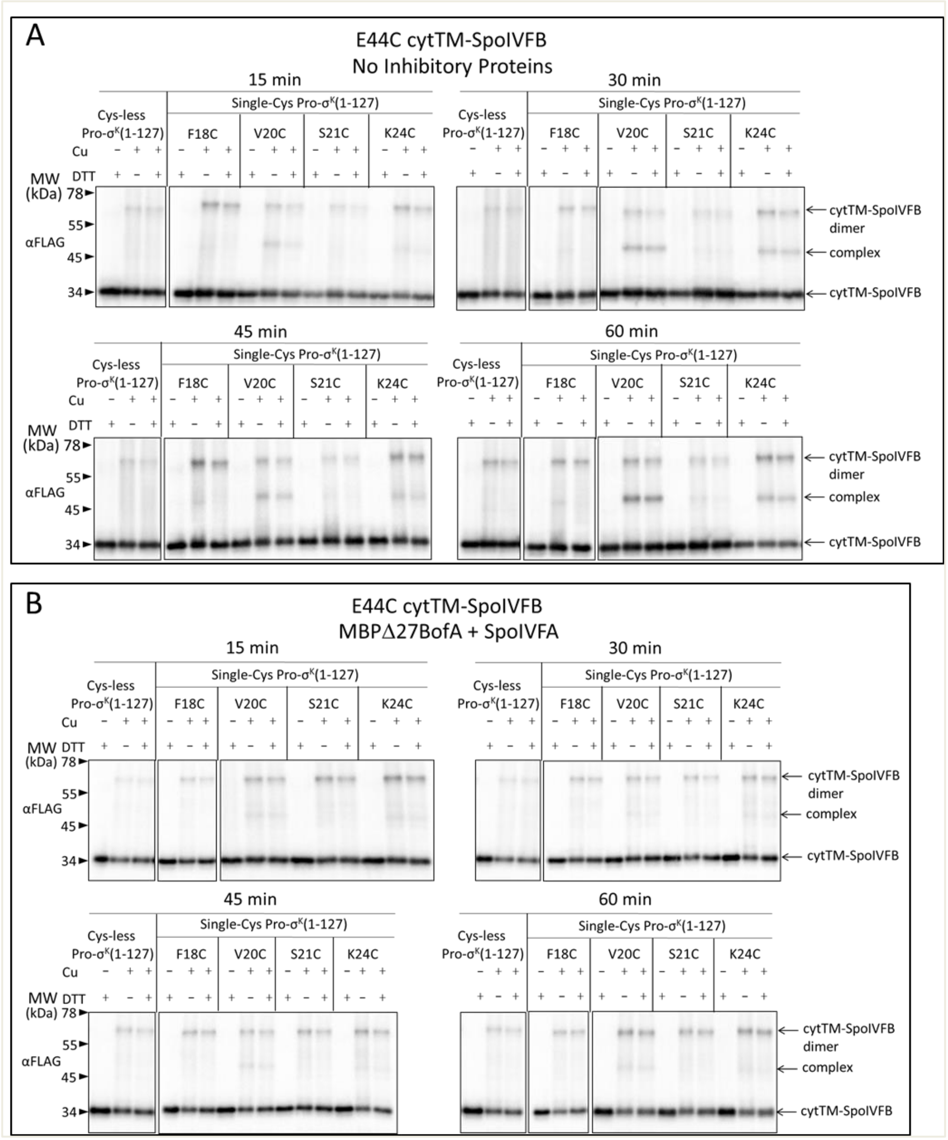


**Disulfide cross-linking between E44C at the active site of cytTM-SpoIVFB and the Proregion of Pro-σ$^K$(1-127) in the absence of inhibitory proteins or in the presence of MBPΔ27BofA and SpoIVFA.**

(**A**) Time course of cross-linking between single-Cys E44C cytTM-SpoIVFB and single-Cys Pro-σ$^K$(1-127) variants in the absence of inhibitory proteins. See the Figure 6A legend for explanation of the experiment. (**B**) Time course of cross-linking between single-Cys E44C cytTM-SpoIVFB and single-Cys Pro-σ$^K$(1-127) variants in the presence of Cys-less variants of MBPΔ27BofA and SpoIVFA. See the Figure 6C legend for explanation of the experiment. Representative results from two biological replicates are shown in (**A**) and (**B**).

**Figure 6-figure supplement 1-source data 1**
**Immunoblot images (raw and annotated) (Figure 6-figure supplement 1A).**

**Figure 6-figure supplement 1-source data 2**
**Immunoblot images (raw and annotated) (Figure 6-figure supplement 1B).**

**Figure 6-figure supplement 2**

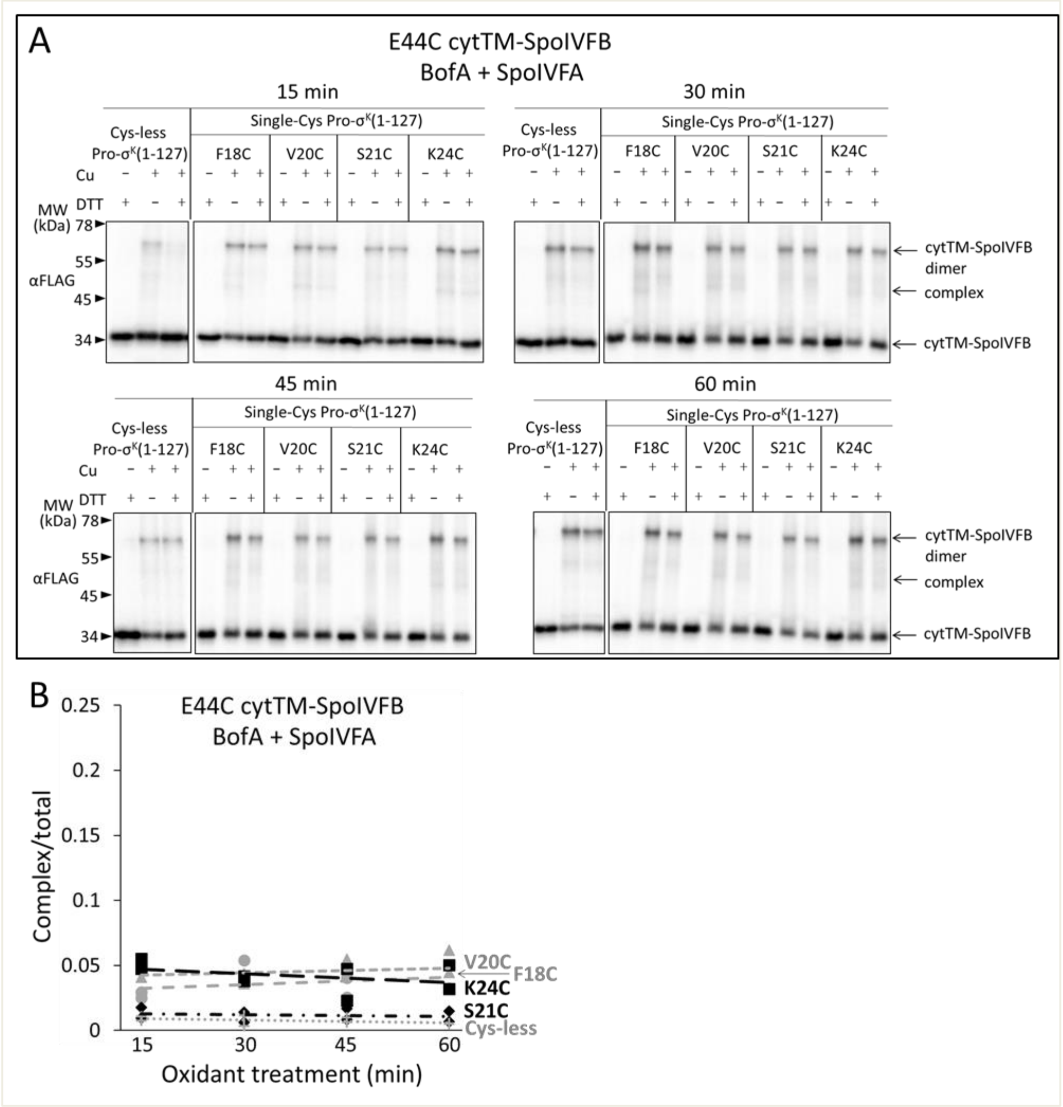


**Full-length BofA and SpoIVFA decrease cross-linking between E44C cytTM-SpoIVFB and V20C or K24C Pro-σ$^K$(1-127).**

(**A**) Time course of cross-linking between single-Cys E44C cytTM-SpoIVFB and single-Cys Pro-σ$^K$(1-127) variants in the presence of Cys-less variants of BofA and SpoIVFA. pET Quartet plasmids were used to produce single-Cys E44C cytTM-SpoIVFB in combination with single-Cys F18C (pSO238), V20C (pSO234), S21C (pSO235), or K24C (pSO239) Pro-σ$^K$(1-127), or with Cys-less Pro-σ$^K$(1-127) (pSO229) as a negative control, and Cys-less variants of BofA and SpoIVFA in *E. coli*. Samples collected after 2 h of IPTG induction were treated and subjected to immunoblot analysis as explained in the Figure 6A legend. A representative result from two biological replicates is shown. (**B**) Quantification of cross-linking for the experiment described in (**A**). Abundance of the complex was divided by the total amount of cytTM-SpoIVFB monomer, dimer, and complex. The ratio over time was plotted (n=2) with a best-fit trend line.

**Figure 6-figure supplement 2-source data 1**
**Immunoblot images (raw and annotated) (Figure 6-figure supplement 2A) and quantification of cross-linking (Figure 6-figure supplement 2B).**

**Figure 6-figure supplement 3**

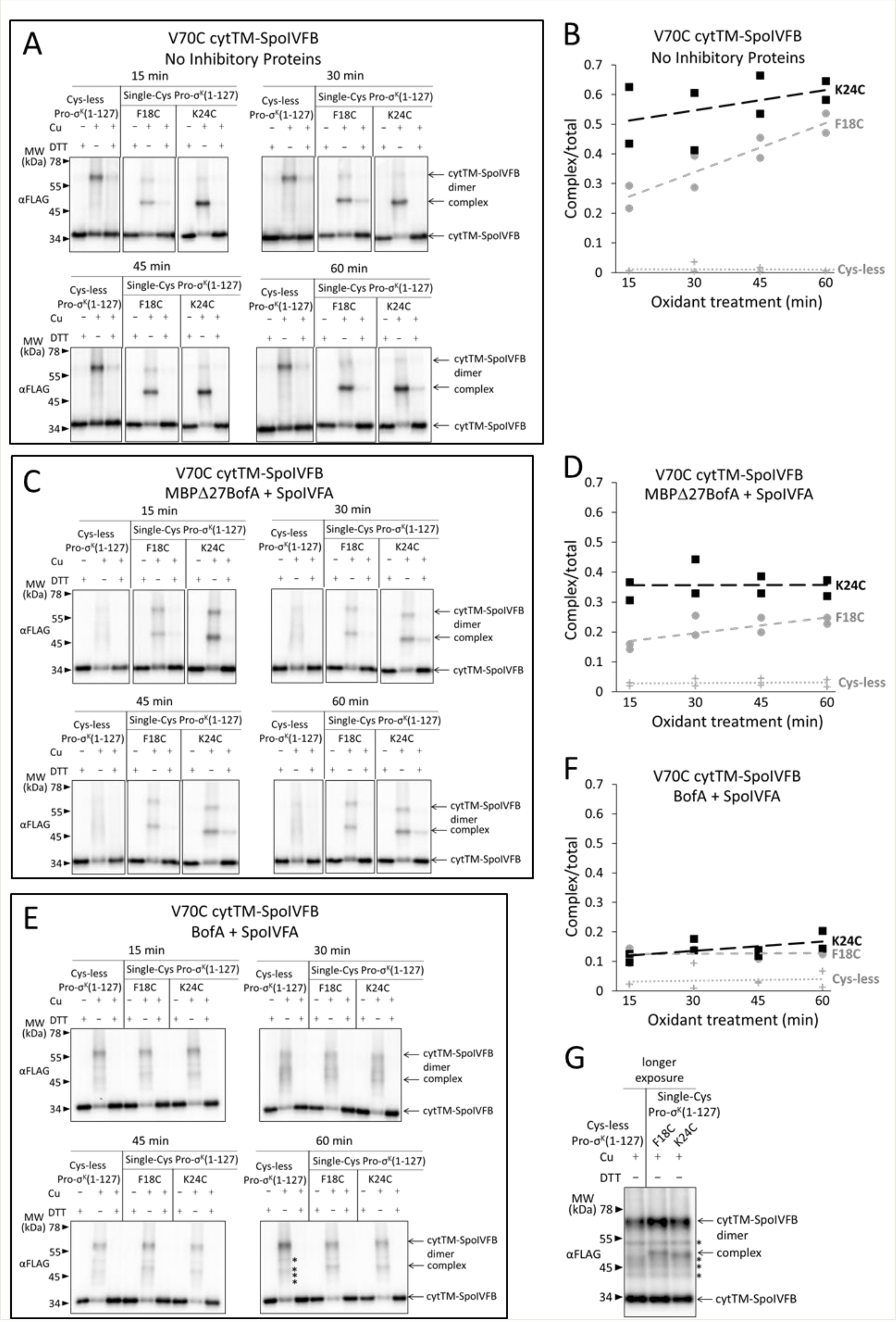

**Disulfide cross-linking between V70C in the cytTM-SpoIVFB membrane-reentrant loop and F18C or K24C in the Pro-$\sigma^K$(1-127) N-terminal region is decreased more by full-length BofA than by MBPΔ27BofA (lacking TMS1).**

(**A**) Time course of cross-linking between the single-Cys V70C cytTM-SpoIVFB E44Q and single-Cys Pro-$\sigma^K$(1-127) variants in the absence of inhibitory proteins. pET Duet plasmids were used to produce single-Cys V70C cytTM-SpoIVFB E44Q in combination with single-Cys F18C (pSO168) or K24C (pSO134) Pro-$\sigma^K$(1-127), or with Cys-less Pro-$\sigma^K$(1-127) (pSO136) as a negative control, in *E. coli*. Samples collected after 2 h of IPTG induction were treated and subjected to immunoblot analysis as explained in the Figure 6A legend. A representative result from two biological replicates is shown. (**B**) Quantification of cross-linking for the experiment described in (**A**). Abundance of the complex was divided by the total amount of cytTM-SpoIVFB monomer, dimer, and complex. The ratio over time was plotted (n=2) with a best-fit trend line. (**C**) Time course of cross-linking between single-Cys V70C cytTM-SpoIVFB E44Q and single-Cys Pro-$\sigma^K$(1-127) variants in the presence of Cys-less variants of MBPΔ27BofA and SpoIVFA. pET Quartet plasmids were used to produce single-Cys V70C cytTM-SpoIVFB E44Q in combination with single-Cys F18C (pSO164) or K24C (pSO132) Pro-$\sigma^K$(1-127), or with Cys-less Pro-$\sigma^K$(1-127) (pSO111) as a negative control, and Cys-less variants of MBPΔ27BofA and SpoIVFA in *E. coli*. Samples collected after 2 h of IPTG induction were treated and subjected to immunoblot analysis as explained in the Figure 6A legend. A representative result from two biological replicates is shown. (**D**) Quantification of cross-linking for the experiment described in (**C**). Quantification was performed as described in (**B**). (**E**) Time course of cross-linking between single-Cys V70C cytTM-SpoIVFB E44Q and single-Cys Pro-$\sigma^K$(1-127) variants in the presence of Cys-less variants of full-length BofA and SpoIVFA. pET Quartet plasmids were used to produce the single-Cys V70C cytTM-SpoIVFB E44Q in combination with single-Cys F18C (pSO236) or K24C (pSO237) Pro-$\sigma^K$(1-127), or with Cys-less Pro-$\sigma^K$(1-127) (pSO230) as a negative control, and Cys-less variants of BofA and SpoIVFA in *E. coli*. Samples collected after 2 h of IPTG induction were treated and subjected to immunoblot analysis as explained in the Figure 6A legend. A representative result from two biological replicates is shown. (**F**) Quantification of cross-linking for the experiment described in *E*. Quantification was performed as described in (**B**). (**G**) Immunoblot of 60-min samples (Cu +) from the experiment described in *E* with a longer exposure (10 sec). Stars (*) indicate four novel species.

**Figure 6-figure supplement 3-source data 1**
**Immunoblot images (raw and annotated) (Figure 6-figure supplement 3 A, C, E, and G) and quantification of cross-linking (Figure 6-figure supplement 3 B, D, and F).**

**Figure 6-figure supplement 4**

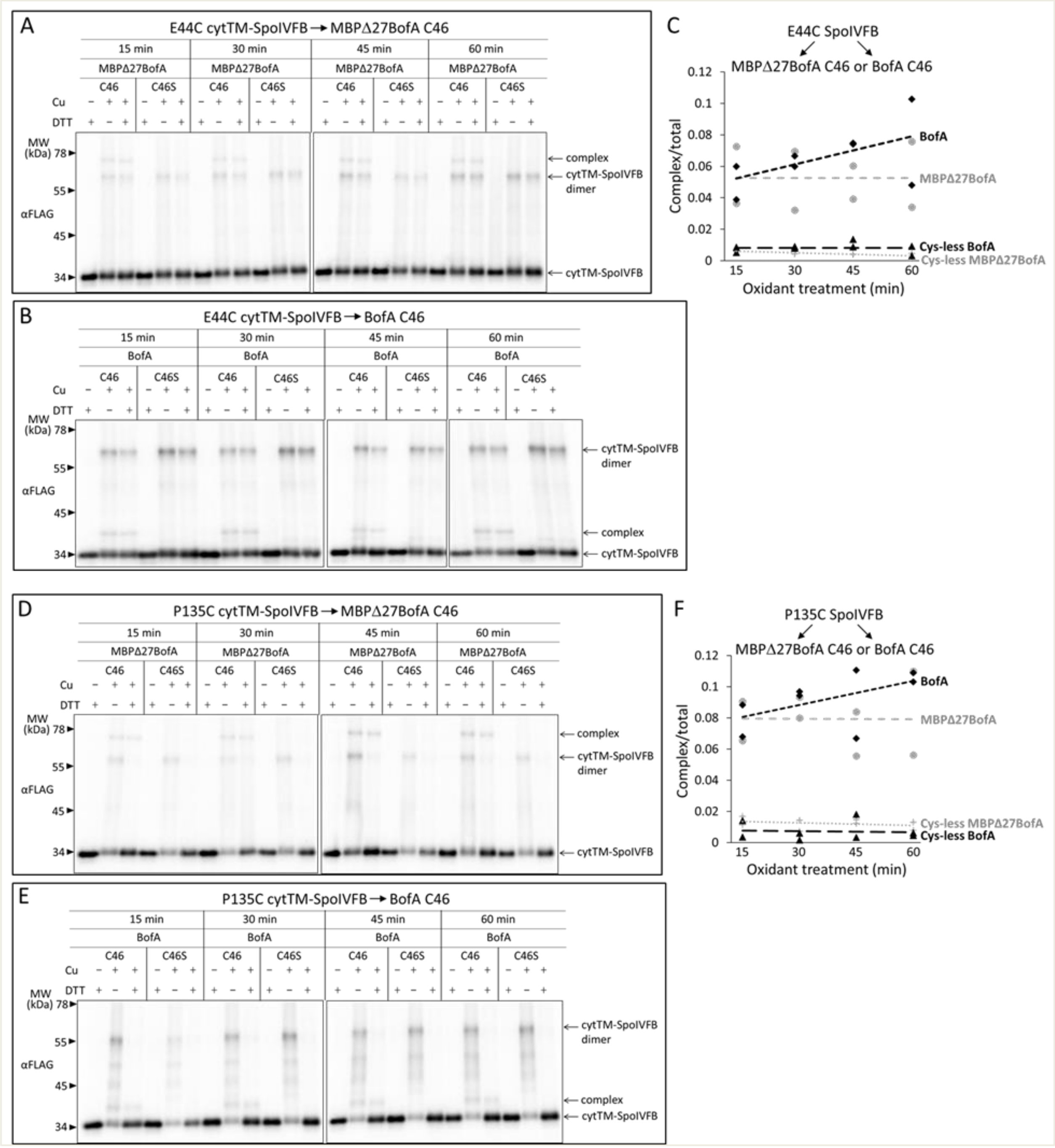


**Comparison of disulfide cross-linking between C46 in TMS2 of full-length BofA or MBPΔ27BofA (lacking TMS1) and E44C at or P135C near the active site of cytTM-SpoIVFB.**

(**A** and **B**) Time courses of cross-linking between single-Cys E44C cytTM-SpoIVFB and MBPΔ27BofA C46 or full-length BofA C46. pET Quartet plasmids were used to produce single-Cys E44C cytTM-SpoIVFB in combination with MBPΔ27BofA C46 (pSO91) or BofA C46 (pSO226), or with Cys-less MBPΔ27BofA C46S (pSO110) or BofA C46S (pSO229) variants as negative controls, and Cys-less variants of Pro-$\sigma^K$(1-127) and SpoIVFA in *E. coli*. Samples collected after 2 h of IPTG induction were treated and subjected to immunoblot analysis as explained in the Figure 6A legend. A representative result from two biological replicates is shown. (**C**) Quantification of cross-linking for the experiments described in (**A** and **B**). Abundance of the complex was divided by the total amount of cytTM-SpoIVFB monomer, dimer, and complex. The ratio over time was plotted (n=2) with a best-fit trend line. (**D** and **E**) Time courses of cross-linking between single-Cys P135C cytTM-SpoIVFB and MBPΔ27BofA C46 or full-length BofA C46. pET Quartet plasmids were used to produce single-Cys P135C cytTM-SpoIVFB E44Q in combination with MBPΔ27BofA C46 (pSO93) or BofA C46 (pSO228), or with Cys-less MBPΔ27BofA C46S (pSO112) or (BofA C46S (pSO231) variants as negative controls, and Cys-less variants of Pro-$\sigma^K$(1-127) and SpoIVFA in *E. coli*. Samples collected after 2 h of IPTG induction were treated and subjected to immunoblot analysis as explained in the Figure 6A legend. A representative result from two biological replicates is shown. (**F**) Quantification of cross-linking for the experiments described in (**D** and **E**). Quantification was performed as described in (**C**).

**Figure 6-figure supplement 4-source data 1**
**Immunoblot images (raw and annotated) (Figure 6-figure supplement 4 A, B, D, and E) and quantification of cross-linking (Figure 6-figure supplement 4 C and F).**

**Figure 6-figure supplement 5**

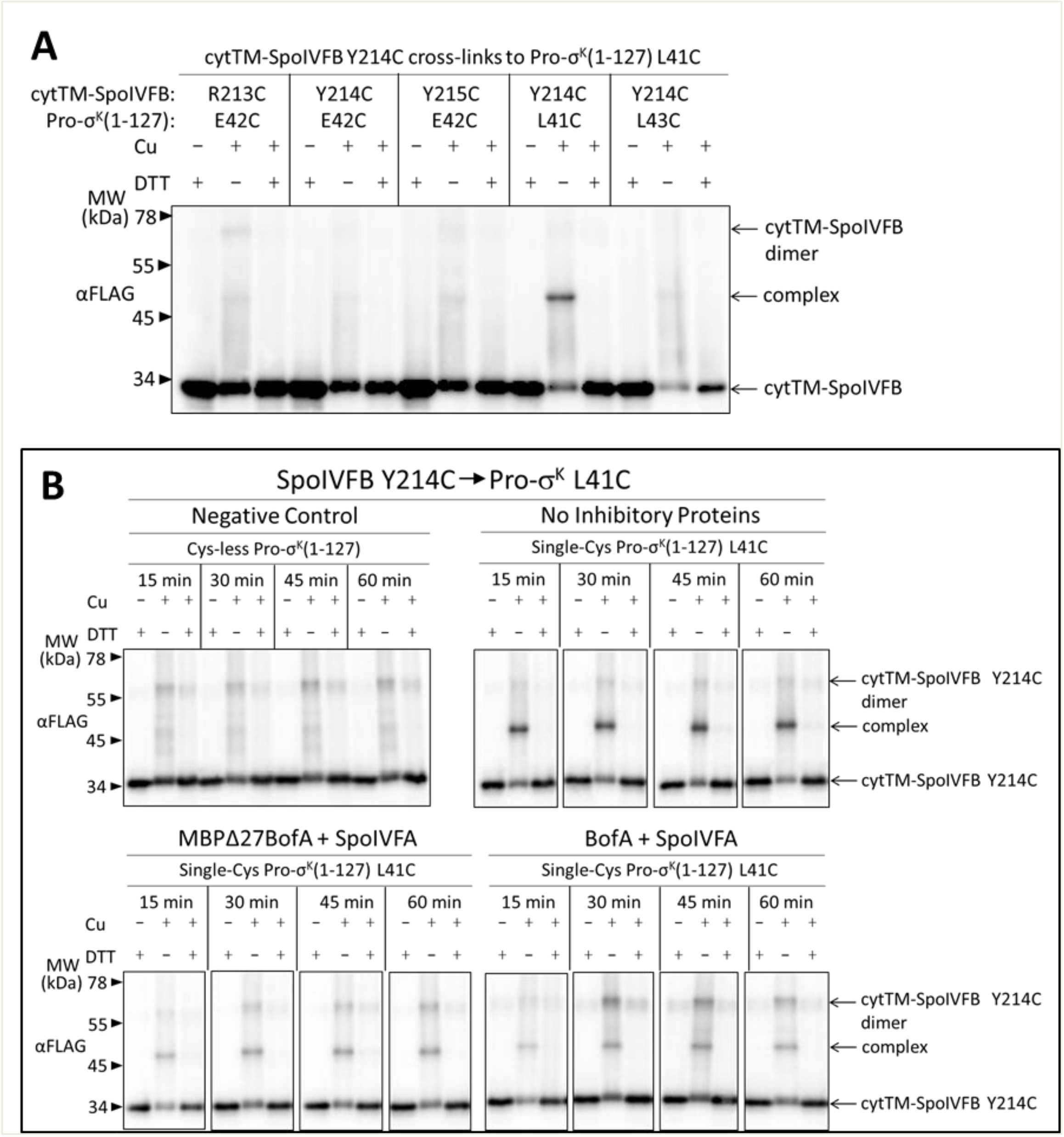


**Disulfide cross-linking between Y214C in the cytTM-SpoIVFB interdomain linker and L41C in the Pro-σ$^{K}$(1-127) N-terminal region is decreased more by full-length BofA than by MBPΔ27BofA (lacking TMS1).**

(**A**) Cross-linking between single-Cys cytTM-SpoIVFB variants and single-Cys Pro-σ$^{K}$(1-127) variants. pET Duet plasmids (pSO117- pSO121) were used to produce single-Cys R213C, Y214C, or Y215C cytTM-SpoIVFB E44Q variants in combination with single-Cys L41C, E42C, or L43C Pro-σ$^{K}$(1-127) variants in *E. coli*. Samples collected after 2 h of IPTG induction were treated and subjected to immunoblot analysis as explained in the Figure 5-figure supplement 3A legend. A representative result from two biological replicates is shown. (**B**) Time course of cross-linking between Y214C in the cytTM-SpoIVFB interdomain linker and L41C in the Pro-σ$^{K}$(1-127) N-terminal region in the absence or presence of inhibitory proteins. pET Duet plasmids were used to produce single-Cys Y214C cytTM-SpoIVFB

E44Q in combination with single-Cys L41C Pro-$\sigma^K$(1-127) from pSO120 or with Cys-less Pro-$\sigma^K$(1-127) from pSO114 as a negative control, in *E. coli*. pET Quartet plasmids were used to produce single-Cys Y214C cytTM-SpoIVFB E44Q, single-Cys L41C Pro-$\sigma^K$(1-127), and Cys-less SpoIVFA in combination with Cys-less MBPΔ27BofA from pSO127 or with Cys-less full-length BofA from pSO245 in *E. coli*. Samples collected after 2 h of IPTG induction were treated and subjected to immunoblot analysis as explained in the Figure 6A legend. A representative result from two biological replicates is shown.

**Figure 6-figure supplement 5-source data 1**
**Immunoblot images (raw and annotated).**

**Figure 7-figure supplement 1**

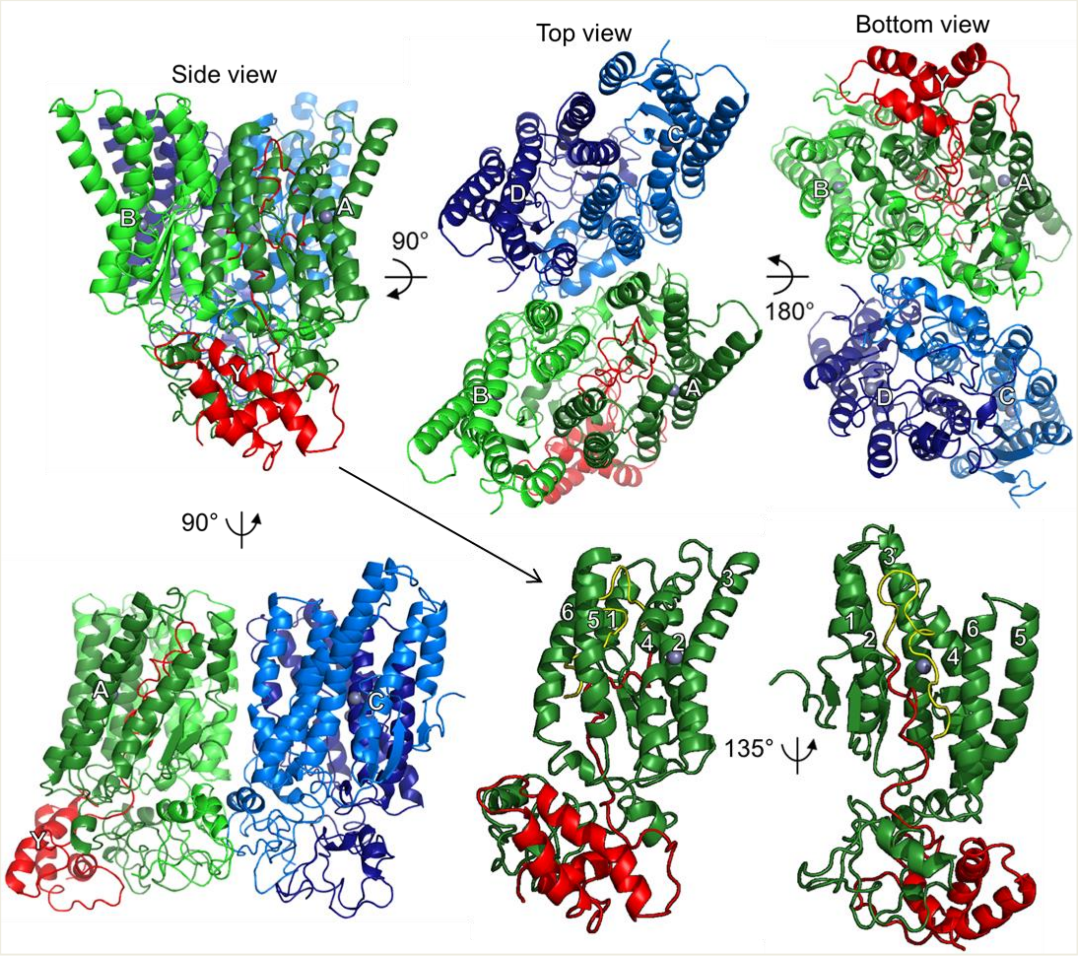


**Model of SpoIVFB tetramer with part of one Pro-σ$^K$ molecule.**
At *Upper Left*, a side view shows the SpoIVFB membrane domains at the top and the CBS domains at the bottom, facing the A (dark green) and B chains (light green), whose CBS domains primarily provide the dimerization interface. The SpoIVFB A chain also interacts with the Pro-σ$^K$ Y chain (residues 1-114) (red). The top view (*Center*) reveals the SpoIVFB C (light blue) and D (dark blue) chains, whose CBS domains dimerize. The bottom view (*Upper Right*) emphasizes the CBS domains, as well as the interface between the A/B and C/D dimers, formed primarily by the CBS domains of the B and C chains. At *Lower Left*, a side view facing the A and C chains also shows the interface between the A/B and C/D dimers of the SpoIVFB tetramer. Each SpoIVFB chain is labeled near its zinc ion (gray), which is hidden in some views. At *Lower Center*, a side view of the SpoIVFB A chain monomer (TMSs 1-6 labeled) interacting with the Pro-σ$^K$ Y chain (Proregion residues 1-21 yellow and σ$^K$ residues 22-114 red), in the same orientation as at *Upper Left* (hence the arrow), but with the other SpoIVFB chains hidden. At *Lower Right*, a side view into the active site cleft of the SpoIVFB A chain emphasizes proximity between the zinc ion and the cleavage site in the Pro-σ$^K$ Y chain (between residues 21 and 22).

**Figure 7-figure supplement 1-source data 1**
**PyMOL session file used to produce the images and PDB file of the model of a SpoIVFB tetramer with part of one molecule of Pro-σ$^K$.**
