## Supplementary File 1 for "Inhibitory proteins block substrate access by occupying the active site cleft of *Bacillus subtilis* intramembrane protease SpoIVFB"

**Plasmids used in this study**

| **Name** | **Description** | **Construction** | **Reference** |
| --- | --- | --- | --- |
| pDR95 | Ap^R^ Sp^R^; P*_spoIID_*-GFPΔ27BofA |  | (1) |
| pETproσ^K^ | Km^R^; T7-Pro-σ^K^-His_6_ |  | (2) |
| pPL29 | Ap^R^; single-Cys P135C cytTM-SpoIVFB E44Q-FLAG_2_-His_6_, which also has C35S C165L C167L C172S C246S substitutions in SpoIVFB |  | (3) |
| pSO6 | Ap^R^; T7-GFPΔ36BofA/SpoIVFA | pYZ46 was subjected to SDM^a^ using primers SO-P13 and SO-P14, deleting residues 28-36 from GFPΔ27BofA. | This study |
| pSO10 | Ap^R^; T7-GFPΔ27+Δ85-87BofA/SpoIVFA | pYZ46 was subjected to SDM using primers SO-P11 and SO-P12, deleting residues 85-87 from GFPΔ27BofA. | This study |
| pSO25 | Ap^R^; T7-GFPΔ27BofA G40A/SpoIVFA | pYZ46 was subjected to SDM using primers SO-P48 and SO-P49, substituting G40A in GFPΔ27BofA. | This study |
| pSO26 | Ap^R^; T7-GFPΔ27BofA L44A/SpoIVFA | pYZ46 was subjected to SDM using primers SO-P50 and SO-P51, substituting L44A in GFPΔ27BofA. | This study |
| pSO27 | Ap^R^; T7-GFPΔ27BofA N48A/SpoIVFA | pYZ46 was subjected to SDM using primers SO-P52 and SO-P53, substituting N48A in GFPΔ27BofA. | This study |
| pSO28 | Ap^R^; T7-GFPΔ27BofA G51A/SpoIVFA | pYZ46 was subjected to SDM using primers SO-P54 and SO-P55, substituting G51A in GFPΔ27BofA. | This study |
| pSO29 | Ap^R^; T7-GFPΔ27BofA I60A/SpoIVFA | pYZ46 was subjected to SDM using primers SO-P56 and SO-P57, substituting I60A in GFPΔ27BofA. | This study |
| pSO30 | Ap^R^; T7-GFPΔ27BofA N61A/SpoIVFA | pYZ46 was subjected to SDM using primers SO-P58 and SO-P59, substituting N61A in GFPΔ27BofA. | This study |
| pSO31 | Ap^R^; T7-GFPΔ27BofA T64A/SpoIVFA | pYZ46 was subjected to SDM using primers SO-P60 and SO-P61, substituting T64A in GFPΔ27BofA. | This study |
| pSO32 | Ap^R^; T7-GFPΔ27BofA G69A/SpoIVFA | pYZ46 was subjected to SDM using primers SO-P62 and SO-P63, substituting G69A in GFPΔ27BofA. | This study |
| pSO33 | Ap^R^; T7-GFPΔ27BofA L71A/SpoIVFA | pYZ46 was subjected to SDM using primers SO-P64 and SO-P65, substituting L71A in GFPΔ27BofA. | This study |
| pSO34 | Ap^R^; T7-GFPΔ27BofA G72A/SpoIVFA | pYZ46 was subjected to SDM using primers SO-P66 and SO-P67, substituting G72A in GFPΔ27BofA. | This study |
| pSO35 | Ap^R^; T7-GFPΔ27BofA P74A/SpoIVFA | pYZ46 was subjected to SDM using primers SO-P68 and SO-P69, substituting P74A in GFPΔ27BofA. | This study |
| pSO36 | Ap^R^; T7-GFPΔ27BofA G75A/SpoIVFA | pYZ46 was subjected to SDM using primers SO-P70 and SO-P71, substituting G75A in GFPΔ27BofA. | This study |
| pSO37 | Ap^R^; T7-GFPΔ27BofA L79A/SpoIVFA | pYZ46 was subjected to SDM using primers SO-P72 and SO-P73, substituting L79A in GFPΔ27BofA. | This study |
| pSO38 | Ap^R^; T7-GFPΔ27BofA I86A/SpoIVFA | pYZ46 was subjected to SDM using primers SO-P74 and SO-P75, substituting I86A in GFPΔ27BofA. | This study |
| pSO39 | Ap^R^; T7-GFPΔ27BofA I87A/SpoIVFA | pYZ46 was subjected to SDM using primers SO-P76 and SO-P77, substituting I87A in GFPΔ27BofA. | This study |
| pSO40 | Km^R^; T7-Pro-σ^K^(1-127)-His_6_/T7-cytTM-SpoIVFB-FLAG_2_-His_6_/T7-GFPΔ27BofA/SpoIVFA | Fragment *T7-gfpΔ27bofA/spoIVFA* was amplified from pYZ46 using primers SO-P82 and SO-P90. Vector pYZ2 was amplified using primers SO-P80 and SO-P89. Fragment was joined to pYZ2 using GA^b^. | This study |
| pSO42 | Km^R^; T7-Pro-σ^K^(1-127)-His_6_/T7-cytTM-SpoIVFB-FLAG_2_-His_6_/T7-GFPΔ36BofA/SpoIVFA | Fragment *gfpΔ36bofA/spoIVFA* was amplified from pSO6 using primers SO-P82 and SO-P83. Fragment was joined to pSO40 digested with NotI and NheI using GA. | This study |
| pSO43 | Km^R^; T7-Pro-σ^K^(1-127)-His_6_/T7-cytTM-SpoIVFB-FLAG_2_-His_6_/T7-GFPΔ27+Δ85-87BofA/SpoIVFA | Fragment *gfpΔ27+Δ85-87bofA/spoIVFA* was amplified from pSO10 using primers SO-P82 and SO-P83. Fragment was joined to pSO40 digested with NotI and NheI using GA. | This study |
| pSO44 | Km^R^; T7-Pro-σ^K^(1-127)-His_6_/T7-cytTM-SpoIVFB-FLAG_2_-His_6_/T7-GFPΔ27BofA G40A/SpoIVFA | Fragment *gfpΔ27bofA G40A/spoIVFA* was amplified from pSO25 using primers SO-P82 and SO-P83. Fragment was joined to pSO40 digested with NotI and NheI using GA. | This study |
| pSO45 | Km^R^; T7-Pro-σ^K^(1-127)-His_6_/T7-cytTM-SpoIVFB-FLAG_2_-His_6_/T7-GFPΔ27BofA L44A/SpoIVFA | Fragment *gfpΔ27bofA L44A/spoIVFA* was amplified from pSO26 using primers SO-P82 and SO-P83. Fragment was joined to pSO40 digested with NotI and NheI using GA. | This study |
| pSO46 | Km^R^; T7-Pro-σ^K^(1-127)-His_6_/T7-cytTM-SpoIVFB-FLAG_2_-His_6_/T7-GFPΔ27BofA N48A/SpoIVFA | Fragment *gfpΔ27bofA N48A/spoIVFA* was amplified from pSO27 using primers SO-P82 and SO-P83. Fragment was joined to pSO40 digested with NotI and NheI using GA. | This study |
| pSO47 | Km^R^; T7-Pro-σ^K^(1-127)-His_6_/T7-cytTM-SpoIVFB-FLAG_2_-His_6_/T7-GFPΔ27BofA G51A/SpoIVFA | Fragment *gfpΔ27bofA G51A/spoIVFA* was amplified from pSO28 using primers SO-P82 and SO-P83. Fragment was joined to pSO40 digested with NotI and NheI using GA. | This study |
| pSO48 | Km^R^; T7-Pro-σ^K^(1-127)-His_6_/T7-cytTM-SpoIVFB-FLAG_2_-His_6_/T7-GFPΔ27BofA I60A/SpoIVFA | Fragment *gfpΔ27bofA I60A/spoIVFA* was amplified from pSO29 using primers SO-P82 and SO-P83. Fragment was joined to pSO40 digested with NotI and NheI using GA. | This study |
| pSO49 | Km^R^; T7-Pro-σ^K^(1-127)-His_6_/T7-cytTM-SpoIVFB-FLAG_2_-His_6_/T7-GFPΔ27BofA N61A/SpoIVFA | Fragment *gfpΔ27bofA N61A/spoIVFA* was amplified from pSO30 using primers SO-P82 and SO-P83. Fragment was joined to pSO40 digested with NotI and NheI using GA. | This study |
| pSO50 | Km^R^; T7-Pro-σ^K^(1-127)-His_6_/T7-cytTM-SpoIVFB-FLAG_2_-His_6_/T7-GFPΔ27BofA T64A/SpoIVFA | Fragment *gfpΔ27bofA T64A/spoIVFA* was amplified from pSO31using primers SO-P82 and SO-P83. Fragment was joined to pSO40 digested with NotI and NheI using GA. | This study |
| pSO51 | Km^R^; T7-Pro-σ^K^(1-127)-His_6_/T7-cytTM-SpoIVFB-FLAG_2_-His_6_/T7-GFPΔ27BofA G69A/SpoIVFA | Fragment *gfpΔ27bofA G69A/spoIVFA* was amplified from pSO32 using primers SO-P82 and SO-P83. Fragment was joined to pSO40 digested with NotI and NheI using GA. | This study |
| pSO52 | Km^R^; T7-Pro-σ^K^(1-127)-His_6_/T7-cytTM-SpoIVFB-FLAG_2_-His_6_/T7-GFPΔ27BofA L71A/SpoIVFA | Fragment *gfpΔ27bofA L71A/spoIVFA* was amplified from pSO33 using primers SO-P82 and SO-P83. Fragment was joined to pSO40 digested with NotI and NheI using GA. | This study |
| pSO53 | Km^R^; T7-Pro-σ^K^(1-127)-His_6_/T7-cytTM-SpoIVFB-FLAG_2_-His_6_/T7-GFPΔ27BofA G72A/SpoIVFA | Fragment *gfpΔ27bofA G72A/spoIVFA* was amplified from pSO34 using primers SO-P82 and SO-P83. Fragment was joined to pSO40 digested with NotI and NheI using GA. | This study |
| pSO54 | Km^R^; T7-Pro-σ^K^(1-127)-His_6_/T7-cytTM-SpoIVFB-FLAG_2_-His_6_/T7-GFPΔ27BofA P74A/SpoIVFA | Fragment *gfpΔ27bofA P74A/spoIVFA* was amplified from pSO35 using primers SO-P82 and SO-P83. Fragment was joined to pSO40 digested with NotI and NheI using GA. | This study |
| pSO55 | Km^R^; T7-Pro-σ^K^(1-127)-His_6_/T7-cytTM-SpoIVFB-FLAG_2_-His_6_/T7-GFPΔ27BofA G75A/SpoIVFA | Fragment *gfpΔ27bofA G75A/spoIVFA* was amplified from pSO36 using primers SO-P82 and SO-P83. Fragment was joined to pSO40 digested with NotI and NheI using GA. | This study |
| pSO56 | Km^R^; T7-Pro-σ^K^(1-127)-His_6_/T7-cytTM-SpoIVFB-FLAG_2_-His_6_/T7-GFPΔ27BofA L79A/SpoIVFA | Fragment *gfpΔ27bofA L79A/spoIVFA* was amplified from pSO37 using primers SO-P82 and SO-P83. Fragment was joined to pSO40 digested with NotI and NheI using GA. | This study |
| pSO57 | Km^R^; T7-Pro-σ^K^(1-127)-His_6_/T7-cytTM-SpoIVFB-FLAG_2_-His_6_/T7-GFPΔ27BofA I86A/SpoIVFA | Fragment *gfpΔ27bofA I86A/spoIVFA* was amplified from pSO38 using primers SO-P82 and SO-P83. Fragment was joined to pSO40 digested with NotI and NheI using GA. | This study |
| pSO58 | Km^R^; T7-Pro-σ^K^(1-127)-His_6_/T7-cytTM-SpoIVFB-FLAG_2_-His_6_/T7-GFPΔ27BofA I87A/SpoIVFA | Fragment *gfpΔ27bofA I87A/spoIVFA* was amplified from pSO39 using primers SO-P82 and SO-P83. Fragment was joined to pSO40 digested with NotI and NheI using GA. | This study |
| pSO60 | Km^R^; T7-Pro-σ^K^(1-127)-His_6_/T7-cytTM-SpoIVFB-FLAG_2_-His_6_/T7-GFPΔ27BofA H57A/SpoIVFA | pSO40 was subjected to SDM using primers SO-P91 and SO-P92, substituting H57A in GFPΔ27BofA. | This study |
| pSO61 | Km^R^; T7-Pro-σ^K^(1-127)-His_6_/T7-cytTM-SpoIVFB-FLAG_2_-His_6_/T7-GFPΔ27BofA P59A/SpoIVFA | pSO40 was subjected to SDM using primers SO-P93 and SO-P94, substituting P59A in GFPΔ27BofA. | This study |
| pSO62 | Km^R^; T7-Pro-σ^K^(1-127)-His_6_/T7-cytTM-SpoIVFB-FLAG_2_-His_6_/T7-GFPΔ27BofA I82A/SpoIVFA | pSO40 was subjected to SDM using primers SO-P95 and SO-P96, substituting I82A in GFPΔ27BofA. | This study |
| pSO63 | Km^R^; T7-Pro-σ^K^(1-127)-His_6_/T7-cytTM-SpoIVFB-FLAG_2_-His_6_/T7-GFPΔ27BofA F85A/SpoIVFA | pSO40 was subjected to SDM using primers SO-P97 and SO-P98, substituting F85A in GFPΔ27BofA. | This study |
| pSO64 | Km^R^; T7-Pro-σ^K^(1-127)-His_6_/T7-cytTM-SpoIVFB-FLAG_2_-His_6_/T7-GFPΔ27BofA | Fragment *gfpΔ27bofA* was amplified from pYZ46 using primers SO-P99 and SO-P100. Fragment was joined to pSO40 digested with NotI and NheI using GA. | This study |
| pSO65 | Km^R^; T7-Pro-σ^K^(1-127)-His_6_/T7-cytTM-SpoIVFB-FLAG_2_-His_6_/T7-SpoIVFA | Fragment *spoIVFA* was amplified from pYZ46 using primers SO-P82 and SO-P101. Fragment was joined to pSO40 digested with NotI and NheI using GA. | This study |
| pSO67 | Km^R^; T7-Pro-σ^K^(1-127)-His_6_/T7-cytTM-SpoIVFB-FLAG_2_-His_6_/T7-GFPΔ27BofA FII85-87AAA/SpoIVFA | pSO63 was subjected to SDM using primers SO-P106 and SO-P107, substituting FII85-87AAA in GFPΔ27BofA. | This study |
| pSO68 | Km^R^; T7-Pro-σ^K^(1-127)-His_6_/T7-cytTM-SpoIVFB-FLAG_2_-His_6_/T7-GFPΔ27BofA/Cys-less SpoIVFA | pSO40 was subjected to SDM using primers SO-P116 and SO-P117, substituting C77L and C82L in SpoIVFA. | This study |
| pSO69 | Km^R^; T7-Pro-σ^K^(1-127)-His_6_/T7-cytTM-SpoIVFB-FLAG_2_-His_6_/T7-GFP-G/S linker-Δ36BofA/SpoIVFA | pSO42 was subjected to SDM using primers SO-P108 and SO-P109, adding a nine-residue G/S linker (GGSGGSGGS) to GFPΔ36BofA. | This study |
| pSO70 | Km^R^; T7-Pro-σ^K^(1-127)-His_6_/T7-cytTM-SpoIVFB-FLAG_2_/T7-GFPΔ27BofA/SpoIVFA | Fragment *T7*-*gfpΔ27bofA/spoIVFA* was amplified from pYZ46 using primers SO-P82 and SO-P120. Vector pYZ2 was amplified using primers SO-P118 and SO-P119 (removing His_6_ from cytTM-SpoIVFB). Fragment was joined to pYZ2 using GA. | This study |
| pSO71 | Km^R^; T7-Pro-σ^K^(1-127)-His_6_/T7-cytTM-SpoIVFB-FLAG_2_-His_6_/T7-GFP C48S-Δ27BofA/Cys-less SpoIVFA | pSO68 (Cys-less SpoIVFA) was subjected to SDM using primers SO-P112 and SO-P113, substituting C48S in GFP. | This study |
| pSO72 | Km^R^; T7-Pro-σ^K^(1-127)-His_6_/T7-cytTM-SpoIVFB-FLAG_2_-His_6_/T7-GFP C48S C70S-Δ27BofA/Cys-less SpoIVFA | pSO71 was subjected to SDM using primers SO-P114 and SO-P115, substituting C70S in GFP. | This study |
| pSO73 | Km^R^; T7-Pro-σ^K^(1-127)-His_6_/T7-cytTM-SpoIVFB E44C-FLAG_2_/T7-GFPΔ27BofA/SpoIVFA | pSO70 was subjected to SDM using primers YZ1 and YZ2, substituting E44C in cytTM-SpoIVFB. | This study |
| pSO75 | Ap^R^ Sp^R^; P_s_*_poIID_*-GFPΔ27BofA | pDR95 was subjected to SDM using primers SO-P134 and SO-P135, replacing a HindIII site with a SphI site. | This study |
| pSO76 | Km^R^; T7-Cys-less Pro-σ^K^(1-127)-His_6_/T7-cytTM-SpoIVFB-FLAG_2_-His_6_ | pYZ2 was subjected to SDM using primers SO-P136 and SO-P137, substituting C109S in Pro-σ^K^(1-127). | This study |
| pSO78 | Ap^R^ Sp^R^; P*_bofA_*-GFPΔ27BofA | Fragment P*_bofA_* was amplified from *B. subtilis* strain PY79 DNA using primers SO-P132 and SO-P133. Fragment was digested with EcoRI and SphI, and ligated with EcoRI-SphI-digested pSO75. | This study |
| pSO79 | Km^R^; T7-Cys-less Pro-σ^K^(1-127)-His_6_/T7-single-Cys E44C cytTM-SpoIVFB-FLAG_2_-His_6_ | Fragment *spoIVFB* (single-Cys E44C) was amplified from pYZ40 using primers SO-P138 and SO-P139. Vector pSO76 was amplified using primers SO-P140 and SO-P141. Fragment was joined to pSO76 using GA. | This study |
| pSO80 | Km^R^; T7-Cys-less Pro-σ^K^(1-127)-His_6_/T7-single-Cys E44C cytTM- SpoIVFB-FLAG_2_-His_6_/T7-single-Cys GFPΔ27BofA/Cys-less SpoIVFA | Fragment *T7-gfpΔ27bofA/spoIVFA* (single-Cys GFPΔ27BofA/Cys-less SpoIVFA) was amplified from pSO72 using primers SO-P82 and SO-P90. Vector pSO79 was amplified using primers SO-P89 and SO-P80. Fragment was joined to pSO79 using GA. | This study |
| pSO82 | Km^R^; T7-Pro-σ^K^(1-127)/T7-cytTM-SpoIVFB E44C-FLAG_2_/T7-GFPΔ27BofA/SpoIVFA | pSO73 was subjected to SDM using primers SO-P148 and SO-P149, deleting His_6_ from Pro-σ^K^(1-127). | This study |
| pSO83 | Km^R^; T7-Cys-less Pro-σ^K^(1-127)-His_6_/T7-single-Cys V70C cytTM-SpoIVFB E44Q-FLAG_2_-His_6_/T7-single-Cys GFPΔ27BofA/Cys-less SpoIVFA | Fragment *spoIVFB* (single-Cys V70C) was amplified from pYZ77 using primers SO-P138 and SO-P139. Vector pSO80 was amplified using primers SO-P140 and SO-P141. Fragment was joined to pSO80 using GA. | This study |
| pSO84 | Km^R^; T7-Cys-less Pro-σ^K^(1-127)-His_6_/T7-single-Cys P135C cytTM-SpoIVFB E44Q-FLAG_2_-His_6_/T7-single-Cys GFPΔ27BofA/Cys-less SpoIVFA | Fragment *spoIVFB* (single-Cys P135C) was amplified from pYZ28 using primers SO-P138 and SO-P139. Vector pSO80 was amplified using primers SO-P140 and SO-P141. Fragment was joined to pSO80 using GA. | This study |
| pSO86 | Ap^R^ Sp^R^; P_b_*_ofA_*-GFPΔ27BofA N48A | Fragment *gfpΔ27bofA N48A* was amplified from pSO27 using primers SO-P142 and SO-P143. Fragment was joined to pSO78 digested with SphI and BamHI using GA. | This study |
| pSO87 | Ap^R^ Sp^R^; P_b_*_ofA_*-GFPΔ27BofA N61A | Fragment *gfpΔ27bofA* *N61A* was amplified from pSO30 using primers SO-P142 and SO-P143. Fragment was joined to pSO78 digested with SphI and BamHI using GA. | This study |
| pSO88 | Ap^R^ Sp^R^; P_b_*_ofA_*-GFPΔ27BofA T64A | pSO78 was subjected to SDM using primers SO-P60 and SO-P61, substituting T64A in GFPΔ27BofA. | This study |
| pSO90 | Km^R^; T7-Pro-σ^K^(1-127)-His_6_/T7-cytTM-SpoIVFB-FLAG_2_-His_6_/T7-MBPΔ27BofA/Cys-less SpoIVFA | Fragment of *mbp* was amplified from pYZ112 using primers SO-P156 and SO-P157. Vector pSO72 was amplified using primers SO-P158 and SO-P159. Fragment was joined to pSO72 using GA. | This study |
| pSO91 | Km^R^; T7-Cys-less Pro-σ^K^(1-127)-His_6_/T7- single-Cys E44C cytTM-SpoIVFB-FLAG_2_-His_6_/T7-MBPΔ27BofA/Cys-less SpoIVFA | Fragment *T7-mbpΔ27bofA/spoIVFA* (single-Cys MBPΔ27BofA/Cys-less SpoIVFA) was amplified from pSO90 using primers SO-P82 and SO-P90. Vector pSO80 was amplified using primers SO-P89 and SO-P80. Fragment was joined to pSO80 using GA. | This study |
| pSO92 | Km^R^; T7-Cys-less Pro-σ^K^(1-127)-His_6_/T7-single-Cys V70C cytTM-SpoIVFB E44Q-FLAG_2_-His_6_/T7-MBPΔ27BofA/Cys-less SpoIVFA | Fragment *T7-mbpΔ27bofA/spoIVFA* (single-Cys MBPΔ27BofA/Cys-less SpoIVFA) was amplified from pSO90 using primers SO-P82 and SO-P90. Vector pSO83 was amplified using primers SO-P89 and SO-P80. Fragment was joined to pSO83 using GA. | This study |
| pSO93 | Km^R^; T7-Cys-less Pro-σ^K^(1-127)-His_6_/T7-single-Cys P135C cytTM-SpoIVFB E44Q-FLAG_2_-His_6_/T7-MBPΔ27BofA/Cys-less SpoIVFA | Fragment *T7-mbpΔ27bofA/spoIVFA* (single-Cys MBPΔ27BofA/Cys-less SpoIVFA) was amplified from pSO90 using primers SO-P82 and SO-P90. Vector pSO84 was amplified using primers SO-P89 and SO-P80. Fragment was joined to pSO84 using GA. | This study |
| pSO94 | Km^R^; T7-Cys-less Pro-σ^K^(1-127)-His_6_/T7-Cys-less cytTM-SpoIVFB E44Q-FLAG_2_-His_6_/T7-MBPΔ27BofA/ Cys-less SpoIVFA | pSO91 was subjected to SDM using primers LK2691 and YZ11, substituting E44Q in cytTM-SpoIVFB. | This study |
| pSO96 | Km^R^; T7-Cys-less Pro-σ^K^(1-127)-His_6_/T7-Cys-less cytTM-SpoIVFB E44Q-FLAG_2_-His_6_ | pSO79 was subjected to SDM using primers LK2691 and YZ11, substituting E44Q in cytTM-SpoIVFB. | This study |
| pSO97 | Km^R^; T7-Pro-σ^K^(1-127)-His_6_/T7-cytTM-SpoIVFB-FLAG_2_-His_6_/T7-Cys-less MBPΔ27BofA/Cys-less SpoIVFA | pSO90 was subjected to SDM using primers SO-P152 and SO-P153, substituting C46S in MBPΔ27BofA. | This study |
| pSO110 | Km^R^; T7-Cys-less Pro-σ^K^(1-127)-His_6_/T7-single-Cys E44C cytTM-SpoIVFB-FLAG_2_-His_6_/T7-Cys-less MBPΔ27BofA/Cys-less SpoIVFA | Fragment *T7-mbpΔ27bofA/spoIVFA* (Cys-less MBPΔ27BofA/Cys-less SpoIVFA) was amplified from pSO97 using primers SO-P82 and SO-P90. Vector pSO80 was amplified using primers SO-P89 and SO-P80. Fragment was joined to pSO80 using GA. | This study |
| pSO111 | Km^R^; T7-Cys-less Pro-σ^K^(1-127)-His_6_/T7-single-Cys V70C cytTM-SpoIVFB E44Q-FLAG_2_-His_6_/T7-Cys-less MBPΔ27BofA/Cys-less SpoIVFA | Fragment *T7-mbpΔ27bofA/spoIVFA* (Cys-less MBPΔ27BofA/Cys-less SpoIVFA) was amplified from pSO97 using primers SO-P82 and SO-P90. Vector pSO83 was amplified using primers SO-P89 and SO-P80. Fragment was joined to pSO83 using GA. | This study |
| pSO112 | Km^R^; T7-Cys-less Pro-σ^K^(1-127)-His_6_/T7-single-Cys P135C cytTM-SpoIVFB E44Q-FLAG_2_-His_6_/T7-Cys-less MBPΔ27BofA/Cys-less SpoIVFA | Fragment *T7-mbpΔ27bofA/spoIVFA* (Cys-less MBPΔ27BofA and Cys-less SpoIVFA) was amplified from pSO97 using primers SO-P82 and SO-P90. Vector pSO84 was amplified using primers SO-P89 and SO-P80. Fragment was joined to pSO84 using GA. | This study |
| pSO113 | Km^R^; T7-single-Cys E42C Pro-σ^K^(1-127)-His_6_/T7-Cys-less cytTM-SpoIVFB E44Q-FLAG_2_-His_6_ | pSO96 was subjected to SDM using primers SO-P162 and SO-P163, substituting E42C in Pro-σ^K^(1-127). | This study |
| pSO114 | Km^R^; T7-Cys-less Pro-σ^K^(1-127)-His_6_/T7-single-Cys Y214C cytTM-SpoIVFB E44Q-FLAG_2_-His_6_ | pSO96 was subjected to SDM using primers SO-P170 and SO-P171, substituting Y214C in cytTM-SpoIVFB. | This study |
| pSO115 | Km^R^; T7-single-Cys A97C Pro-σ^K^(1-127)-His_6_/T7-Cys-less cytTM-SpoIVFB E44Q-FLAG_2_-His_6_ | pSO96 was subjected to SDM using primers SO-P176 and SO-P177, substituting A97C in Pro-σ^K^(1-127). | This study |
| pSO116 | Km^R^; T7-Cys-less Pro-σ^K^(1-127)-His_6_/T7-single-Cys V229C cytTM-SpoIVFB E44Q-FLAG_2_-His_6_ | pSO96 was subjected to SDM using primers SO-P180 and SO-P181, substituting V229C in cytTM-SpoIVFB. | This study |
| pSO117 | Km^R^; T7-single-Cys E42C Pro-σ^K^(1-127)-His_6_/T7-single-Cys R213C cytTM-SpoIVFB E44Q-FLAG_2_-His_6_ | pSO113 was subjected to SDM using primers SO-P168 and SO-P169, substituting R213C in cytTM-SpoIVFB. | This study |
| pSO118 | Km^R^; T7-single-Cys E42C Pro-σ^K^(1-127)-His_6_/T7-single-Cys Y214C cytTM-SpoIVFB E44Q-FLAG_2_-His_6_ | pSO113 was subjected to SDM using primers SO-P170 and SO-P171, substituting Y214C in cytTM-SpoIVFB. | This study |
| pSO119 | Km^R^; T7-single-Cys E42C Pro-σ^K^(1-127)-His_6_/T7-single-Cys Y215C cytTM-SpoIVFB E44Q-FLAG_2_-His_6_ | pSO113 was subjected to SDM using primers SO-P172 and SO-P173, substituting Y215C in cytTM-SpoIVFB. | This study |
| pSO120 | Km^R^; T7-single-Cys L41C Pro-σ^K^(1-127)-His_6_/T7-single-Cys Y214C cytTM-SpoIVFB E44Q-FLAG_2_-His_6_ | pSO114 was subjected to SDM using primers SO-P186 and SO-P187, substituting L41C in Pro-σ^K^(1-127). | This study |
| pSO121 | Km^R^; T7-single-Cys L43C Pro-σ^K^(1-127)-His_6_/T7-single-Cys Y214C cytTM-SpoIVFB E44Q-FLAG_2_-His_6_ | pSO114 was subjected to SDM using primers SO-P164 and SO-P165, substituting L43C in Pro-σ^K^(1-127). | This study |
| pSO122 | Km^R^; T7-single-Cys A97C Pro-σ^K^(1-127)-His_6_/T7-single-Cys S228C cytTM-SpoIVFB E44Q-FLAG_2_-His_6_ | pSO115 was subjected to SDM using primers SO-P188 and SO-P189, substituting S228C in cytTM-SpoIVFB. | This study |
| pSO123 | Km^R^; T7-single-Cys A97C Pro-σ^K^(1-127)-His_6_/T7-single-Cys V229C cytTM-SpoIVFB E44Q-FLAG_2_-His_6_ | pSO115 was subjected to SDM using primers SO-P180 and SO-P181, substituting V229C in cytTM-SpoIVFB. | This study |
| pSO124 | Km^R^; T7-single-Cys A97C Pro-σ^K^(1-127)-His_6_/T7-single-Cys K230C cytTM-SpoIVFB E44Q-FLAG_2_-His_6_ | pSO115 was subjected to SDM using primers SO-P182 and SO-P183, substituting K230C in cytTM-SpoIVFB. | This study |
| pSO125 | Km^R^; T7-single-Cys S96C Pro-σ^K^(1-127)-His_6_/T7-single-Cys V229C cytTM-SpoIVFB E44Q-FLAG_2_-His_6_ | pSO116 was subjected to SDM using primers SO-P174 and SO-P175, substituting S96C in Pro-σ^K^(1-127). | This study |
| pSO126 | Km^R^; T7-single-Cys G98C Pro-σ^K^(1-127)-His_6_/T7-single-Cys V229C cytTM-SpoIVFB E44Q-FLAG_2_-His_6_ | pSO116 was subjected to SDM using primers SO-P196 and SO-P197, substituting G98C in Pro-σ^K^(1-127). | This study |
| pSO127 | Km^R^; T7-single-Cys L41C Pro-σ^K^(1-127)-His_6_/T7-single-Cys Y214C cytTM-SpoIVFB E44Q-FLAG_2_-His_6_/T7-Cys-less MBPΔ27BofA/Cys-less SpoIVFA | Fragment *T7-mbpΔ27bofA/spoIVFA* (Cys-less MBPΔ27BofA/Cys-less SpoIVFA) was amplified from pSO112 using primers SO-P82 and SO-P90. Vector pSO120 was amplified using primers SO-P89 and SO-P80. Fragment was joined to pSO120 using GA. | This study |
| pSO128 | Km^R^; T7-single-Cys K24C Pro-σ^K^(1-127)-His_6_/T7-single-Cys E44C cytTM-SpoIVFB-FLAG_2_-His_6_ | pSO79 was subjected to SDM using primers LK2473 and LK2474, substituting K24C in Pro-σ^K^(1-127). | This study |
| pSO130 | Km^R^; T7-single-Cys A97C Pro-σ^K^(1-127)-His_6_/T7-single-Cys A231C cytTM-SpoIVFB E44Q-FLAG_2_-His_6_ | pSO115 was subjected to SDM using primers SO-P184 and SO-P185, substituting A231C in cytTM-SpoIVFB. | This study |
| pSO131 | Km^R^; T7-single-Cys K24C Pro-σ^K^(1-127)-His_6_/T7-single-Cys E44C cytTM-SpoIVFB-FLAG_2_-His_6_/T7-Cys-less MBPΔ27BofA/Cys-less SpoIVFA | pSO110 was subjected to SDM using primers LK2473 and LK2474, substituting K24C in Pro-σ^K^(1-127). | This study |
| pSO132 | Km^R^; T7-single-Cys K24C Pro-σ^K^(1-127)-His_6_/T7-single-Cys V70C cytTM-SpoIVFB E44Q-FLAG_2_-His_6_/T7-Cys-less MBPΔ27BofA/Cys-less SpoIVFA | pSO111 was subjected to SDM using primers LK2473 and LK2474, substituting K24C in Pro-σ^K^(1-127). | This study |
| pSO133 | Km^R^; T7-single-Cys A97C Pro-σ^K^(1-127)-His_6_/T7-single-Cys A231C cytTM-SpoIVFB E44Q-FLAG_2_-His_6_/T7-Cys-less MBPΔ27BofA/Cys-less SpoIVFA | Fragment *T7-mbpΔ27bofA/spoIVFA* (Cys-less MBPΔ27BofA/Cys-less SpoIVFA) was amplified from pSO112 using primers SO-P82 and SO-P90. Vector pSO130 was amplified using primers SO-P89 and SO-P80. Fragment was joined to pSO130 using GA. | This study |
| pSO134 | Km^R^; T7-single-Cys K24C Pro-σ^K^(1-127)-His_6_/T7-single-Cys V70C cytTM-SpoIVFB E44Q-FLAG_2_-His_6_ | Fragment *spoIVFB* (single-Cys V70C) was amplified from pSO111 using primers SO-P138 and SO-P139. Vector pSO128 was amplified from primers SO-P140 and SO-P141. Fragment was joined to pSO128 using GA. | This study |
| pSO136 | Km^R^; T7-Cys-less Pro-σ^K^(1-127)-His_6_/T7-single-Cys V70C cytTM-SpoIVFB E44Q-FLAG_2_-His_6_ | Fragment of *spoIVFB* (single-Cys V70C) was amplified from pSO111 using primers SO-P138 and SO-P139. Vector pSO79 was amplified fromprimers SO-P140 and SO-P141. Fragment was joined to pSO79 using GA. | This study |
| pSO139 | Km^R^; T7-Cys-less Pro-σ^K^(1-127)-His_6_/T7-Cys-less cytTM-SpoIVFB E44Q-FLAG_2_-His_6_/T7-Cys-less MBPΔ27BofA/Cys-less SpoIVFA | pSO94 was subjected to SDM using primers SO-P152 and SO-P153, substituting C46S in MBPΔ27BofA. | This study |
| pSO141 | Km^R^; T7-Pro-σ^K^(1-127)-His_6_/T7-cytTM-SpoIVFB M30C-FLAG_2_-His_6_ | pYZ2 was subjected to SDM using primers SO-P202 and SO-P203, substituting M30C in cytTM-SpoIVFB. | This study |
| pSO142 | Km^R^; T7-Pro-σ^K^(1-127)-His_6_/T7-cytTM-SpoIVFB-FLAG_2_-His_6_/T7-GFPΔ27BofA L62C/SpoIVFA | pSO40 was subjected to SDM using primers SO-P204 and SO-P205, substituting L62C in GFPΔ27BofA. | This study |
| pSO143 | Km^R^; T7-Pro-σ^K^(1-127)-His_6_/T7-cytTM-SpoIVFB-FLAG_2_-His_6_/T7-GFPΔ27BofA V63C/SpoIVFA | pSO40 was subjected to SDM using primers SO-P206 and SO-P207, substituting V63C in GFPΔ27BofA. | This study |
| pSO144 | Km^R^; T7-Cys-less Pro-σ^K^(1-127)-His_6_/T7-single-Cys M30C cytTM-SpoIVFB E44Q-FLAG_2_-His_6_/T7-Cys-less MBPΔ27BofA/Cys-less SpoIVFA | pSO139 was subjected to SDM using primers SO-P209 and SO-P210, substituting M30C in cytTM-SpoIVFB. | This study |
| pS0147 | Km^R^; T7-Cys-less Pro-σ^K^(1-127)-His_6_/T7-single-Cys M30C cytTM-SpoIVFB E44Q-FLAG_2_-His_6_/T7-single-Cys L62C MBPΔ27BofA/Cys-less SpoIVFA | pSO144 was subjected to SDM using primers SO-P204 and SO-P205, substituting L62C in MBPΔ27BofA. | This study |
| pSO148 | Km^R^; T7-Cys-less Pro-σ^K^(1-127)-His_6_/T7-single-Cys M30C cytTM-SpoIVFB E44Q-FLAG_2_-His_6_/T7-single-Cys V63C MBPΔ27BofA/Cys-less SpoIVFA | pSO144 was subjected to SDM using primers SO-P206 and SO-P207, substituting V63C in MBPΔ27BofA. | This study |
| pSO149 | Km^R^; T7-Pro-σ^K^(1-127)-His_6_/T7-cytTM-SpoIVFB E44C/T7-GFPΔ27BofA/SpoIVFA | pSO73 was subjected to SDM using primers SO-P211 and SO-P212, deleting FLAG_2_ from cytTM-SpoIVFB. | This study |
| pSO157 | Km^R^; T7-Pro-σ^K^(1-127) L41C-His_6_/T7-cytTM-SpoIVFB-FLAG_2_-His_6_ | pYZ2 was subjected to SDM using primers SO-P186 and SO-P187, substituting L41C in Pro-σ^K^(1-127). | This study |
| pSO158 | Km^R^; T7-Pro-σ^K^(1-127) A97C-His_6_/T7-cytTM-SpoIVFB-FLAG_2_-His_6_ | pYZ2 was subjected to SDM using primers SO-P176 and SO-P177, substituting A97C in Pro-σ^K^(1-127). | This study |
| pSO159 | Km^R^; T7-Pro-σ^K^(1-127)-His_6_/T7-cytTM-SpoIVFB Y214C-FLAG_2_-His_6_ | pYZ2 was subjected to SDM using primers SO-P170 and SO-P171, substituting Y214C in cytTM-SpoIVFB. | This study |
| pSO160 | Km^R^; T7-Pro-σ^K^(1-127)-His_6_/T7-cytTM-SpoIVFB A231C-FLAG_2_-His_6_ | pYZ2 was subjected to SDM using primers SO-P184 and SO-P185, substituting Y231C in cytTM-SpoIVFB. | This study |
| pSO163 | Km^R^; T7-single-Cys F18C Pro-σ^K^(1-127)-His_6_/T7-single-Cys E44C cytTM-SpoIVFB-FLAG_2_-His_6_/T7-Cys-less MBPΔ27BofA/Cys-less SpoIVFA | pSO110 was subjected to SDM using primers LK2465 and LK2466, substituting F18C in Pro-σ^K^(1-127). | This study |
| pSO164 | Km^R^; T7-single-Cys F18C Pro-σ^K^(1-127)-His_6_/T7-single-Cys V70C cytTM-SpoIVFB E44Q-FLAG_2_-His_6_/T7-Cys-less MBPΔ27BofA/Cys-less SpoIVFA | pSO111 was subjected to SDM using primers LK2465 and LK2466, substituting F18C in Pro-σ^K^(1-127). | This study |
| pSO165 | Km^R^; T7-single-Cys V20C Pro-σ^K^(1-127)-His_6_/T7-single-Cys E44C cytTM-SpoIVFB-FLAG_2_-His_6_/T7-Cys-less MBPΔ27BofA/Cys-less SpoIVFA | pSO110 was subjected to SDM using primers LK2467 and LK2468, substituting V20C in Pro-σ^K^(1-127). | This study |
| pSO166 | Km^R^; T7-single-Cys S21C Pro-σ^K^(1-127)-His_6_/T7-single-Cys E44C cytTM-SpoIVFB-FLAG_2_-His_6_/T7-Cys-less MBPΔ27BofA/Cys-less SpoIVFA | pSO110 was subjected to SDM using primers SO-P218 and SO-P219, substituting S21C in Pro-σ^K^(1-127). | This study |
| pSO167 | Km^R^; T7-single-Cys F18C Pro-σ^K^(1-127)-His_6_/T7-single-Cys E44C cytTM-SpoIVFB-FLAG_2_-His_6_ | pSO79 was subjected to SDM using primers LK2465 and LK2466, substituting F18C in Pro-σ^K^(1-127). | This study |
| pSO168 | Km^R^; T7-single-Cys F18C Pro-σ^K^(1-127)-His_6_/T7-single-Cys V70C cytTM-SpoIVFB E44Q-FLAG_2_-His_6_ | pSO136 was subjected to SDM using primers LK2465 and LK2466, substituting F18C in Pro-σ^K^(1-127). | This study |
| pSO169 | Km^R^; T7-single Cys V20C Pro-σ^K^(1-127)-His_6_/T7-single-Cys E44C cytTM- SpoIVFB-FLAG_2_-His_6_ | pSO79 was subjected to SDM using primers LK2467 and LK2468, substituting V20C in Pro-σ^K^(1-127). | This study |
| pSO170 | Km^R^; T7-single Cys S21C Pro-σ^K^(1-127)-His_6_/T7-single-Cys E44C cytTM- SpoIVFB-FLAG_2_-His_6_ | pSO79 was subjected to SDM using primers SO-P218 and SO-P219, substituting S21C in Pro-σ^K^(1-127). | This study |
| pSO181 | Km^R^; T7-Cys-less Pro-σ^K^(1-127)-His_6_/T7-Cys-less cytTM-SpoIVFB E44Q-FLAG_2_-His_6_/T7-single-Cys I56C MBPΔ27BofA/Cys-less SpoIVFA | pSO139 was subjected to SDM using primers SO-P222 and SO-P223, substituting I56C in MBPΔ27BofA. | This study |
| pSO182 | Km^R^; T7-Cys-less Pro-σ^K^(1-127)-His_6_/T7-Cys-less cytTM-SpoIVFB E44Q-FLAG_2_-His_6_/T7-single-Cys H57C MBPΔ27BofA/Cys-less SpoIVFA | pSO139 was subjected to SDM using primers SO-P224 and SO-P225, substituting H57C in MBPΔ27BofA. | This study |
| pSO183 | Km^R^; T7-Cys-less Pro-σ^K^(1-127)-His_6_/T7-Cys-less cytTM-SpoIVFB E44Q-FLAG_2_-His_6_/T7-single-Cys G40C MBPΔ27BofA/Cys-less SpoIVFA | pSO139 was subjected to SDM using primers SO-P226 and SO-P227, substituting G40C in MBPΔ27BofA. | This study |
| pSO184 | Km^R^; T7-Cys-less Pro-σ^K^(1-127)-His_6_/T7-Cys-less cytTM-SpoIVFB E44Q-FLAG_2_-His_6_/T7-single-Cys A41C MBPΔ27BofA/Cys-less SpoIVFA | pSO139 was subjected to SDM using primers SO-P228 and SO-P229, substituting A41C in MBPΔ27BofA. | This study |
| pSO186 | Km^R^; T7-Cys-less Pro-σ^K^(1-127)-His_6_/T7-single-Cys A32C cytTM-SpoIVFB E44Q-FLAG_2_-His_6_/T7-single-Cys I56C MBPΔ27BofA/Cys-less SpoIVFA | pSO181 was subjected to SDM using primers SO-P232 and SO-P233, substituting A32C in cytTM-SpoIVFB. | This study |
| pSO187 | Km^R^; T7-Cys-less Pro-σ^K^(1-127)-His_6_/T7-single-Cys L33C cytTM-SpoIVFB E44Q-FLAG_2_-His_6_/T7-single-Cys I56C MBPΔ27BofA/Cys-less SpoIVFA | pSO181 was subjected to SDM using primers SO-P234 and SO-P235, substituting L33C in cytTM-SpoIVFB. | This study |
| pSO188 | Km^R^; T7-Cys-less Pro-σ^K^(1-127)-His_6_/T7-single-Cys Q181C cytTM-SpoIVFB E44Q-FLAG_2_-His_6_/T7-single-Cys H57C MBPΔ27BofA/Cys-less SpoIVFA | pSO182 was subjected to SDM using primers SO-P236 and SO-P237, substituting Q181C in cytTM-SpoIVFB. | This study |
| pSO189 | Km^R^; T7-Cys-less Pro-σ^K^(1-127)-His_6_/T7-single-Cys V86C cytTM-SpoIVFB E44Q-FLAG_2_-His_6_/T7-single-Cys G40C MBPΔ27BofA/Cys-less SpoIVFA | pSO183 was subjected to SDM using primers SO-P238 and SO-P239, substituting V86C in cytTM-SpoIVFB. | This study |
| pSO190 | Km^R^; T7-Cys-less Pro-σ^K^(1-127)-His_6_/T7-single-Cys V86C cytTM-SpoIVFB E44Q-FLAG_2_-His_6_/T7-single-Cys A41C MBPΔ27BofA/Cys-less SpoIVFA | pSO184 was subjected to SDM using primers SO-P238 and SO-P239, substituting V86C in cytTM-SpoIVFB. | This study |
| pSO192 | Ap^R^; T7-SpoIVB S378A-His_6_ | pZR53 was subjected to SDM using primers SO-P244 and SO-P245, substituting S378A in SpoIVB. | This study |
| pSO193 | Km^R^; T7-Pro-σ^K^(1-127)-His_6_/T7-cytTM- SpoIVFB F66A-FLAG_2_-His_6_/T7-GFPΔ27BofA/SpoIVFA | pSO40 was subjected to SDM using primers SO-P242 and SO-P243, substituting F66A in cytTM-SpoIVFB. | This study |
| pSO203 | Km^R^; T7-Pro-σ^K^(1-127)-His_6_/T7-cytTM-SpoIVFB-FLAG_2_-His_6_/T7-GFPΔ27BofA N48D/SpoIVFA | pSO40 was subjected to SDM using primers SO-P251 and SO-P252, substituting N48D in GFPΔ27BofA. | This study |
| pSO211 | Km^R^; T7-Pro-σ^K^-His_6_/T7-cytTM- SpoIVFB E44C-FLAG_2_/T7-GFPΔ27BofA/SpoIVFA | Fragment of 5’ end of *pro-*σ*^K^-His_6_* was amplified from pETproσ^K^ using primers SO-P260 and SO-P270. Vector pSO73 was amplified using primers SO-P257 and SO-P269. Fragment was joined to pSO73 using GA. | This study |
| pSO212 | Ap^R^; T7-BofA/SpoIVFA | Fragment of full-length *bofA* was amplified from *B. subtilis* strain PY79 DNA using primers SO-P271 and SO-P272. Fragment was used as template with primers SO-P277 and SO-P276 to add regions of homology to pYZ46. Vector pYZ46 was amplified with primers SO-P273 and SO-P280. Fragment with homology was joined to pYZ46 using GA. | This study |
| pSO213 | Km^R^; T7-Pro-σ^K^(1-127)-His_6_/T7-cytTM-SpoIVFB-FLAG_2_-His_6_/T7-BofA/SpoIVFA | Fragment *T7-bofA/spoIVFA* was amplified from pSO212 using primers SO-P82 and SO-P90. Vector pYZ2 was amplified using primers SO-P89 and SO-P80. Fragment was joined to pYZ2 using GA. | This study |
| pSO215 | Km^R^; T7-Pro-σ^K^(1-127)-His_6_/T7-cytTM-SpoIVFB E44C-FLAG_2_/T7-BofA/SpoIVFA | Fragment *spoIVFB/T7-bofA/spoIVFA* was made by OEPCR^c^. Fragment #1 (*T7-bofA/spoIVFA*) was amplified from pSO212 using primers SO-P120 and SO-P286. Fragment #2 (*spoIVFB*) was amplified from pSO70 using primers SO-P288 and SO-P293. Fragments #1 and #2 were used as template for OEPCR using primers SO-P294 and SO-P286 (removing His_6_ from cytTM-SpoIVFB). Vector pSO73 was amplified using primers SO-P287 and SO-P295. Product from OEPCR was joined to pSO73 using GA. | This study |
| pSO216 | Km^R^; T7-Pro-σ^K^(1-127)/T7-cytTM-SpoIVFB E44C-FLAG_2_/T7-BofA/SpoIVFA | pSO215 was subjected to SDM using primers SO-P148 and SO-P149, deleting His_6_ from Pro-σ^K^(1-127). | This study |
| pSO217 | Km^R^; T7-Pro-σ^K^(1-127)-His_6_/T7-cytTM-SpoIVFB E44C/T7-BofA/SpoIVFA | pSO215 was subjected to SDM using primers SO-P211 and SO-P212, deleting FLAG_2_ from cytTM-SpoIVFB. | This study |
| pSO220 | Km^R^; T7-Pro-σ^K^/T7-cytTM-SpoIVFB E44C-FLAG_2_/T7-GFPΔ27BofA/SpoIVFA | pSO211 was subjected to SDM using primers SO-P148 and SO-P149, deleting His_6_ from Pro-σ^K^. | This study |
| pSO221 | Km^R^; T7-Pro-σ^K^-His_6_/T7-cytTM- SpoIVFB E44C/T7-GFPΔ27BofA/SpoIVFA | pSO211 was subjected to SDM using primers: SO-P211 and SO-P212, deleting FLAG_2_ from cytTM-SpoIVFB. | This study |
| pSO224 | Km^R^; T7-Cys-less Pro-σ^K^(1-127)-His_6_/T7-cytTM-SpoIVFB-FLAG_2_-His_6_/T7-BofA/SpoIVFA | pSO213 was subjected to SDM using primers SO-P136 and SO-P137, substituting C109S in Pro-σ^K^(1-127). | This study |
| pSO225 | Km^R^; T7-Cys-less Pro-σ^K^(1-127)-His_6_/T7-cytTM-SpoIVFB-FLAG_2_-His_6_/T7-BofA/Cys-less SpoIVFA | pSO224 was subjected to SDM using primers SO-P116 and SO-P117, substituting C77L and C82L in SpoIVFA. | This study |
| pSO226 | Km^R^; T7-Cys-less Pro-σ^K^(1-127)-His_6_/T7-single-Cys E44C cytTM-SpoIVFB-FLAG_2_-His_6_/T7-BofA/Cys-less SpoIVFA | Fragment of *spoIVFB* (single-Cys E44C) was amplified from pSO91 using primers SO-P138 and SO-P139. Vector pSO225 was amplified using primers SO-P140 and SO-P141. Fragment was joined to pSO225 using GA. | This study |
| pSO227 | Km^R^; T7-Cys-less Pro-σ^K^(1-127)-His_6_/T7-single-Cys V70C cytTM-SpoIVFB E44Q-FLAG_2_-His_6_/T7-BofA/Cys-less SpoIVFA | Fragment of *spoIVFB* (single-Cys V70C) was amplified from pSO92 using primers SO-P138 and SO-P139. Vector pSO225 was amplified using primers SO-P140 and SO-P141. Fragment was joined to pSO225 using GA. | This study |
| pSO228 | Km^R^; T7-Cys-less Pro-σ^K^(1-127)-His_6_/T7-single-Cys P135C cytTM-SpoIVFB E44Q-FLAG_2_-His_6_/T7-BofA/Cys-less SpoIVFA | Fragment of *spoIVFB* (single-Cys P135C) was amplified from pSO93 using primers SO-P138 and SO-P139. Vector pSO225 was amplified using primers SO-P140 and SO-P141. Fragment was joined to pSO225 using GA. | This study |
| pSO229 | Km^R^; T7-Cys-less Pro-σ^K^(1-127)-His_6_/T7-single-Cys E44C cytTM-SpoIVFB-FLAG_2_-His_6_/T7-Cys-less BofA/Cys-less SpoIVFA | pSO226 was subjected to SDM using primers SO-P152 and SO-P153, substituting C46S in BofA. | This study |
| pSO230 | Km^R^; T7-Cys-less Pro-σ^K^(1-127)-His_6_/T7-single-Cys V70C cytTM-SpoIVFB E44Q-FLAG_2_-His_6_/T7-Cys-less BofA/Cys-less SpoIVFA | pSO227 was subjected to SDM using primers SO-P152 and SO-P153, substituting C46S in BofA. | This study |
| pSO231 | Km^R^; T7-Cys-less Pro-σ^K^(1-127)-His_6_/T7-single-Cys P135C cytTM-SpoIVFB E44Q-FLAG_2_-His_6_/T7-Cys-less BofA/Cys-less SpoIVFA | pSO228 was subjected to SDM using primers SO-P152 and SO-P153, substituting C46S in BofA. | This study |
| pSO232 | Ap^R^; T7-GFPΔ27BofA/SpoIVFA/SpoIVB S378A | Fragment *spoIVFA/spoIVB* *S378A* was made by OEPCR. Fragment #1 (*spoIVB S378A*) was amplified from pSO192 using primers SO-P302 and SO-P282. Fragment #2 (*spoIVFA*) was amplified from pYZ46 using primers SO-P289 and SO-P303. Fragments #1 and #2 were used as template for OEPCR using primers SO-P304 and SO-305. Vector pYZ46 was amplfied using primers SO-P306 and SO-P307. Product from OEPCR was joined to pYZ46 using GA. | This study |
| pSO233 | Ap^R^; T7-GFPΔ27BofA/SpoIVFA/SpoIVB | Fragment *spoIVFA/spoIVB* was made by OEPCR. Fragment #1 (*spoIVB*) was amplified from pZR53 using primers SO-P302 and SO-P282. Fragment #2 (*spoIVFA*) was amplified from pYZ46 using primers SO-P289 and SO-P303. Fragments #1 and #2 were used as template for OEPCR using primers SO-P304 and SO-305. Vector pYZ46 was amplified using primers SO-P306 and SO-P307. Product from OEPCR was joined to pYZ46 using GA. | This study |
| pSO234 | Km^R^; T7-single-Cys V20C Pro-σ^K^(1-127)-His_6_/T7-single-Cys E44C cytTM-SpoIVFB-FLAG_2_-His_6_/T7-Cys-less BofA/Cys-less SpoIVFA | pSO229 was subjected to SDM using primers LK2467 and LK2468, substituting V20C in Pro-σ^K^(1-127). | This study |
| pSO235 | Km^R^; T7-single-Cys S21C Pro-σ^K^(1-127)-His_6_/T7-single-Cys E44C cytTM-SpoIVFB-FLAG_2_-His_6_/T7-Cys-less BofA/Cys-less SpoIVFA | pSO229 was subjected to SDM using primers SO-P218 and SO-P219, substituting S21C in Pro-σ^K^(1-127). | This study |
| pSO236 | Km^R^; T7-single-Cys F18C Pro-σ^K^(1-127)-His_6_/T7-single-Cys V70C cytTM-SpoIVFB E44Q-FLAG_2_-His_6_/T7-Cys-less BofA/Cys-less SpoIVFA | pSO230 was subjected to SDM using primers LK2465 and LK2466, substituting F18C in Pro-σ^K^(1-127). | This study |
| pSO237 | Km^R^; T7-single-Cys K24C Pro-σ^K^(1-127)-His_6_/T7-single-Cys V70C cytTM-SpoIVFB E44Q-FLAG_2_-His_6_/T7-Cys-less BofA/Cys-less SpoIVFA | pSO230 was subjected to SDM using primers LK2473 and LK2474, substituting K24C in Pro-σ^K^(1-127). | This study |
| pSO238 | Km^R^; T7-single-Cys F18C Pro-σ^K^(1-127)-His_6_/T7-single-Cys E44C cytTM-SpoIVFB-FLAG_2_-His_6_/T7-Cys-less BofA/Cys-less SpoIVFA | pSO229 was subjected to SDM using primers LK2465 and LK2466, substituting F18C in Pro-σ^K^(1-127). | This study |
| pSO239 | Km^R^; T7-single-Cys K24C Pro-σ^K^(1-127)-His_6_/T7-single-Cys E44C cytTM-SpoIVFB-FLAG_2_-His_6_/T7-Cys-less BofA/Cys-less SpoIVFA | pSO229 was subjected to SDM using primers LK2473 and LK2474, substituting K24C in Pro-σ^K^(1-127). | This study |
| pSO240 | Km^R^; T7-Pro-σ^K^(1-127)-His_6_/T7-cytTM-SpoIVFB-FLAG_2_-His_6_/T7-GFPΔ27BofA/SpoIVFA/SpoIVB S378A | Fragment *gfpΔ27bofA/spoIVFA/spoIVB* *S378A* was amplified from pSO232 using primers SO-P284 and SO-P285. Vector pSO40 was amplified using primers SO-P80 and SO-P283. Fragment was joined to pSO40 using GA. | This study |
| pSO241 | Km^R^; T7-Pro-σ^K^(1-127)-His_6_/T7-cytTM-SpoIVFB-FLAG_2_-His_6_/T7-GFPΔ27BofA/SpoIVFA/SpoIVB | Fragment *gfpΔ27bofA/spoIVFA/spoIVB* was amplified from pSO233 using primers SO-P284 and SO-P285. Vector pSO40 was amplified using primers SO-P80 and SO-P283. Fragment was joined to pSO40 using GA. | This study |
| pSO242 | Km^R^; T7-Cys-less Pro-σ^K^(1-127)-His_6_/T7-Cys-less cytTM-SpoIVFB E44Q-FLAG_2_-His_6_/T7-Cys-less BofA/Cys-less SpoIVFA | pSO229 was subjected to SDM using primers LK2691 and YZ11, substituting E44Q in cytTM-SpoIVFB. | This study |
| pSO243 | Km^R^; T7-Cys-less Pro-σ^K^(1-127)-His_6_/T7-single-Cys Y214C cytTM-SpoIVFB E44Q-FLAG_2_-His_6_/T7-Cys-less BofA/Cys-less SpoIVFA | pSO242 was subjected to SDM using primers SO-P170 and SO-P171, substituting Y214C in cytTM-SpoIVFB. | This study |
| pSO244 | Km^R^; T7-Cys-less Pro-σ^K^(1-127)-His_6_/T7-single-Cys A231C cytTM-SpoIVFB E44Q-FLAG_2_-His_6_/T7-Cys-less BofA/Cys-less SpoIVFA | pSO242 was subjected to SDM using primers SO-P184 and SO-P185, substituting A231C in cytTM-SpoIVFB. | This study |
| pSO245 | Km^R^; T7-single-Cys L41C Pro-σ^K^(1-127)-His_6_/T7-single-Cys Y214C cytTM-SpoIVFB E44Q-FLAG_2_-His_6_/T7-Cys-less BofA/Cys-less SpoIVFA | pSO243 was subjected to SDM using primers SO-P186 and SO-P187, substituting L41C in Pro-σ^K^(1-127). | This study |
| pSO246 | Km^R^; T7-single-Cys A97C Pro-σ^K^(1-127)-His_6_/T7-single-Cys A231C cytTM-SpoIVFB E44Q-FLAG_2_-His_6_/T7-Cys-less BofA/Cys-less SpoIVFA | pSO244 was subjected to SDM using primers SO-P176 and SO-P177, substituting A97C in Pro-σ^K^(1-127). | This study |
| pSO247 | Ap^R^; T7-GFPΔ27BofA/SpoIVFA/T7-SpoIVB S378A | Fragment *spoIVFA/T7-spoIVB S378A* was made by OEPCR. Fragment #1 (*T7-spoIVB S378A*) was amplified from pSO192 using primers SO-P248 and SO-P282. Fragment #2 (*spoIVFA*) was amplified from pYZ46 using primers SO-P289 and SO-P290. Fragments #1 and #2 were used as template for OEPCR using primers SO-P304 and SO-305. Vector pYZ46 was amplified using primers SO-P306 and SO-P307. Product from OEPCR was joined to pYZ46 using GA. | This study |
| pSO248 | Ap^R^; T7-BofA/SpoIVFA/SpoIVB | Fragment *spoIVFA/spoIVB* was made by OEPCR. Fragment #1 (*spoIVB*) was amplified from pZR53 using primers SO-P302 and SO-P282. Fragment #2 (*spoIVFA*) was amplified from pYZ46 using primers SO-P289 and SO-P303. Fragments #1 and #2 were used as template for OEPCR using primers SO-P304 and SO-305. Vector pSO212 was amplified using SO-P306 and SO-P307. Product from OEPCR was joined to pSO212 using GA. | This study |
| pSO249 | Ap^R^; T7-BofA/SpoIVFA/SpoIVB S378A | Fragment *spoIVFA/spoIVB* *S378A* was made by OEPCR. Fragment #1 (*spoIVB S378A*) was amplified from pSO192 using primers SO-P302 and SO-P282. Fragment #2 (*spoIVFA*) was amplified from pYZ46 using primers SO-P289 and SO-P303. Fragments #1 and #2 were used as template for OEPCR using primers SO-P304 and SO-305. Vector pSO212 was amplified using SO-P306 and SO-P307. Product from OEPCR was joined to pSO212 using GA. | This study |
| pSO250 | Ap^R^; T7-BofA/SpoIVFA/T7-SpoIVB S378A | Fragment *spoIVFA/T7-spoIVB* *S378A* was made by OEPCR. Fragment #1 (*T7-spoIVB S378A*) was amplified from pSO192 using primers SO-P248 and SO-P282. Fragment #2 (*spoIVFA*) was amplified from pYZ46 using primers SO-P289 and SO-P290. Fragments #1 and #2 were used as template for OEPCR using primers SO-P304 and SO-305. Vector pSO212 was amplified using primers SO-P306 and SO-P307. Product from OEPCR was joined to pYZ46 using GA. | This study |
| pSO251 | Km^R^; T7-Pro-σ^K^(1-127)-His_6_/T7-cytTM-SpoIVFB-FLAG_2_-His_6_/T7-GFPΔ27BofA/SpoIVFA/T7-SpoIVB S378A | Fragment *gfpΔ27bofA/spoIVFA/T7-spoIVB* *S378A* was amplified from pSO247 using primers SO-P284 and SO-P285. Vector pSO40 was amplified using primers SO-P80 and SO-P283. Fragment was joined to pSO40 using GA. | This study |
| pSO252 | Km^R^; T7-Pro-σ^K^(1-127)-His_6_/T7-cytTM-SpoIVFB-FLAG_2_-His_6_/T7-BofA/SpoIVFA/SpoIVB | Fragment *bofA/spoIVFA/spoIVB* was amplified from pSO248 usng primers SO-P310 and SO-P285. Vector pSO213 was amplified using primers SO-P80 and SO-P311. Fragment was joined to pSO213 using GA. | This study |
| pSO253 | Km^R^; T7-Pro-σ^K^(1-127)-His_6_/T7-cytTM-SpoIVFB-FLAG_2_-His_6_/T7-BofA/SpoIVFA/SpoIVB S378A | Fragment *bofA/spoIVFA/spoIVB* *S378A* was amplified from pSO249 usng primers SO-P310 and SO-P285. Vector pSO213 was amplified using primers SO-P80 and SO-P311. Fragment was joined to pSO213 using GA. | This study |
| pSO254 | Km^R^; T7-Pro-σ^K^(1-127)-His_6_/T7-cytTM-SpoIVFB-FLAG_2_-His_6_/T7-BofA/SpoIVFA/T7-SpoIVB S378A | Fragment *bofA/spoIVFA/T7-spoIVB* *S378A* was amplified from pSO250 using primers SO-P310 and SO-P285. Vector pSO213 was amplified using primers SO-P80 and SO-P311. Fragment was joined to pSO213 using GA. | This study |
| pSO255 | Km^R^; T7-Cys-less Pro-σ^K^(1-127)-His_6_/T7-single-Cys A231C cytTM-SpoIVFB E44Q-FLAG_2_-His_6_ | pSO96 was subjected to SDM using primers SO-P184 and SO-P185, substituting A231C in cytTM-SpoIVFB. | This study |
| pSO256 | Km^R^; T7-Pro-σ^K^(1-127)-His_6_/T7-cytTM-SpoIVFB A32C C35S-FLAG_2_-His_6_ | pYZ2 was subjected to SDM using primers SO-P232 and SO-P233, substituting A32C and C35S in cytTM-SpoIVFB. | This study |
| pSO257 | Km^R^; T7-Pro-σ^K^(1-127)-His_6_/T7-cytTM-SpoIVFB L33C C35S-FLAG_2_-His_6_ | pYZ2 was subjected to SDM using primers SO-P234 and SO-P235, substituting L33C and C35S in cytTM-SpoIVFB. | This study |
| pSO258 | Km^R^; T7-Pro-σ^K^(1-127)-His_6_/T7-cytTM-SpoIVFB V86C-FLAG_2_-His_6_ | pYZ2 was subjected to SDM using primers SO-P238 and SO-P239, substituting V86C in cytTM-SpoIVFB. | This study |
| pSO259 | Km^R^; T7-Pro-σ^K^(1-127)-His_6_/T7-cytTM-SpoIVFB Q181C-FLAG_2_-His_6_ | pYZ2 was subjected to SDM using primers SO-P236 and SO-P237, substituting Q181C in cytTM-SpoIVFB. | This study |
| pSO260 | Km^R^; T7-Pro-σ^K^(1-127)-His_6_/T7-cytTM-SpoIVFB-FLAG_2_-His_6_/T7-GFPΔ27BofA G40C/SpoIVFA | pSO40 was subjected to SDM using primers SO-P226 and SO-P227, substituting G40C in GFPΔ27BofA. | This study |
| pSO261 | Km^R^; T7-Pro-σ^K^(1-127)-His_6_/T7-cytTM-SpoIVFB-FLAG_2_-His_6_/T7-GFPΔ27BofA A41C/SpoIVFA | pSO40 was subjected to SDM using primers SO-P228 and SO-P229, substituting A41C in GFPΔ27BofA. | This study |
| pSO262 | Km^R^; T7-Pro-σ^K^(1-127)-His_6_/T7-cytTM-SpoIVFB-FLAG_2_-His_6_/T7-GFPΔ27BofA I56C/SpoIVFA | pSO40 was subjected to SDM using primers SO-P222 and SO-P223, substituting I56C in GFPΔ27BofA. | This study |
| pSO263 | Km^R^; T7-Pro-σ^K^(1-127)-His_6_/T7-cytTM-SpoIVFB-FLAG_2_-His_6_/T7-GFPΔ27BofA H57C/SpoIVFA | pSO40 was subjected to SDM using primers SO-P224 and SO-P225, substituting H57C in GFPΔ27BofA. | This study |
| pSO289 | Km^R^; T7-Pro-σ^K^-His_6_/T7-cytTM- SpoIVFB-FLAG_2_/T7-GFPΔ27BofA/SpoIVFA | Fragment *spoIVFB* was amplified from pSO40 using primers SO-P138 and SO-P139. Vector pSO211 was amplified using primers SO-P140 and SO-P141. Fragment was joined to pSO211 using GA. | This study |
| pSO290 | Km^R^; T7-Pro-σ^K^-His_6_/T7-cytTM- SpoIVFB-FLAG_2_ | Vector pSO289 was amplified using primers SO-P319 and SO-P320, deleting *T7-gfpΔ27bofA/spoIVFA*. Vector ends were joined using GA. | This study |
| pSO312 | Km^R^; T7-Pro-σ^K^(1-127)-His_6_/T7-cytTM- SpoIVFB-FLAG_2_/T7-BofA | Fragment *bofA* was amplified from pSO212 using primers SO-P99 and SO-P100. Fragment was joined to pSO64 digested with NotI and NheI using GA. | This study |
| pSO313 | Km^R^; T7-Pro-σ^K^-His_6_/T7-cytTM- SpoIVFB-FLAG_2_/T7-GFPΔ27BofA | Fragment *gfpΔ27bofA* was amplified from pYZ46 using primers SO-P99 and SO-P100. Fragment was joined to pSO289 digested with NotI and NheI using GA. | This study |
| pSO314 | Km^R^; T7-Pro-σ^K^-His_6_/T7-cytTM- SpoIVFB-FLAG_2_/T7-BofA | Fragment *bofA* was amplified from pSO212 using primers SO-P99 and SO-P100. Fragment was joined to pSO289 digested with NotI and NheI using GA. | This study |
| pSO315 | Km^R^; T7-Pro-σ^K^-His_6_/T7-cytTM- SpoIVFB-FLAG_2_/T7-SpoIVFA | Fragment *spoIVFA* was amplified from pSO212 using primers SO-P82 and SO-P101. Fragment was joined to pSO289 digested with NotI and NheI using GA. | This study |
| pYZ2 | Km^R^; T7-Pro-σ^K^(1-127)-His_6_/T7-cytTM-SpoIVFB-FLAG_2_-His_6_ | pZR209 was digested with BglII and XhoI, and fragment *T7-cytTM-SpoIVFB-FLAG_2_-His_6_* was ligated with BglII-XhoI-digested pZR27. | Zhang, Y. unpublished |
| pYZ28 | Km^R^; T7-single-Cys P135C cytTM-SpoIVFB E44Q-FLAG_2_-His_6_ | pPL29 was digested with XbaI and XhoI, and fragment *T7-single-Cys P135C cytTM-SpoIVFB E44Q-FLAG_2_-His_6_* was ligated with XbaI-XhoI-digested pZR8. | Zhang, Y. unpublished |
| pYZ40 | Ap^R^; T7-single-Cys E44C cytTM-SpoIVFB-FLAG_2_-His_6_, which also has C35S C165L C167L C172S C246S |  | (3) |
| pYZ46 | Ap^R^; T7-GFPΔ27BofA/SpoIVFA | pZR33 was digested with BamHI and NotI, and fragment *spoIVFA* was ligated with BamHI-NotI-digested pZR62 | Zhang, Y. unpublished |
| pYZ77 | Ap^R^; T7-single-Cys V70C cytTM-SpoIVFB E44Q-FLAG_2_-His_6_, which also has C35S C165L C167L C172S C246S |  | (3) |
| pYZ112 | Ap^R^; T7-His_6_-MBP-FtsL(23-117) |  | (4) |
| pZR8 | Km^R^; T7-Pro-σ^K^(1-109)-His_6_ |  | (5) |
| pZR27 | Km^R^; T7-Pro-σ^K^(1-127)-His_6_ |  | (5) |
| pZR33 | Ap^R^; T7-His_10_-SpoIVFB-GFP/SpoIVFA |  | (5) |
| pZR53 | Ap^R^; T7-SpoIVB-His_6_ |  | (6) |
| pZR62 | Ap^R^; T7-GFPΔ27BofA |  | (5) |
| pZR209 | Ap^R^; T7-cytTM-SpoIVFB-FLAG_2_-His_6_ |  | (7) |

^a^Site-directed mutagenesis using the QuikChange kit (Stratagene).

^b^Gibson assembly (8).

^c^Overlap extension polymerase chain reaction.

**Primers used in this study**

| **Primer** | **Sequence** |
| --- | --- |
| SO-P11 | GTTAGTCGTCATTAAGCAATAAGATCCGAAGGAGATATAC |
| SO-P12 | GTATATCTCCTTCGGATCTTATTGCTTAATGACGACTAAC |
| SO-P13 | GAACTATACAAACTCGAGTTTGTGGCAGGTGCTTTG |
| SO-P14 | CAAAGCACCTGCCACAAACTCGAGTTTGTATAGTTC |
| SO-P48 | CTGTTAAATTTGTGGCAGCTGCTTTGCTGCTGGTTTG |
| SO-P49 | CAAACCAGCAGCAAAGCAGCTGCCACAAATTTAACAG |
| SO-P50 | GTGGCAGGTGCTTTGCTGGCGGTTTGTGTAAATATGTT |
| SO-P51 | AACATATTTACACAAACCGCCAGCAAAGCACCTGCCAC |
| SO-P52 | TTGCTGCTGGTTTGTGTAGCTATGTTTGGCGGCAGTCT |
| SO-P53 | AGACTGCCGCCAAACATAGCTACACAAACCAGCAGCAA |
| SO-P54 | GTTTGTGTAAATATGTTTGCCGGCAGTCTTGGCATTCA |
| SO-P55 | TGAATGCCAAGACTGCCGGCAAACATATTTACACAAAC |
| SO-P56 | CTTGGCATTCATGTGCCGGCTAATCTGGTTACAACAGC |
| SO-P57 | GCTGTTGTAACCAGATTAGCCGGCACATGAATGCCAAG |
| SO-P58 | GGCATTCATGTGCCGATTGCTCTGGTTACAACAGCTATC |
| SO-P59 | GATAGCTGTTGTAACCAGAGCAATCGGCACATGAATGCC |
| SO-P60 | GTGCCGATTAATCTGGTTGCAACAGCTATCAGCGGAA |
| SO-P61 | TTCCGCTGATAGCTGTTGCAACCAGATTAATCGGCAC |
| SO-P62 | GTTACAACAGCTATCAGCGCAATTTTAGGAATACCCGG |
| SO-P63 | CCGGGTATTCCTAAAATTGCGCTGATAGCTGTTGTAAC |
| SO-P64 | ACAGCTATCAGCGGAATTGCAGGAATACCCGGAATAGC |
| SO-P65 | GCTATTCCGGGTATTCCTGCAATTCCGCTGATAGCTGT |
| SO-P66 | CTATCAGCGGAATTTTAGCAATACCCGGAATAGCTGC |
| SO-P67 | CTATCAGCGGAATTTTAGCAATACCCGGAATAGCTGC |
| SO-P68 | CAGCGGAATTTTAGGAATAGCCGGAATAGCTGCGTTAG |
| SO-P69 | CTAACGCAGCTATTCCGGCTATTCCTAAAATTCCGCTG |
| SO-P70 | GAATTTTAGGAATACCCGCAATAGCTGCGTTAGTCGTC |
| SO-P71 | GACGACTAACGCAGCTATTGCGGGTATTCCTAAAATTC |
| SO-P72 | ATACCCGGAATAGCTGCGGCAGTCGTCATTAAGCAATT |
| SO-P73 | AATTGCTTAATGACGACTGCCGCAGCTATTCCGGGTAT |
| SO-P74 | GTCGTCATTAAGCAATTTGCCATTTAAGGATCCGAAGG |
| SO-P75 | CCTTCGGATCCTTAAATGGCAAATTGCTTAATGACGAC |
| SO-P76 | GTCATTAAGCAATTTATCGCTTAAGGATCCGAAGGAGA |
| SO-P77 | TCTCCTTCGGATCCTTAAGCGATAAATTGCTTAATGAC |
| SO-P80 | TGAGATCCGGCTGCTAACAAAGCCC |
| SO-P82 | GGCTTTGTTAGCAGCCGGATCTCAGCGGCCGCTTATTCAAATGAAATC |
| SO-P83 | CTTTAAGAAGGAGATATACATATGGCTAGCATGA |
| SO-P89 | CTTTAAGAAGGAGATATACATATGGCTAGCATGA |
| SO-P90 | ATGACAAGCTCGAGCACCACCACCACCACCACTGAGATCTCGATCCCGCGAAATTAATA |
| SO-P91 | GGCGGCAGTCTTGGCATTGCTGTGCCGATTAATCTGGT |
| SO-P92 | ACCAGATTAATCGGCACAGCAATGCCAAGACTGCCGCC |
| SO-P93 | CAGTCTTGGCATTCATGTGGCGATTAATCTGGTTACAAC |
| SO-P94 | GTTGTAACCAGATTAATCGCCACATGAATGCCAAGACTG |
| SO-P95 | ATAGCTGCGTTAGTCGTCGCTAAGCAATTTATCATTTA |
| SO-P96 | TAAATGATAAATTGCTTAGCGACGACTAACGCAGCTAT |
| SO-P97 | GTTAGTCGTCATTAAGCAAGCTATCATTTAAGGATCCGAAG |
| SO-P98 | CTTCGGATCCTTAAATGATAGCTTGCTTAATGACGACTAAC |
| SO-P99 | TAAGAAGGAGATATACATATGGCTAGC |
| SO-P100 | GGCTTTGTTAGCAGCCGGATCTCAGCGGCCGCGGATCCTTAAATGATAAATTGCTTAATG |
| SO-P101 | CTTTAAGAAGGAGATATACATATGGCTAGCATGAGTCACAGAGCAGATGAAATCAG |
| SO-P106 | GTCGTCATTAAGCAAGCTGCCGCTTAAGGATCCGAAGGAGA |
| SO-P107 | TCTCCTTCGGATCCTTAAGCGGCAGCTTGCTTAATGACGAC |
| SO-P108 | GAACTATACAAACTCGAGGGCGGCAGCGGCGGCAGCGGCGGCAGCTTTGTGGCAGGTGCTTTG |
| SO-P109 | CAAAGCACCTGCCACAAAGCTGCCGCCGCTGCCGCCGCTGCCGCCCTCGAGTTTGTATAGTTC |
| SO-P112 | CTTACCCTTAAATTTATTAGCACTACTGGAAAACTAC |
| SO-P113 | GTAGTTTTCCAGTAGTGCTAATAAATTTAAGGGTAAG |
| SO-P114 | TTCGCGTATGGTCTTCAAAGCTTTGCGAGATACCCAG |
| SO-P115 | CTGGGTATCTCGCAAAGCTTTGAAGACCATACGCGAA |
| SO-P116 | GATTCAATTATCCTGAAATTACTTCTGTCGGCCTTACTTGTTCTCGTTTCAGCT |
| SO-P117 | AGCTGAAACGAGAACAAGTAAGGCCGACAGAAGTAATTTCAGGATAATTGAATC |
| SO-P118 | TGAGATCCGGCTGCTAACA |
| SO-P119 | CTTGTCATCGTCATCCTTGTAATCC |
| SO-P120 | GGATTACAAGGATGACGATGACAAGTAATGAGATCTCGATCCCGCGAAATTAATA |
| SO-P132 | GCATGAATTCTTTTCTTCGCAAGAAAACATTAAGAAG |
| SO-P133 | GCATGCATGCGCTTGTTCTAGTACAAGCTTATGAG |
| SO-P134 | GACTAGAGTCGAATCCCGGCATGCACATAAGGAGGAACTACT |
| SO-P135 | AGTAGTTCCTCCTTATGTGCATGCCGGGATTCGACTCTAGTC |
| SO-P136 | GCGACGTATGCAGCGAGGAGTATTGAAAATGAAATCC |
| SO-P137 | GGATTTCATTTTCAATACTCCTCGCTGCATACGTCGC |
| SO-P138 | ATGAATAAATGGCTCGACCTTATCTTAAAG |
| SO-P139 | GTAGGGCAGAAGCAGTTCC |
| SO-P140 | CTTCCATGGAGGAACTGCTTC |
| SO-P141 | GATAAGGTCGAGCCATTTATTCATGG |
| SO-P142 | GCATGCACATAAGGAGGAACTACTATGAGTAAAGGAGAAGAACTTTTC |
| SO-P143 | CGACCGGCGCTCAGGATCCTTAAATGATAAATTGCTT |
| SO-P148 | CGCATTGAAAAAAACAAAAAAATAAGATCCATGAGTAAAGGAGAAGATCT |
| SO-P149 | AGATCTTCTCCTTTACTCATGGATCTTATTTTTTTGTTTTTTTCAATGCG |
| SO-P152 | GGTGCTTTGCTGCTGGTTAGTGTAAATATGTTTGGCG |
| SO-P153 | CGCCAAACATATTTACACTAACCAGCAGCAAAGCACC |
| SO-P156 | GAGATATACATATGGCTAGCATGGAAACTGAAGAAGGTAAACTGGTAATCTGGATTAACG |
| SO-P157 | CTGATGCCAATCCACTTCTCGAGGGCCGCCGCGTCTTT |
| SO-P158 | CTCGAGAAGTGGATTGGC |
| SO-P159 | CATGCTAGCCATATGTATATCTCC |
| SO-P162 | GAAGAAAAAAAATACTTATGCCTCATGGCTAAAGGGGATG |
| SO-P163 | CATCCCCTTTAGCCATGAGGCATAAGTATTTTTTTTCTTC |
| SO-P164 | AGAAAAAAAATACTTAGAGTGCATGGCTAAAGGGGATGA |
| SO-P165 | TCATCCCCTTTAGCCATGCACTCTAAGTATTTTTTTTCT |
| SO-P168 | GTGAGATTTCTCCTCGAATGCTATTACGGAAAAAACAGG |
| SO-P169 | CCTGTTTTTTCCGTAATAGCATTCGAGGAGAAATCTCAC |
| SO-P170 | GATTTCTCCTCGAAAGGTGCTACGGAAAAAACAGGGAG |
| SO-P171 | CTCCCTGTTTTTTCCGTAGCACCTTTCGAGGAGAAATC |
| SO-P172 | TTCTCCTCGAAAGGTATTGCGGAAAAAACAGGGAGC |
| SO-P173 | GCTCCCTGTTTTTTCCGCAATACCTTTCGAGGAGAA |
| SO-P174 | AAGGAATTGAAAGCTATTGCGCTGGAAAAGGGACAAAG |
| SO-P175 | CTTTGTCCCTTTTCCAGCGCAATAGCTTTCAATTCCTT |
| SO-P176 | GGAATTGAAAGCTATTCCTGTGGAAAAGGGACAAAGCTG |
| SO-P177 | CAGCTTTGTCCCTTTTCCACAGGAATAGCTTTCAATTCC |
| SO-P180 | GAAACTTCTGCCGCTGACATGCAAGGCGGAGGATAAAGTC |
| SO-P181 | GACTTTATCCTCCGCCTTGCATGTCAGCGGCAGAAGTTTC |
| SO-P182 | CTTCTGCCGCTGACAGTATGCGCGGAGGATAAAGTCTATC |
| SO-P183 | GATAGACTTTATCCTCCGCGCATACTGTCAGCGGCAGAAG |
| SO-P184 | CTGCCGCTGACAGTAAAGTGCGAGGATAAAGTCTATCATG |
| SO-P185 | CATGATAGACTTTATCCTCGCACTTTACTGTCAGCGGCAG |
| SO-P186 | GCGAAGAAAAAAAATACTGCGAGCTCATGGCTAAAGGG |
| SO-P187 | CCCTTTAGCCATGAGCTCGCAGTATTTTTTTTCTTCGC |
| SO-P188 | GAGAAACTTCTGCCGCTGTGCGTAAAGGCGGAGGATAAAG |
| SO-P189 | CTTTATCCTCCGCCTTTACGCACAGCGGCAGAAGTTTCTC |
| SO-P196 | GGAATTGAAAGCTATTCCGCTTGCAAAGGGACAAAGCTGG |
| SO-P197 | CCAGCTTTGTCCCTTTGCAAGCGGAATAGCTTTCAATTCC |
| SO-P202 | GGCTTGCTCACAGGCCATTGCAAAGCATTATTATGTCTG |
| SO-P203 | CAGACATAATAATGCTTTGCAATGGCCTGTGAGCAAGCC |
| SO-P204 | CATTCATGTGCCGATTAATTGCGTTACAACAGCTATCAGC |
| SO-P205 | GCTGATAGCTGTTGTAACGCAATTAATCGGCACATGAATG |
| SO-P206 | CATGTGCCGATTAATCTGTGTACAACAGCTATCAGCGGA |
| SO-P207 | TCCGCTGATAGCTGTTGTACACAGATTAATCGGCACATG |
| SO-P209 | GGCTTGCTCACAGGCCATTGCAAAGCATTATTATCTCTG |
| SO-P210 | CAGAGATAATAATGCTTTGCAATGGCCTGTGAGCAAGCC |
| SO-P211 | GCTTCTGCCCTACAAGCTTTAATGAGATCTCGATCCC |
| SO-P212 | GGGATCGAGATCTCATTAAAGCTTGTAGGGCAGAAGC |
| SO-P218 | GAGCTTGTCTTTTTAGTATGTTACGTGAAAAACAATGC |
| SO-P219 | GCATTGTTTTTCACGTAACATACTAAAAAGACAAGCTC |
| SO-P222 | GTTTGGCGGCAGTCTTGGCTGTCATGTGCCGATTAATCTG |
| SO-P223 | CAGATTAATCGGCACATGACAGCCAAGACTGCCGCCAAAC |
| SO-P224 | GGCGGCAGTCTTGGCATTTGTGTGCCGATTAATCTGGT |
| SO-P225 | ACCAGATTAATCGGCACACAAATGCCAAGACTGCCGCC |
| SO-P226 | GCTGTTAAATTTGTGGCATGTGCTTTGCTGCTGGTTAG |
| SO-P227 | CTAACCAGCAGCAAAGCACATGCCACAAATTTAACAGC |
| SO-P228 | GTTAAATTTGTGGCAGGTTGTTTGCTGCTGGTTAGTGT |
| SO-P229 | ACACTAACCAGCAGCAAACAACCTGCCACAAATTTAAC |
| SO-P232 | CTCACAGGCCATATGAAATGTTTATTATCTCTGCTCCTG |
| SO-P233 | CAGGAGCAGAGATAATAAACATTTCATATGGCCTGTGAG |
| SO-P234 | CAGGCCATATGAAAGCATGTTTATCTCTGCTCCTGATTG |
| SO-P235 | CAATCAGGAGCAGAGATAAACATGCTTTCATATGGCCTG |
| SO-P236 | TTATTCGTGATTCCTCTGTGTATCAGCGCATGGGTTTTG |
| SO-P237 | CAAAACCCATGCGCTGATACACAGAGGAATCACGAATAA |
| SO-P238 | GTTAAAGGAAGAGTTTGCGTGCATTATTGCCGGACCTCT |
| SO-P239 | AGAGGTCCGGCAATAATGCACGCAAACTCTTCCTTTAAC |
| SO-P242 | CGTGTTTTTTTGCTGCCGGCTGGCGGAACGGTCGAAGTG |
| SO-P243 | CACTTCGACCGTTCCGCCAGCCGGCAGCAAAAAAACACG |
| SO-P244 | GGCATCGTACAGGGGATGGCCGGAAGCCCGATCATTCAA |
| SO-P245 | TTGAATGATCGGGCTTCCGGCCATCCCCTGTACGATGCC |
| SO-P248 | GATTTCATTTGAATAAGCGGCCGCGTCCGGCGTAGAGGATCG |
| SO-P251 | TTGCTGCTGGTTTGTGTAGATATGTTTGGCGGCAGTC |
| SO-P252 | GACTGCCGCCAAACATATCTACACAAACCAGCAGCAA |
| SO-P257 | AAAGGATCCCACCACCAC |
| SO-P260 | GATCTTAGTGGTGGTGGTGGTGGTGGGATCCTTTCCCCTTCGCCTTCTTC |
| SO-P269 | CAATGCGCGCAAATGCATG |
| SO-P270 | TGAAATCCTCATGCATTTGCGC |
| SO-P271 | CTTAAGGGAAGCGAAGCAGC |
| SO-P272 | CCGCTACTGTCAACAACTTTTTAGC |
| SO-P273 | TAAGGATCCGAAGGAGATATACATATGAG |
| SO-P276 | GACTCATATGTATATCTCCTTCGGATCCTTAAATGATAAATTGCTTAATGACGACTAACG |
| SO-P277 | TTAAGAAGGAGATATACATATGGCTAGCATGGAGCCTATTTTTATTATTGGGATTATTTT |
| SO-P280 | CATGCTAGCCATATGTATATCTCCTTCTTAAAG |
| SO-P282 | CTCAGTGGTGGTGGTGGTGGTGCTCGAGGCTTGCTTTTTCTTTTCCATAAATATCGATTC |
| SO-P283 | CTGCTAGTTGAACGCTTCCATCTTCAATGTTG |
| SO-P284 | CAACATTGAAGATGGAAGCGTTCAACTAGCAG |
| SO-P285 | GGCTTTGTTAGCAGCCGGATCTCAGTG |
| SO-P286 | CTAATCGTTTTCTGATTTCATCTGCTCTGTG |
| SO-P287 | ATGAGTCACAGAGCAGATGAAATCAGAAAACGATTAG |
| SO-P288 | ATCTCATTACTTGTCATCGTCATCCTTGTAATCCTTGTCATCGTCATCCTTGTAATC |
| SO-P289 | CTTGCTTTCCTGGCTCCTGAA |
| SO-P290 | CGATCCTCTACGCCGGACGCGGCCGCTTATTCAAATGAAATC |
| SO-P293 | GCAAATCAGCGCATGGGTTTTGTTTGTCTTTCTGG |
| SO-P294 | GCAAATCAGCGCATGGGTTTTG |
| SO-P295 | CAAACAAAACCCATGCGCTGATTTGC |
| SO-P302 | GATTTCATTTGAATAAGCGGCCGCTGAAGGAGATATACATATGCCCGATAAC |
| SO-P303 | ATGTTATCGGGCATATGTATATCTCCTTCAGCGGCCGCTTATTCAAATG |
| SO-P304 | CTTGCTTTCCTGGCTCCTGAACACAAAAATAAGGAACAGC |
| SO-P305 | CTCAGTGGTGGTGGTGGTGGTGCTC |
| SO-P306 | GAGCACCACCACCACCACCACTG |
| SO-P307 | CTGTTCCTTATTTTTGTGTTCAGGAGCCAGGAAAGCAAG |
| SO-P310 | GCTGTTAAATTTGTGGCAGGTGCTTTGCTG |
| SO-P311 | CAGCAAAGCACCTGCCACAAATTTAACAGC |
| SO-P319 | CGATGACAAGTAATGAGATCCGGCTGCTAACAAAGCCC |
| SO-P320 | GATCTCATTACTTGTCATCGTCATCCTTGTAATCCTTGTCATCGTCATCCTTGTAATCAA |
| LK2465 | AAAGAGCTTGTCTGTTTAGTATCTTAC |
| LK2466 | GTAAGATACTAAACAGACAAGCTCTTT |
| LK2467 | CTTGTCTTTTTATGTTCTTACGTGAAA |
| LK2468 | TTTCACGTAAGAACATAAAAAGACAAC |
| LK2473 | GTATCTTACGTGTGTAACAATGCCTTT |
| LK2474 | AAAGGCATTGTTACACACGTAAGATAC |
| LK2691 | GCAGCATGCCCCAGCTGATGAATCAATACAATC |
| YZ1 | GATTGTATTGATTCATTGCCTGGGGCATGCTGCTC |
| YZ2 | GAGCAGCATGCCCCAGGCAATGAATCAATACAATC |
| YZ11 | GATTGTATTGATTCATCAGCTGGGGCATGCTGC |

**Sequencing primers used in this study**

| **Primer** | **Sequence** | **Notes** |
| --- | --- | --- |
| SO-P23 | CATTACCTGTCCACACAATCTGC | Forward primer. Binds in GFP. |
| SO-P84 | CATACCCACGCCGAAACAAG | Forward primer. Binds upstream of the T7 promoter in pET29b. |
| SO-P86 | GAGTAAAGGAGAAGATCTCGATCC | Forward primer. Binds in Pro-σ^K^. |
| SO-P87 | CTCCAGTGAAAAGTTCTTCTCC | Reverse primer. Binds in GFP. |
| SO-P104 | CATGGAGGAACTGCTTCTGC | Forward primer. Binds in SpoIVFB. |
| SO-P121 | GCAGATTGTGTGGACAGGTAATG | Reverse primer. Binds in GFP. |
| SO-P122 | CTGTCAGTGGAGAGGGTGA | Forward primer. Binds in GFP. |
| SO-P123 | TCACCCTCTCCACTGACAG | Reverse primer. Binds in GFP. |
| SO-P157 | CTGATGCCAATCCACTTCTCGAGGGCCGCCGCGTCTTT | Reverse primer. Binds in MBP. |
| SO-P160 | CTTCAGCGAGACCGTTATAGC | Reverse primer. Binds in MBP. |
| SO-P161 | ACTGACGGGTCCAATGTTTG | Reverse primer. Binds in SpoIVFA. |
| SO-P276 | GACTCATATGTATATCTCCTTCGGATCCTTAAATGATAAATTGCTTAATGACGACTAACG | Reverse primer. Binds in BofA |
| SO-P279 | GACTGACGGTTAAGGTGCAG | Forward primer. Binds in SpoIVFA |
| DP18 | GCTAGTTATTGCTCAGCGG | Reverse primer. Binds T7 terminator |
| DP89 | TAATACGACTCACTATAGGG | Forward primer. Binds T7 promoter. |

***B. subtilis* strains used in this study**

| **Strain** | **Genotype** | **Construction** | **Citation** |
| --- | --- | --- | --- |
| PY79 | Prototrophic wild-type strain |  | (9) |
| BK754 | *spoIVB165* |  | (10) |
| ZR264 | *spoIVB165 bofA*::*erm* |  | (5) |
| SO3 | *spoIVB165 bofA*::*erm amyE*::P*_bofA_-gfp*Δ*27bofA* | ZR264 was transformed with pSO78 | This study |
| SO6 | *spoIVB165 bofA*::*erm amyE*::P*_bofA_-gfp*Δ*27bofA N48A* | ZR264 was transformed with pSO86 | This study |
| SO8 | *spoIVB165 bofA*::*erm amyE*::P*_bofA_-gfp*Δ*27bofA N61A* | ZR264 was transformed with pSO87 | This study |
| SO10 | *spoIVB165 bofA*::*erm amyE*::P*_bofA_-gfp*Δ*27bofA T64A* | ZR264 was transformed with pSO88 | This study |

**References**

1.	Rudner DZ, Losick R (2002) A sporulation membrane protein tethers the pro-σ^K^ processing enzyme to its inhibitor and dictates its subcellular localization. *Genes Dev.* 16:1007-1018.

2.	Green D, Cutting S (2000) Membrane topology of the *Bacillus subtilis* Pro-σ^K^ processing complex. *J. Bacteriol.* 182:278-285.

3.	Zhang Y, Luethy PM, Zhou R, Kroos L (2013) Residues in conserved loops of intramembrane metalloprotease SpoIVFB interact with residues near the cleavage site in Pro-σ^K^. *J. Bacteriol.* 195:4936-4946.

4.	Parrell D, Zhang Y, Olenic S, Kroos L (2017) *Bacillus subtilis* intramembrane protease RasP activity in *Escherichia coli* and *in vitro*. *J. Bacteriol.* 199:e00381-00317.

5.	Zhou R, Kroos L (2004) BofA protein inhibits intramembrane proteolysis of pro-σ^K^ in an intercompartmental signaling pathway during *Bacillus subtilis* sporulation. *Proc. Natl. Acad. Sci. USA* 101:6385-6390.

6.	Zhou R, Kroos L (2005) Serine proteases from two cell types target different components of a complex that governs regulated intramembrane proteolysis of pro-σ^K^ during *Bacillus subtilis* development. *Mol. Microbiol.* 58:835-846.

7.	Zhou R, Cusumano C, Sui D, Garavito RM, Kroos L (2009) Intramembrane proteolytic cleavage of a membrane-tethered transcription factor by a metalloprotease depends on ATP. *Proc. Natl. Acad. Sci. USA* 106:16174-16179.

8.	Gibson DG*, et al.* (2009) Enzymatic assembly of DNA molecules up to several hundred kilobases. *Nat. Methods* 6:343-345.

9.	Youngman P, Perkins JB, Losick R (1984) Construction of a cloning site near one end of Tn*917* into which foreign DNA may be inserted without affecting transposition in *Bacillus subtilis* or expression of the transposon-borne *erm* gene. *Plasmid* 12:1-9.

10.	Cutting S, Driks A, Schmidt R, Kunkel B, Losick R (1991) Forespore-specific transcription of a gene in the signal transduction pathway that governs pro-σ^K^ processing in *Bacillus subtilis*. *Genes Dev.* 5:456-466.
